# Conformationally responsive dyes enable protein-adaptive differential scanning fluorimetry

**DOI:** 10.1101/2023.01.23.525251

**Authors:** Taiasean Wu, Joshua C. Yu, Arundhati Suresh, Zachary J. Gale-Day, Matthew G. Alteen, Amanda S. Woo, Zoe Millbern, Oleta T. Johnson, Emma C. Carroll, Carrie L. Partch, Denis Fourches, Nelson R. Vinueza, David J. Vocadlo, Jason E. Gestwicki

## Abstract

Flexible *in vitro* methods alter the course of biological discoveries. Differential Scanning Fluorimetry (DSF) is a particularly versatile technique which reports protein thermal unfolding via fluorogenic dye. However, applications of DSF are limited by widespread protein incompatibilities with the available DSF dyes. Here, we enable DSF applications for 66 of 70 tested proteins (94%) including 10 from the SARS-CoV2 virus using a chemically diverse dye library, Aurora, to identify compatible dye-protein pairs in high throughput. We find that this protein-adaptive DSF platform (paDSF) not only triples the previous protein compatibility, but also fundamentally extends the processes observable by DSF, including interdomain allostery in O-GlcNAc Transferase (OGT). paDSF enables routine measurement of protein stability, dynamics, and ligand binding.

**One-Sentence Summary:** Next generation protein-adaptive DSF (paDSF) enables rapid and general measurements of protein stability and dynamics.

## Main Text

A major challenge in biochemistry is the lack of flexible methods to monitor protein biochemical states *in vitro*. Among the emerging solutions (*1*, *2*), techniques using protein apparent thermal stability as a proxy for biochemical state (*3*–*7*) excel in simplicity and generality. These strengths are especially true of Differential Scanning Fluorimetry (DSF), which is unique in its use of an environmentally sensitive dye to report unfolding (*7*). This dye, typically SYPRO Orange or ANS, reports unfolding by fluorescing brightly in response to its selective interaction with a protein’s unfolded or partially unfolded states (*7*, *8*) (Figs. 1A and B).

**Fig. 1.**
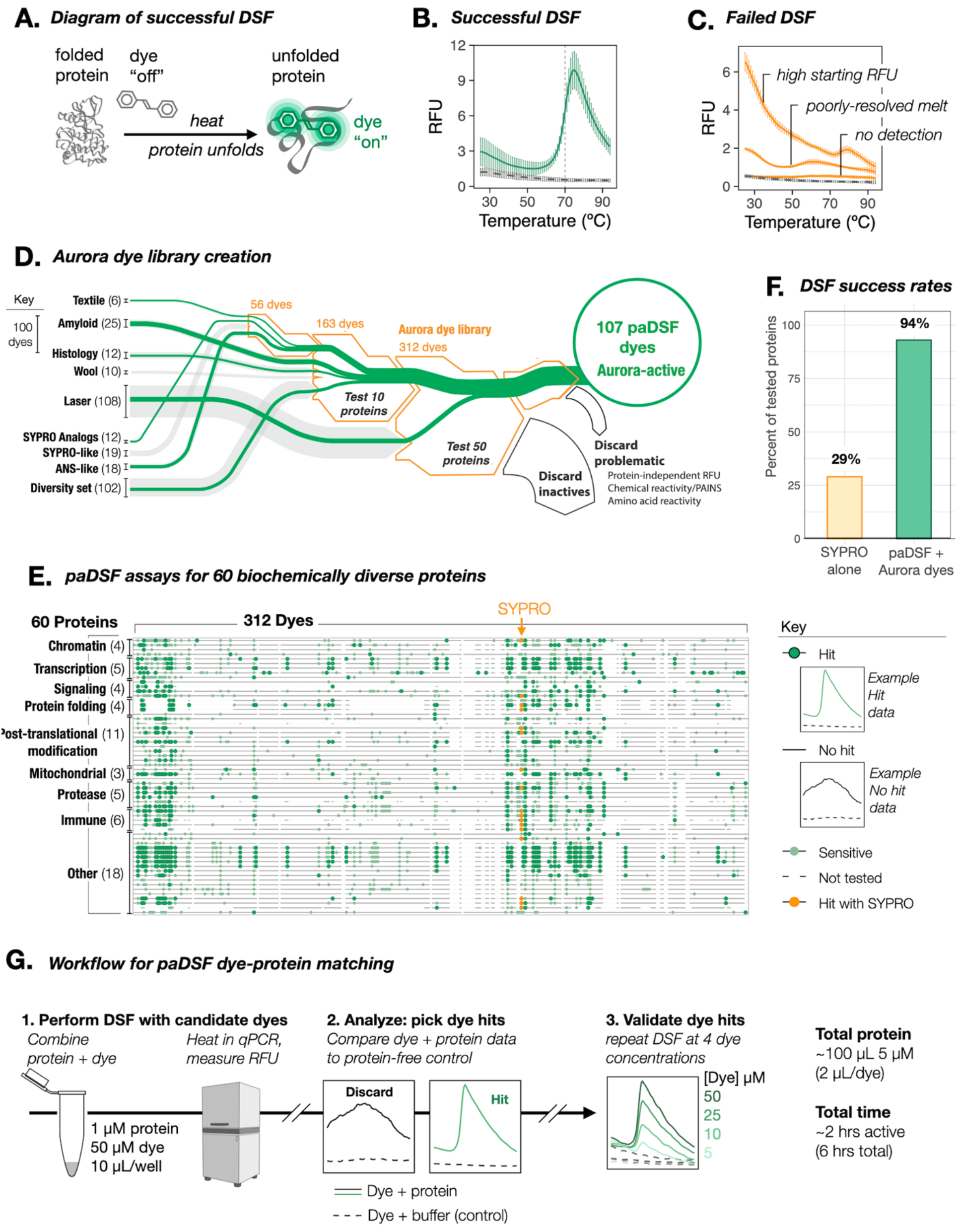
Resolution of DSF incompatibility with a systematically curated dye library. (**A**) Schematic of successful DSF. Dye interacts selectively with thermally denatured states of the protein, and fluoresces brightly in response. (**B**) Example raw DSF data when protein and SYPRO Orange are compatible (data shown for lysozyme; solid green, buffer and dye alone: dashed grey), and (**C**) incompatible (with proteins Bag2, BSA, and Hsp10: solid grey, buffer and dye alone: dashed grey). (**D**) Sankey diagram summarizing the creation and assembly process for the Aurora dye library. (Line widths: number of dyes in a group; green line widths: number of novel DSF dyes within a group). Counter screens include: folding-independent protein reactivity (free amino acids, linear peptides, and HA, FLAG, 6xHis epitopes) and practical incompatibilities (0.01% Triton X-100, EDTA, BME, DTT, TCEP; HEPES, Tris, PBS, MOPS). (**E**) Summary of the paDSF dye-protein pairs identified for the 60 tested proteins. Individual horizontal lines are individual screened proteins. Individual columns are individual dyes. (**F**) Fraction of tested proteins compatible with DSF before and after introduction of paDSF and Aurora dyes. (**G**) Schematic of streamlined workflow to pair proteins with paDSF dyes. See fig. S3 and Movie S1.

DSF is widely used at both exploratory and industrial scales, reporting relevant biochemical or biophysical changes via associated shifts in protein apparent melting temperature (T_ma_). Common applications include the observation of many non-enzymatic processes, including the binding of ligands, peptides, and co-factors, as well as characterizing the biophysical impacts of mutations or buffer conditions (*9*, *10*). DSF is used with biochemically diverse proteins and across multiple disciplines (*2*, *10*), including structural biology (*11*, *12*), small molecule discovery (*9*, *10*), protein design (*13*), and therapeutic formulation (*14*). DSF fills a crucial niche as an agile and low-risk assay in the technological ecosystem of these fields.

Yet DSF frequently fails in practice, reporting unfolding inaccurately (*10*, *15*) (Fig. 1C). We hypothesized that these failures were not random, but instead, examples of proteins for which the fidelity of the DSF dye to unfolded states broke down. This premise could collapse widespread DSF incompatibilities into the potentially solvable task of identifying a diversified set of DSF dyes. Bespoke DSF dyes reported for specific proteins (*14*, *16*) demonstrated that alternative dye-protein pairings were possible. Given a platform to streamline protein-dye pairing, DSF compatibility could theoretically be achieved on demand. We term this approach and the resulting assays “protein-adaptive DSF” (paDSF).

Here, we report the successful creation of paDSF assays for over 94% (66 of 70) of tested proteins, via a library of paDSF dyes and rapid dye-protein pairing protocol. The key development in paDSF was the creation and systematic testing of an initial library of 312 chemically diverse dyes, which we term the Aurora library (Fig. 1D, table S1 and data S1). The paDSF assays described here overcome DSF incompatibilities of high-priority targets from diverse biological processes, including gene regulation, inflammation, development, and SARS-CoV2 infection. On average, proteins paired to 13 dyes. By selecting among hit dyes for the multidomain enzyme O-GlcNAc transferase (OGT), paDSF could be used to monitor its interdomain allostery. This application demonstrates how the advances made by paDSF extend beyond protein compatibility, facilitating previously challenging biochemical measurements. paDSF and the Aurora dyes may extend technological access to both established and emerging research questions.

## Results

### Identification of paDSF dyes

To create the Aurora library (table S1, data S1 and S4), dyes were drawn heavily from established fields (*17*) which, like DSF, rely on dye-protein interactions or environmentally sensitive fluorescence (Fig. 1D). This included textile, histology, amyloid (*18*), film, and laser dyes, as well as a subset of the Eastman-Kodak Max Weaver Dye Library (*19*). To systematically assess candidate dyes for paDSF activity, we assembled a panel of 60 purified proteins. In addition to including diverse cellular processes, these proteins included individual high-priority targets, such as hormone receptors, caspases, and transcription factors. The panel was also varied in biochemical properties such as oligomeric state, secondary structure composition, extent of intrinsic disorder, function, and size (table S2 and data S3).

The 312 candidate dyes in Aurora were then assessed by performing DSF on each protein with each dye (Fig. 1G). Preliminary activity was defined as the production of a sigmoidal thermal transition in the presence of protein, with negligible fluorescence in a protein-free, buffer-matched control (Fig. 1E, fig. S1 and data S2). Dyes were considered paDSF active “hits” only if they also passed the 20 counter screens for problematic properties, including protein-independent fluorescence, amino acid detection and buffer incompatibilities (Fig. 1D and Fig. 1D legend, fig. S1 and data S2).

Excitingly, the final dye library achieved paDSF activity for 56 of the 60 tested proteins, amounting to 94% protein compatibility, a three-fold improvement over SYPRO alone (29%, Figs. 1E and F). Each protein matched to an average of 13 dyes (lower quartile = 4 dyes, median = 9.5, upper-quartile = 21), with the lowest hit rates typically observed for very small (< 15 kDa) proteins. Of the 15,658 protein-dye pairs assessed here, 781 were hits, amounting to a 5% hit rate overall. In some cases, proteins were not screened against the full Aurora library, so this success rate is likely an underestimate. The T_ma_s calculated from these screens were consistent with benchmark biophysical measurements and SYPRO-based experiments where available (fig. S1 and S2). paDSF results were highly reproducible and demonstrated dye concentration dose-dependence in validation experiments (fig. S1 and data S2). paDSF dyes also overcame reported incompatibilities of SYPRO with buffer additives including detergent and EDTA (*9*, *14*) (data S2). Conditions and raw data for the discovered paDSF assays for all 60 proteins can be explored online (https://padsfdyes.shinyapps.io/Exp1243_heatmap_cache/).

From the Aurora library, we defined the subset of 107 dyes with paDSF activity for at least one protein as Aurora-active. To streamline future efforts, we defined another subset of dyes, Aurora-concise, which contains the 48-highest performing, readily-accessible dyes (table S1). Aurora-concise could be an especially powerful tool to rapidly identify paDSF dyes for most proteins. A detailed guide for library assembly and screening, including a video protocol, is provided in the supplement (fig. S3, movie S1). A typical dye screen requires 100 μL of 5 μM protein, uses no specialized instrumentation, and can be completed within a day (Fig. 1G).

### Rapid paDSF assay development for SARS-CoV2 proteins

As a real-world test of paDSF and Aurora-concise, we then attempted to create paDSF assays for 10 proteins from the SARS-CoV2 virus (Fig. 2A). For each protein, multiple dye-protein pairs were identified, and screening was completed in approximately three hours (Fig. 2B and C, data S1). With the resulting paDSF assays, thermal shifts were observed in response to known ligands for PLPro (*20*) and Nsp3 macrodomain-1 (nsp3-mac1) (*21*), consistent with accurate monitoring of their binding (fig. S4).

**Fig. 2.**
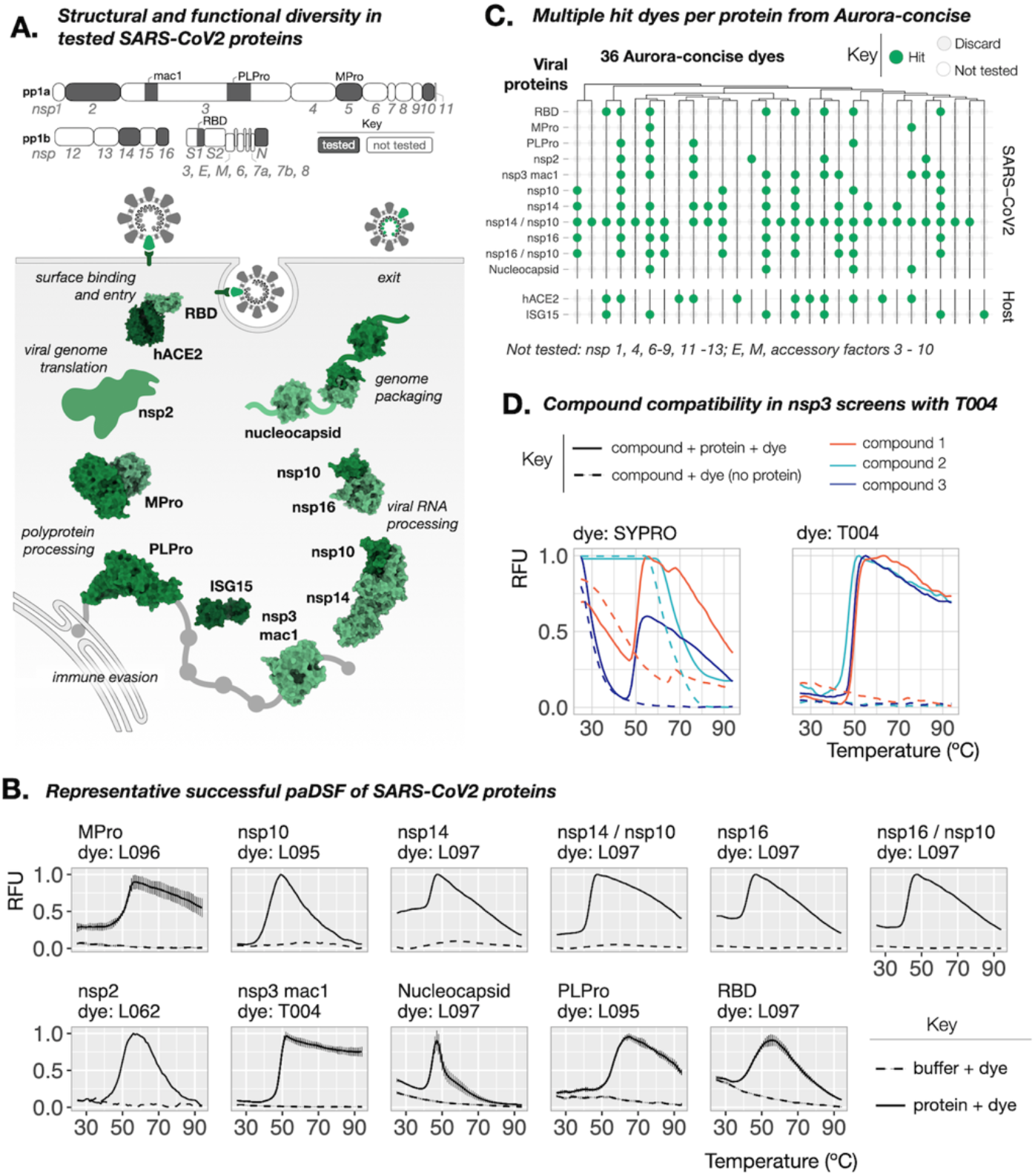
Streamlined creation of paDSF assays for SARS-CoV2 proteins. (**A**) Schematic of the SARS-CoV2 proteins tested here. Top: full SARS-CoV2 transcriptome, and the proteins tested. Bottom: schematic of the tested proteins, demonstrating variation in structure, stage in the viral life cycle, and currently understood functions (PDBs: 5EMZ, 6XAA, 6M3M, 6WZO, 7JYC, 6MOJ, 1Z2M, 6YWL, 6W4H, 5C8U). (**B**) Representative raw data from successful paDSF for all tested proteins from Aurora-concise dyes. Shown dyes have pre-existing commercial sources. See supplement for full results. (**C**) Summary “Willow plot” of paDSF hit for all tested viral proteins and host proteins hACE2 and ISG15, highlighting the presence of multiple hit dyes for each protein. Individual rows represent individual proteins. Individual columns represent individual dyes. Data shown for 36-dye subset of Aurora-concise already accessible from existing commercial sources. Dyes are arranged by hierarchical clustering based on chemical similarity. (**D**) Comparison of raw DSF data collected in an nsp3-mac1 compound screen, demonstrating the resolution of DSF failures due to artifactual SYPRO Orange activation by compounds (left) by use of dye T004 (right).

The thermal shifts of PLPro and nsp3-mac1 could be monitored using the multiple hit dyes interchangeably (fig. S4), imparting chemical flexibility in downstream applications. A recent report describes substantial technical barriers in the use of DSF to monitor small molecule binding to nsp3-mac1 (*22*). We performed a pilot small molecule screen against nsp3-mac1 using SYPRO and found that compound-induced SYPRO fluorescence rendered results uninterpretable for 27 of 320 compounds (8%) (Fig. 2D, fig. S5). This incompatibility was reduced to only 2% when SYPRO was replaced with the paDSF dye T004 (Fig. 2D and fig. S5). This compatibility persisted in a subsequent medicinal chemistry campaign, where T004-based paDSF enabled assessment of 189 of 192 (98.5%) lead-like molecules, assisting in the identification of a high-affinity inhibitor of nsp3-mac1 (*23*). Beyond overcoming compound incompatibility, having multiple dyes for each protein also enables measurement in different wavelengths, from blue to near infra-red (ex/em = 470/520 nm to 660/705 nm) (data S2). This spectral diversity could help avoid compound interference and enable broad instrument compatibility.

The paDSF assays reported here could support wide-ranging research on each of these 10 SARS-Cov-2 proteins, including structural characterization and ligand discovery. These proteins play diverse roles in SARS-CoV2 replication and pathogenesis, including entry, translation, proteolysis, RNA binding and genome packing (Fig. 2A; bottom) (*24*). They also comprise many properties which typically complicate standard techniques, including disorder (*25*, *26*), phase separation (*27*, *28*), protein-protein interactions (*29*, *30*), and protein nucleic-acid-interactions (*24*). The rapid identification of paDSF assays for all proteins, regardless of their function and biochemical diversity, corroborates the high success rates seen during paDSF development (see Fig. 1F). Broadly, these dye-based optimizations also demonstrated that paDSF can expand technological access not only to new proteins, but also new applications.

### Extension to interdomain allostery

Despite its complicated basis in molecular thermodynamics (*31*–*33*), T_ma_ is well established as a reasonable and sensitive proxy for a wide variety of biochemical perturbations. We wondered if dye selection could optimize paDSF to monitor biochemical phenomena typically beyond standard DSF applications, such as interdomain allostery. As a test, we used O-GlcNAc transferase (OGT). OGT is a critical conserved enzyme that installs an O-linked N-acetylglucosamine (O-GlcNAc) sugar unit to serine and threonine residues of target proteins (*34*).

OGT comprises a catalytic domain and a client-binding tetratricopeptide repeat (TPR) domain. Dye screening yielded nine hits for OGT from Aurora-concise (data S1) which collectively reported two distinct T_ma_s in the measured temperature range (Figs. 3A and B, fig. S6). We term these transitions T_ma OGT-1_ and T_ma OGT-2_ (T_ma OGT-1_ = 46.55 ± 0.94 °C, T_ma OGT-2_ = 58.97 ± 0.96 °C) (fig. S6). paDSF of the isolated catalytic and TPR domains reported single transitions which matched T_ma_OGT-1_ and T_ma_OGT-2_, respectively (Δ[T_ma cat_ vs T_ma_OGT-1_] = 0.34 °C, p = 0.4; Δ[T_ma TPR_ vs T_ma_OGT-2_] = 1.1 °C, p = 0.02) (Fig. 3B, figs. S6 and S7). Importantly, depending on the dye used, the composition of the raw fluorescence curve for OGT ranged from predominantly first transition (~2:1 RFU_OGT-1_: RFU_OGT-2_) to predominantly second (~1:4 RFU_OGT-1_: RFU_OGT-2_) (Fig. 3A, figs. S6 and S7).

**Fig. 3.**
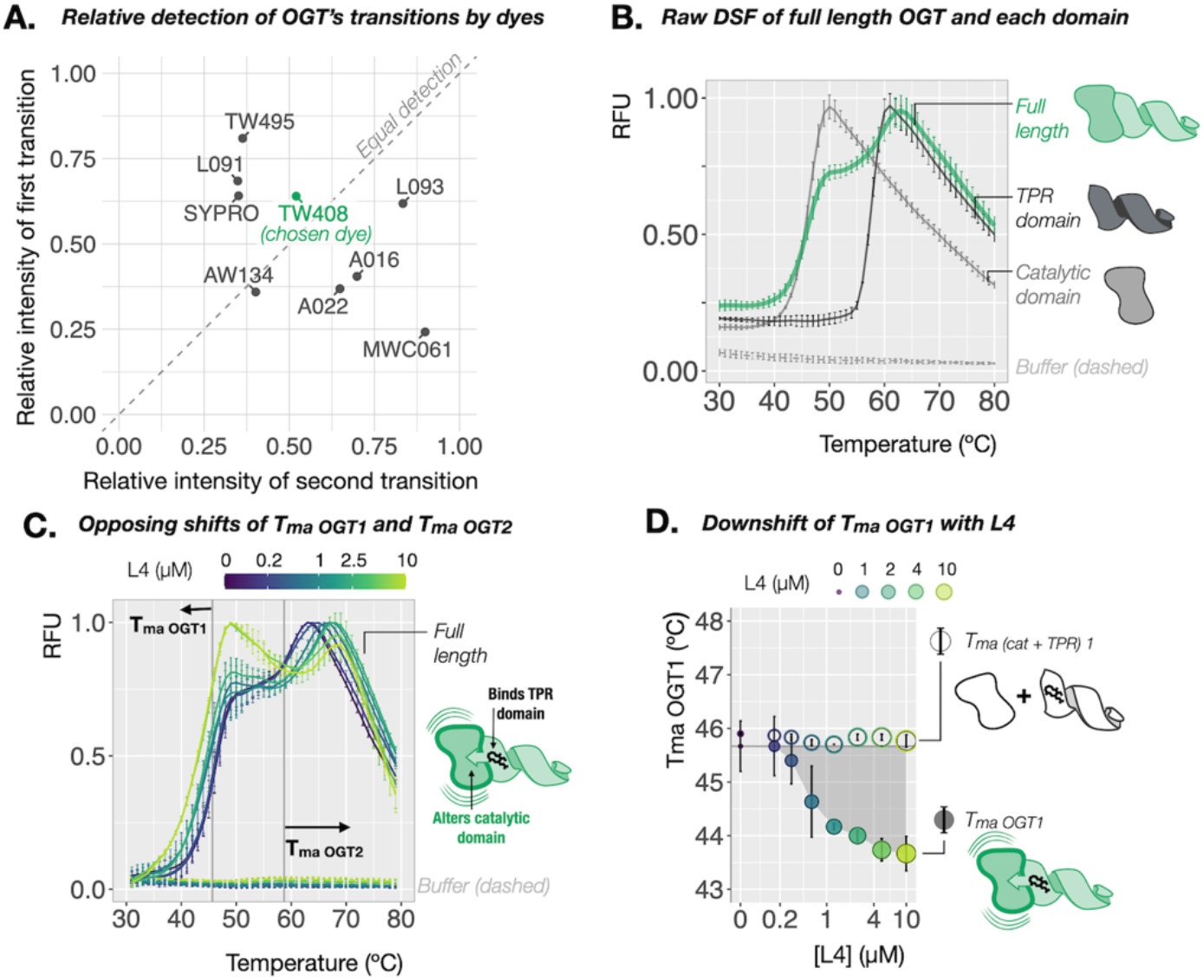
paDSF enables observation of allostery in OGT. (**A**) Dye-dependent variation in the relative detection of the two consecutive transitions observed in paDSF of OGT. Raw paDSF with dye TW408 for full length OGT, (**B**) compared to its isolated catalytic or TPR domains. Full length OGT undergoes two apparent transitions. The isolated catalytic and TPR domains undergo single transitions which align with the first and second transitions of OGT, respectively, (**C**) with increasing concentrations of its high affinity allosteric ligand, L4, demonstrating coupled and opposing thermal shifts of the first and second transitions of OGT. Grey vertical reference lines at T_ma OGT1_ and T_ma OGT2_ in apo protein. (**D**) Effect of L4 titration on T_ma OGT1_, compared to a 1:1 mixture of isolated catalytic and TPR domains. A dose-responsive decrease in the first transition is observed only with full length OGT, demonstrating that physical connection between the domains is required for coupled thermal shifts of the two transitions.

Interested in monitoring the potential crosstalk between these transitions, we selected dye TW408 for further studies because it gave the most equal detection between OGT’s two transitions (1.2:1 RFU_OGT-1_: RFU_OGT-2_) (Fig. 3A) as well as between the isolated domains (1.2:1 RFU_cat_:RFU_TPR_) (fig. S8). Titration of OGT’s glycosyl donor UDP-GlcNAc (*22*) induced a modest selective thermal upshift of T_ma_OGT-1_ but not T_ma_OGT-2_ (ΔT_ma_OGT-1_= 0.58 ± 0.05 °C, p = 1e-10; ΔT_ma_OGT-2_ = 0.20 ± 0.42, p = 0.66) (fig S8). This was mirrored in the isolated domains, where UDP-GlcNAc induced a comparable upshift in the catalytic domain, but not the TPR (ΔT_ma cat_ = 0.69 ± 0.14, p = 1.5e-12; ΔT_ma TPR_ = −0.15 ± 0.22, p = 0.98) (fig. S8). UDP-GlcNAc binds the active site of OGT’s catalytic domain, and this result provides a benchmark for single-domain ligand binding in TW408 paDSF.

OGT was recently found to exhibit interdomain allostery upon binding of the novel macrocyclic ligand L4 (*35*). L4 binds the TPR domain with nanomolar affinity and appears to induce a corresponding conformational change in the catalytic domain. Excitingly, in contrast to the single-transition upshifts observed for UDP-GlcNAc, titration of L4 induced simultaneous but opposing thermal shifts in T_ma_OGT-1_ and T_ma_OGT-2_ (ΔT_ma_OGT-2_ = 6.07 ± 1.02, p = 2.3e-12; ΔT_ma_OGT-1_ = −2 ± 0.79, p = 6.2e-11) (Fig. 3D). Consistent with a possible allosteric mechanism, this coupling occurred only with full-length OGT. L4 titration with the individual domains, alone or in a 1:1 mixture, induced an upshift in only the TPR-associated transition (Fig. 3D and fig. S9) (ΔT_ma_TPR_ = 5.5 ± 0.2, p = 3.6e-14; ΔT_ma_cat_ = −0.1 ± 0.4, p = 0.2, ΔT_ma2 cat + TPR_ = 5.77 ± 0.22, p = 4.7e-15, ΔT_ma1 cat + TPR_ = −0.13 ± 0.12, p = 0.09). This coupling of thermal shifts could be a readout of interdomain allostery in OGT.

To further test this model, we first confirmed that L4 did not impact the oligomerization state of OGT (fig. S10), ruling this out as the source of the behavior. Glycosylation of a model peptide substrate by OGT (KKKYPGGSTPVSSANMM) exhibited a similar L4-induced heat intolerance at temperatures near T_ma_OGT-1_, but well below T_ma_OGT-2_, providing coarse functional corroboration for the observed downshift of T_ma_OGT-1_ (41 °C; Fig. S11). Overall, this result demonstrates how the ability to optimize simultaneous detection of multiple transitions via choice of dye expanded the biochemical processes of OGT accessible using paDSF.

### Chemical trends in dye-protein recognition

paDSF dyes enable the first consideration of DSF dyes as a nascent class of small molecule probes. Chemoinformatic analysis revealed that paDSF activity occurs in chemically diverse dyes (Fig. 4A and fig. S12) and is enriched in specific chemical scaffolds from the full 312-dye Aurora library (Fig. 4A). Performance is not correlated with hydrophobicity (fig. S13), with hydrophilic dyes like MWC007 and MWC061 among the highest performing (Fig. 4B). This suggests that paDSF dyes can, and do, rely on chemical/structural features beyond simple hydrophobic targeting to selectively detect unfolded states of proteins.

**Fig. 4.**
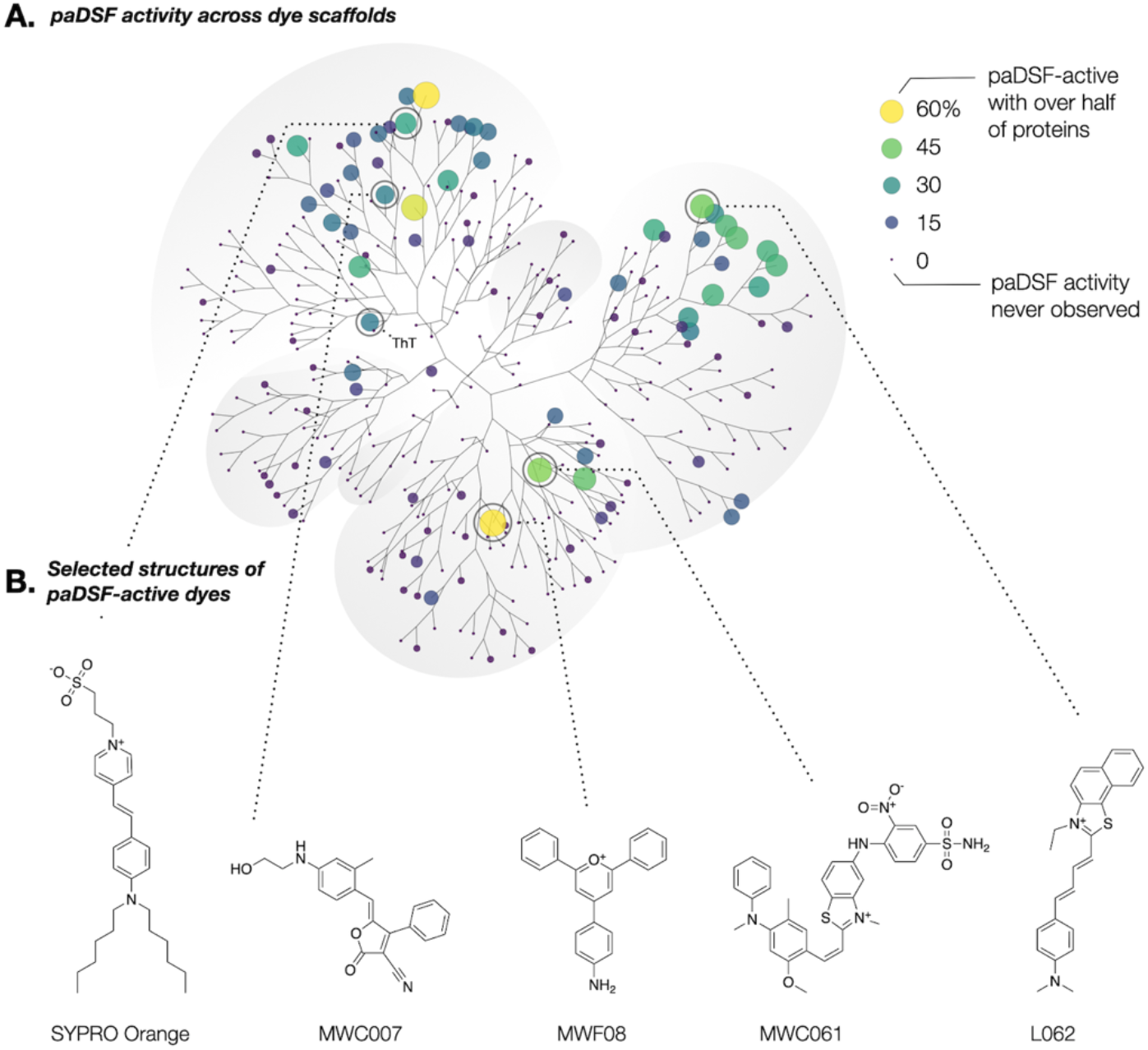
Emergence of structural trends within paDSF dyes. (**A**) Annotated polar dendrogram of structural relationships between the 312 dyes in Aurora, as determined by pairwise ECFP4 Tanimoto coefficient followed by hierarchical clustering. Each point (“node”) represents a single dye. Point size and point color represent the percentage of the 70 tested proteins (60 protein panel + 10 proteins from SARS-CoV2) for which that dye was a paDSF hit. Larger, lighter points indicate a more general dye; smaller, darker points indicate more selective dyes. (**B**) Chemical structures of selected high-performing paDSF dyes from Aurora.

Known fluorogenic motifs (polymethine, merocyanine) occurred in many DSF-active dyes, though not all dyes with known fluorogenic motifs produced paDSF activity (coumarin, rhodamine) (fig. S14 and table S3). Intriguingly, structure-based literature searches of some high-performing paDSF active dyes such as MWF08 and T004 returned little or no precedent for environmental sensitivity, suggesting that the Aurora dyes may contain unstudied fluorogenic motifs (Fig. 4B). Overall, the breadth of paDSF-active chemical space implies that DSF activity can proceed via multiple distinct fluorogenic mechanisms.

The biochemical features detected by each dye remain unknown. However, the unique protein compatibilities of each dye imply that different dyes use different chemical features to distinguish native from non-native states. The protein selectivities of paDSF dyes ranged from detection of a single protein out of the 70 tested to 60% of this same protein set (Fig. 4A and fig. S15). No two proteins paired to an identical set of hit dyes (Fig. 1E, fig. S16), providing a unique “fingerprint” of paDSF dyes responsive to each protein. A typical fingerprint included both promiscuous and selective paDSF dyes (fig. S17), suggesting that a range of common and unique features may be detected in each unfolded protein. Proteins with greater sequence similarity were more likely to produce similar dye fingerprints, represented as a weak but significant correlation between pairwise distances in protein Multiple Sequence Alignments and the Jaccard index of dye fingerprints (fig. S18). Further characterization of paDSF active dyes may illuminate the biochemical features which drive their fluorescence responsiveness, which may in turn guide creation of new paDSF compatible dyes.

## Discussion

Since its introduction in the 1980s, DSF has been limited to the applications possible with SYPRO Orange and ANS alone. paDSF reframes the dye as a key variable in DSF experimental design, enabling dye selection to expand, rather than limit, uses of the technology as a whole. In addition to achieving widespread protein compatibility, this work revealed less obvious, but equally impactful, ways in which dye selection enables new applications. For example, paDSF offers improved resilience to common practical pitfalls, shown here in the resolution of compound incompatibility by exchanging SYPRO Orange for dye T004 in small molecule screens of nsp3-mac1.

Other aspects of dye selection suggest that paDSF may represent a more fundamental evolution in DSF technology. In particular, we find that different dyes emphasize different components of complex unfolding trajectories, and that this variation allows previously cryptic biochemical phenomena to be monitored using paDSF depending on the dye employed. This dye-driven tuning is demonstrated here in the use of dye TW408 to robustly detect two distinct transitions of OGT. The resulting TW408-based paDSF was able to discern both orthosteric and allosteric ligand binding, based on the independence or coupling of the extracted thermal shifts. Intriguingly, the observation of multiple reproducible transitions was not unique to OGT but emerged broadly across the 70 tested proteins. Dye-dependent expansion of the processes observable using paDSF may therefore be possible for proteins beyond OGT.

paDSF, alongside other developing technologies, is poised to meet outstanding needs for flexible and straightforward biochemical assays. Because paDSF and the Aurora library expand the flexibility of DSF while retaining its practical simplicity, paDSF may be particularly useful where practical accessibility or high throughput are required. Advances in the remaining biophysical and optical variables in paDSF, such as denaturation protocols, optical measurements, and data analysis approaches, could synergize with dye selection to extend possible paDSF applications even further. paDSF can help address the growing diversity of processes known to govern protein activities, both immediate and unexplored.

## Acknowledgments

Specific material contributions are listed in the Supporting Information. The authors thank Matt Jacobson, James Fraser, Kevan Shokat, Douglas Wassarman, Daniel Schwarz, Daniel Elnatan, Hao Shao, Megan Moore and Sarah Williams for helpful discussions. Proteins were generously donated by Andrew Ambrose, Aye Thwin, Douglas Wassarman, Erin Thompson, Alfred Freeberg, Galen Correy, Hao Shao, Hayden Saunders, Jeffery Swan, Maria Janowska, Marina Ramirz-Alvarado, Matt Ravalin, Meghna Gupta, Mericka McCabe, Michael Schoof, Rebecca Freilich, and Victor Long from the labs of Michelle Arkin, Ana Maria Cuervo, James Fraser, Jason Gestwicki, John Gross, Rachel Klevit, Geeta Narlikar, Kevan Shokat, Daniel Southworth, Robert Stroud, Klim Verba, Peter Walter, and Jordan Ward. Several SARS-CoV2 proteins were provided by the UCSF QBI Coronavirus Research Group (QCRG). We are grateful for additional support from Mike Capps and Zare Karakesisoglu (Analytik Jena); Carrie Brown and David Schubach (Exciton-Luxottica); Duncan Beniston (ChemBridge Corporation); Kathleen Kitto, the Home Away students; Stefan Gahbauer, Seth Vigneron, Jon Lee, D’Anne Duncan, and Naledi Saul (UCSF).

## Funding

National Institute of Health U19 (JEG, AI171110)

Sandler Program for Breakthrough Biomedical Research (JEG)

Tau Consortium (JEG)

National Science Foundation Graduate Research Fellowship (TW, 1000259744)

National Institute of Standards and Technology (NRV, 60NANB19D115)

National Institutes of Health K99 (OTJ, K99NS128717)

National Institutes of Health F32 (ECC, F32AG076281)

National Institutes of Health R01 (JEG, NRV, ZM, 1R01GM141299-01)

National Institutes of Health R01 (CLP, GM107069, R01GM121507, and R35GM141859)

GlycoNet, the Canadian Glycomics Network (DJV, CD-1)

Canadian Cancer Society Research Institute (DJV, CCSRI-706825)

Canada Research Chairs program (DJV, Tier I CRC in Chemical Biology)

## Author contributions

Conceptualization: TW, JEG, MGA, DJV

Data curation: TW

Formal analysis: TW, ZJD

Funding acquisition: TW, JEG

Investigation: TW, JCY, AS, MGA, ZM, AW, CLP, NRV, DF

Methodology: TW, JCY, AS

Visualization: TW

Project administration: JEG

Supervision: JEG

Writing – original draft: TW, EC, OTJ, ZJD, JEG

Writing – review & editing: TW, MGA, DJV, JEG

## Competing interests

JEG is a consultant for Contour and DiCE and a consultant and shareholder for Protego and Kaizen. All other authors declare that they have no competing interests.

## Data and materials availability

All data are available in the main text or the supplementary materials. A subset of the Aurora library is only available through a material transfer agreement (MTA) with North Carolina State University.

## Notes

https://padsfdyes.shinyapps.io/Exp1243_heatmap_cache/

### Summary of Updates

Misspelled an authors name in the online form (it was correct in the PDF).

