## Supporting Information for "Conformationally responsive dyes enable protein-adaptive differential scanning fluorimetry"

### Supplementary Materials for
#### Conformationally responsive dyes enable protein-adaptive differential scanning fluorimetry

Taiasean Wu, Joshua C. Yu, Arundhati Suresh, Zachary J. Gale-Day, Matthew G. Alteen, Amanda S. Woo, Zoe Millbern, Oleta T. Johnson, Emma C. Carroll, Carrie L. Partch, Denis Fourches, Nelson R. Vinueza, David J. Vocadlo, Jason E. Gestwicki*

**This PDF file includes:**

Materials and Methods
Figs. S1 to S18
Tables S1 to S3
Caption for Movie S1
Captions for Data S1 to S4

**Other Supplementary Materials for this manuscript include the following:**

Movie S1
Data S1 to S4 (Dye library sources and extended details, Full dye screen results for all proteins, Protein panel sources and extended details, Dye screen results for all proteins.)

**Materials and Methods**

Making and storing dye stock solutions

All commercial dyes were used without further purification. The purity of all Max Weaver dyes (*19*) was confirmed >80% by LCMS at North Carolina State University prior to addition to the dye library. All LCMS analyses of the Max Weaver library dyes were performed on an Agilent Technologies 1260 High-Performance Liquid Chromatography (HPLC) system coupled with an Agilent 6520 Q-TOF mass spectrometer. (Agilent Technologies, CA, USA). Ionization was performed via electrospray ionization with the following parameters: gas temperature 350 ºC, drying gas 6 liters per minute, nebulizer 35 psi, $V_{cap}$ voltage 3500 V and fragmentor voltage at 160 V. Powders were stored in RT, sealed and parafilmed, in the dark. From each dye, a 5 mM DMSO was made by transferring ~1 mg of solid dye into a pre-massed 1.5 mL microcentrifuge tube using a fresh microspatula (disposable smartSpatula(R),L anti-static, Cat# Z561762-300EA). The tube was massed again to obtain the exact mass, and the solid dye was resuspended in enough DMSO to create a 5 mM stock solution. To dissolve, the solution was vortexed for 30 seconds, left in the dark at room temperature for 16 hours, and vortexed again. Final dissolved solution was distributed into 15 uL aliquots in PCR strip tubes (0.2mL Split-Strip 8-Strip Tubes w/Individually Attached Flat Caps, Clear) using an E100 ClipTip p125 Matrix Pipette (Yellow). 5 mM stocks were stored in 1.5 mL microcentrifuge tubes, as well as in 15 µL aliquots in PCR strip tubes in the dark in -80 ºC. Fresh dye stocks were prepared for the entire library every 12-18 months. In addition, 40 µL of each dye was place in a LabCyte Echo Qualified 384PP Source Microplate (Cat# PP0200).

Dye library DMSO stock handling

To a LabCyte Echo Qualified 384PP Source Microplate (Cat# PP0200), 40 µL of each dye library dye was added to create a "mother plate" from which single-use dye screening plates "daughter plates" could be dispensed. At regular intervals, the mother plate was thawed, and 250 nL of each dye was transferred into a single well of a 384-well V-bottom PP Greiner BioOne Microplate (Cat #781280) using a LabCyte Echo. Transfer instructions were created using an in-house web app. The mother and daughter plates were sealed in foil (Excel Scientific PolarSeal Aluminum Microplate Seals, FPS-EK-100), and thawed prior to use.

General protein handling.

All proteins were stored and handled as recommended by each protein source. All proteins were screened in the preferred biochemical buffer for that specific protein, including any additives. Proteins with no preferred buffer were screened in 10 mM HEPES pH 7.20, 200 mM NaCl. See table S2 for information on specific protein sources, buffers, and screening concentrations.

Calculation of structural and biochemical protein properties.

Primary sequences were analyzed using the Bio.SeqUtils.ProtParam python module. The specific functions used for each parameter were: .molecular_weight() for **Molecular weight**, .gravy() for Grand average of **hydropathicity** index (GRAVY), .get_amino_acids_percent() for **primary sequence composition**, and .secondary_structure_fraction() for fraction of amino acids found primarily in **helix, turn, and sheet secondary structures**. Code for this analysis was sourced with minor adaptations from "Visualizing and Analyzing Proteins in Python" by Aren Carpenter published online (https://towardsdatascience.com/visualizing-and-analyzing-proteins-in-python-bd99521ccd).

The presence of **disordered regions** was calculated using the IUPred2A(*36*) online server (https://iupred.elte.hu/). A multi-FASTA file containing all protein primary sequences was uploaded, and two separate queries were submitted, (i) searching for long regions of disorder, using the settings “Analysis type: IUPred2A long disorder” (>30 residues) with “Context-dependent predictions (default ANCHOR2)”, and (ii) searching for short regions os disorder, using the settings “Analysis type: IUPred2A long disorder” (<30 residues) with “Context-dependent predictions (default ANCHOR2)”. A single “percent predicted disorder”, presented in table S3 was calculated for each protein by dividing the number of residues with a IUPred long disorder probability ≥0.5 by the total number of residues in that protein.

Calculation of protein sequence similarity

Primary sequences for all tested proteins (see data S2) were converted to FASTA format using basic string handling functions in R, and then read as sequences using the function msa::readAAStringSet(), and a multiple sequence alignment was calculated between all sequences using msa::msa(). The resulting multiple sequence alignment was converted to a seqinr-compatible class using msa::msConvert(type = “seqinr::alignment”). The resulting object was used to calculate pairwise distance matrices for both sequence similarity and sequence identity using seqinr::dist.alignment(matrix = “similarity”) and seqinr::dist.alignment(matrix = “identity”), respectively. The resulting root distance matrix was converted to a similarity matrix (S) by S = 1 - (root distance)^2.

Dye screens

Each protein was screened for DSF compatibility with library dyes in the following manner. (A generalizable protocol for dye screening is provided in fig. S4 and movie S1.) For individual protein sources, concentrations, and screening buffers, see table S2.

*Resuspension of library dyes* A single “daughter plate”, containing 250 nL of 5 mM DMSO stock of each library dye was thawed from -80 ºC. After thawing, the daughter plate was visually inspected to ensure no wells were missing dyes. 10 mL of the desired buffer was placed in a buffer reservoir (VWR Catalog #89094-670), and 20 uL of buffer was transferred into each well of the daughter plate using a E100 ClipTip p125 Matrix Pipette (Thermo Fisher #4671040BT, “Yellow”) and associated tips (ClipTip 384 125 Pipette Tips, #94410153) . The resuspended daughter plate was spun-down in a salad spinner for 30 seconds to remove bubbles, and placed on the deck of an Opentrons OT-2 liquid handling robot at room temperature.

*Protein dispensing and dye addition*. Protein was diluted to 5X screening concentration, and 2 µL of this solution was dispensed into each well of a white, low-volume 384-well qPCR plate (Axygen PCR-284-LC480WNFBC, lot # 23517000) using an E100 ClipTip matrix pipette, and the plate was spun down for 30 seconds to settle droplets to the bottoms of the wells. The plate was placed on the deck of the Opentrons OT-2 liquid handling robot, and the OT-2 was used to transfer 8 µL of resuspended dye from the resuspended dye daughter plate into the protein-containing qPCR plate. Any dyes which remained adhered to the daughter plate were not further mixed or disrupted prior to transferring. Following transfer, each well contained 10 µL of volume, with a typical protein concentration of 1 µM and typical dye concentration of 50 µM. Exact conditions for each screen can be found in table S2.

*Heating and measurement*. The qPCR plate was sealed with optically clear sealing film (Applied Biosystems MicroAmp Optical Adhesive Film #4311971) and spun for 30 seconds to settle droplets into the bottoms of wells and remove bubbles. The qPCR plate was placed in an Analytik Jena 384G qTower qPCR and heated from 25 ºC to 94 ºC in 69 increments of 1 ºC every 30 seconds with lid heating to 98 ºC, and fluorescence measurements were taken at each degree in six channels ("FAM" ex/em = 470/520, "JOE" ex/em = 515 nm/545 nm, "TAMRA" ex/em = 535 nm/580 nm, "ROX" ex/em = 565 nm/605 nm, "Cy5" ex/em = 630 nm/670 nm, and "Cy5.5" ex/em = 660 nm/705 nm). This heating and measuring procedure results in a heating rate of 1 ºC per 1 minute, and a total runtime of approximately 70 minutes. Raw fluorescence readings were exported and analyzed using an in-house script. Tmas were extracted using DSFworld (*15*).

*Hit validation*. Following hit selection, a custom dye daughter plate was created, including the initially-tested dye concentration, as well as three two-fold serial dilutions for each chosen hit dye. Validation daughter plates were dispensed as described above in "Dye library DMSO stock handling"; serial dilutions of dye stocks were performed during the LabCyte Echo transfer by dispensing the appropriate amounts of either dye stock or DMSO into the well. The above procedure for "Dye screens" was repeated using this validation plate, yielding a four-point dose-response for each dye. Raw fluorescence readings were exported and analyzed using an in-house script.

Dye screening of SARS-CoV2 proteins

Dye screens for all SARS-CoV2 proteins were conducted following the procedure described in Fig. 1G and fig. S3, and outlined above "Dye screens". See table S2 for protein sources. Proteins were screened in the following buffer, termed "CoV2 DSF buffer": 50 mM Tris HCl pH 7.5, 150 mM NaCl, 1 mM TCEP, 0.01% Triton X-100. For all experiments, TCEP and Triton were added immediately prior to use. All Tmas were calculated using DSFworld (*15*).

*paDSF of PLPro with GRL0617*

GRL0617 was obtained from TOCRIS (Cat # 7280) and resuspended in DMSO to a stock concentration of 50 mM. In 10 mL of CoV2 DSF buffwer (50 mM Tris HCl pH 7.5, 150 mM NaCl, 1 mM TCEP, 0.01% Triton X-100 a 2.5X GRL0617 stock was prepared by diluting 5 µL of 50 mM GRL0617 stock in 995 µL buffer. Three 2-fold serial dilutions were prepared from this stock. 80 µL of 5X PLPro (5 µM) was prepared in CoV2 DSF buffer from a 32 µM stock solution. 1 mL of 2.5X dye was prepared for each dye. These components were combined and mixed well to yield the final tested conditions: 5 µM dye; 100, 10, and 0 µM GRL0617; 1 µM PLPro, or for no-protein controls, buffer alone in place of protein. 10 µL of solution was transferred to each well of the 384-well plate, and DSF was performed as described in "*Heating and measurement"* above.

*paDSF of nsp3-mac1 with ADP ribose*

A daughter plate containing 250 nL of each dye was dispensed using the LabCyte Echo as described above in *"Dye library DMSO stock handling"*. ADP ribose was obtained from Sigma Aldrich (A0752) and resuspended in DMSO to a stock concentration of 10 mM. To the CoV2 DSF buffer, EDTA was added to a final concentration of 1 mM. In this buffer, seven serial dilutions of ADP ribose were prepared at 1.25X final concentration. The top concentration was prepared by adding 74 µL of 10 mM ADP ribose to CoV2 buffer. To the dye daughter plate, 20 µL of the appropriate concentration of ADP ribose was added to each well. A 5X protein solution was prepared by adding 13.3 µL of 300 µM purified nsp3-mac1 to 386.7 µL of buffer for a final concentration of 10 µM, and 2 µL of this protein solution was added to each well of a white qPCR plate. To the white qPCR palte, 8 µL of dye-ADPribose solution was added. DSF was performed as descrived above in "Heating and Measurement." All dyes were tested at a final concentration of 5 µM with the exception of L094, which was tested at 25 µM. ADP ribose was tested at 300, 150, 75, 37.5, 18.8, 9.3, 4.6 and 0 µM. Protein was tested at a final concentration of 2 µM, or 0 µM for the no-protein controls.

*paDSF of nsp3-mac1 with screening compounds*

To the CoV2 DSF buffer, EDTA was added to a final concentration of 1 mM, DMSO was added to a final concentration of 3%, and SYPRO Orange was added to a final concentration of 5X. In this buffer, a 2X protein solution of nsp3-mac1 was created by dilution of 270 µL of 37 µL nsp3-mac1 stock solution into a 2 mL final volume. Buffer and stock solution of a purine small molecule library was added to the DSF plate, for a final compound concentration of 20 µM. DSF was performed as described above in "heating and measurement." For head-to-head comparison between dye T004 and SYPRO, the above procedure was repeated, using either 5X SYPRO Orange or 5 µM T004, and 192 compounds from the ZINC catalog, as described in (*23*).

Mass photometry of OGT

*Slide cleaning and gasket preparation.* Prior to experimental set up, glass slides (High Precision Deckgläser Microscope cover glass, 24 x 50 mM, 170 +/- 5 µM, No 1.5H) were cleaned by three repetitions of: submersion and swirling in a beaker of isopropyl alcohol, air dry, submersion and swirling in a beaker of miliQ water, air dry, followed by one final round of drying by pressurized air passed through a pasteur pipette tip. Mass photometry gaskets (Grace BioLabs Ref 103250 Culture Well Reuseable Gaskets, CW-50R-1.0, 50-3mm DIA x 1 mm depth, 3-10 uL) were prepared by cutting into 2 x 4 well strips.

*Sample preparation.* 50 mL of filtered OGT buffer was prepared by combining 2.5 mL Tissue-Culture Grade 1M HEPES (Gibco 1 M HEPES Buffer Solution, Ref 15630-106) with 3 mL 5M NaCl and 44.5 uL MiliQ water, both of which had been spin filtered immediately prior to use (Ultrafree PVDF 0.1 µm, Ref R7KA45465). To a 4 mL aliquot of this buffer, 4 µL of DTT was added, and the buffer was spin-filtered twice. In this final buffer, each of full-length OGT, OGT TPR domain, and OGT catalytic domain were diluted to 2 µM, and the ligand L4 was diluted from a 10 mM DMSO stock to a final concentration of 20 µM. All protein and ligand solutions were once again spin filtered. From the 20 µM L4 solution, six two-fold serial dilutions were made in the filtered buffer. To each of three L4 concentrations or L4-free buffer, a 1:1 mixture was made between ligand and each of the protein solutions, or protein-free buffer, for final L4 concentrations of 0, 0.6, and 2.5 µM, and final protein concentrations of 0 or 1 µM. The solutions were left to incubate for 2 hours at room temperature to equilibrate.

*Data collection.* Mass photometry data was collected on a Refeyn OneMP Mass Photometer as follows. Instrument was focused and calibrated to BSA (2 mg/mL BSA solution, Bradford Assay Kit) and thyroglobulin (Cytiva MW Native Marker Kit, Thyroglobulin standard Mr 669000) mass standards. Each sample was measured well as follows: into a fresh gasket well, 10 uL of filtered buffer was pipetted, followed by 2 µL of 2 µM protein-ligand mixture, followed by gentle mixing by pipetting up and down, for a final volume of 12 µL. Final conditions measured were 0, 0.6, and 2.5 μM L4 in the presence of full length OGT, catalytic domain alone, TPR domain alone, or buffer (as a control for protein-independent ligand aggregation).

paDSF of OGT with dye TW408

*OGT buffer:* To 10 mL of OGT buffer, 10 uL of 1 M DTT was added. Following thorough mixing, 100 μL of a 5 mM DMSO stock of TW408 was added to this buffer, to yield the final OGT TW408 buffer (final buffer: 50 mM HEPES pH 7.4, 300 mM NaCl, 1 mM DTT, 50 μM TW408, 1% DMSO). All following solutions were prepared in this buffer.

*2X protein solutions:* 140 μL of 4X solutions of purified TPR and catalytic domains were prepared by dilution of concentrated protein stocks to a final 4X concentration of 4 μM. These 4X protein solutions were used to prepare 140 μL of 2X solutions of either (1) TPR alone (70 μL 4 μM TPR + 70 μL buffer), (2) catalytic alone (70 μL 4 μM catalytic + 70 μL buffer), or (3) 1:1 mix of catalytic and TPR domains (70 μL 4 μM TPR + 70 μL 4 μM catalytic). A 2X solution of OGT was then prepared by first creating a 1.3X concentrated TW408 buffer, to offset the volume displaced by the OGT stock solution, and then combining 210 μL of this buffer with 70 μL of 8 μM OGT stock.

*Compound solutions:* A concentrated solution of the highest tested concentration for each compound was created in OGT buffer. From this solution, six 2-fold serial dilutions were made in OGT buffer.

*UDP GlcNAc:* Ultra Pure UDP-GlcNAc (Promega Cat # V7071) was prepared in buffer in seven 2-fold serial dilutions, from an highest concentration of 100 μM.

*L4:* 1 uL of 50 mM DMSO stock of L4 was diluted in 9 uL of DMSO and 990 uL of TW408 OGT buffer, to yield a final 5X concentrated L4 solution of 50 μM. From this 50 μM solution, seven 2-fold serial dilutions were created in OGT buffer.

*Combination of protein and ligand, and DSF:* In PCR strip tubes, appropriate volumes of concentrated protein, compound, and buffer were combined to a final 1X volume of 36 μL, and the resulting solution was mixed well by pipetting up and down. 10 μL of each sample was then transferred in triplicate into a white, low-volume 384-well qPCR plate (Axygen PCR-284-LC480WNFBC, lot # 23517000) using an E100 ClipTip matrix pipette. The plate was spun down for 30 seconds to settle droplets to the bottoms of the wells. and DSF measurement was performed as described in '*Heating and measurement*' (above).

*Data analysis.*

Transition ratios. Ratios were calculated from paDSF experiments in which all constructs were present at 1 μM, and TW408 was present at 50 μM. paDSF detection ratios between isolated catalytic and TPR domains were calculated between the mean maximum RFU intensities reached, and correspond to the peak of the unfolding transition. paDSF detection ratios between transitions 1 and 2 in OGT (corresponding to $T_{ma\_OGT-1}$ and $T_{ma\_OGT-2}$) were calculated between the mean maximum RFU intensities reached by the two respective sigmoidal components of the fitted model from which $T_{ma\_OGT-1}$ and $T_{ma\_OGT-2}$ were calculated.

Statistical tests. Tests for significance in concentration-dependent thermal shifts with UDPGlcNAc or L4 were calculated using a Spearman correlation between ligand concentration and Tma, using stats::cor.test function in R, with method = "spearman". Tests for significance in differences between two groups (e.g. $T_{ma\_OGT-1}$ vs $T_{ma\_cat}$, $T_{ma\_OGT-1}$ vs $T_{ma\_TPR}$) were performed using Two-tailed Students T-Test, using stats::t.test() function in R. In all cases, correlations were considered significant if $p < 0.01$.

Variable temperature UDP-Glo Enzymatic activity assay of OGT

*Material preparation.* Buffer was prepared fresh by adding 4 uL of 1 M DTT to 4 mL of OGT buffer (final buffer: 50 mM HEPES pH 7.4, 300 mM NaCl, 1 mM DTT). Three 4X-concentrated L4 solutions were prepared by adding 1 uL 10 mM L4 in DMSO to 999 uL of buffer. Following thorough mixing, two 150 μL two-fold serial dilutions were prepared, yielding at least 150 uL each of 20, 10, 5, and 0 μM L4 solutions. A 2X-concentrated protein solution was prepared by adding 50 uL of 8 μM purified OGT to 150 μL buffer, yielding 200 μL of 2 μM OGT. The 4X-concentrated sugar-substrate mix was prepared by adding 2 μL of 100 mM Ultra Pure UDP-GlcNAc (Promega Cat # V7071) to 998 uL buffer, followed by the addition of 80 μL of 1 mM substrate peptide (Anaspec Cat. # 63726, Lot#1958507, >95%, sequence: KKKYPGGSTPVSSANMM) to 720 uL of this UDP-GlCNAc buffer, yielding 800 μL of buffer containing 100 μM substrate and 200 μM UDP-GlcNAc.

*Enzymatic activity at variable temperatures.* Enzymatic reaction mixtures were prepared at each of the four L4 concentrations by combining 48 μL 4X OGT, 96 μL 2X sugar-substrate, and 48 μL 4X L4. Final conditions were: 500 nM OGT, 100 μM UDP-GlcNAc, 50 μM substrate peptide, and either 5, 2.5, 1.25, or 0 μM L4. As a negative control, the identical reactions were set up using buffer in place of OGT, yielding final conditions of 0 nM OGT, 100 μM UDP-GlcNAc, 50 μM substrate peptide, and either 5, 2.5, 1.25, or 0 μM L4. To monitor catalytic activity across a range of temperatures, all reactions were immediately divided into 22 μL aliquots across 8 PCR strip tubes, and these tubes were placed in a PCR thermocycler, and incubated across a gradient of 30 - 50 ºC with a heated lid for 140 minutes.

*Quenching, detection, and data collection of enzymatic activity.* During incubation, the Promega UDP-Glo reagent Promega UDP-Glo Glycosyltransferase assay (Cat # V6961) was prepared. The "UDP-Glo working solution" was prepared by addition of 1 μL "UDP-Glo enzyme" to 74 "Enzyme Dilution Buffer". The "UDP detection reagent" was prepared by adding 10 μL of this "UDP-Glo working solution" to 990 μL "Nucleotide Detection Reagent". Following the 140 min incubation, all samples were rapidly cooled to 12 ºC in the thermocycler, and 5 μL of each solution was immediately transferred into a well of a white, low-volume, hard-sided 384-well plate (Greiner BioOne #784075, Lot 1209043), and 5 μL of the prepared "UDP detection reagent" was added to all wells, simultaneously quenching OGT activity and beginning the UDP detection reaction. The plate was shaken gently by hand to mix, centrifuged for 1 min at 1400 rcf to remove bubbles, and incubated at room temperature in the dark for 65 minutes. Following incubation, resulting luminescence was recorded on a Tecan Spark plate reader.

Calculation of dye physical chemical properties

In python, the SMILES string for each dye was converted to a mol object using Chem.MolFromSmiles. For each mol object, the number of hydrogen bond donors, hydrogen bond acceptors, and heteroatoms were calculated using the rdkit.Chem.Lipinski module; cLogP and Molecular Refractivity were calculated using rdkit.Chem.Crippen module, and molecular weight and Topological Polar Surface Area were calculated using the MolWt and TPSA functions from the rdkit.Chem.Descriptors module respectively. In R, the Pearson correlation was calculated between the dye hit rate and each calculated property using cor.test.

Calculation of ECFP4 Tanimoto coefficients and clustering

SMILES strings of all dyes were cannonicalized using Chem.CannonSmiles from the 'RDKit' package in python 3. From cannonicalized SMILES, pairwise ECFP4 Tanimoto Coefficient was calculated using the Morgan algorithm implemented in the 'RDKit' package in python 3. The resulting pairwise similarity matrix was converted to dissimilarities by subtracting each value from 1. This dissimilarity matrix was passed to hierarchical cluster analysis using Lance-Williams dissimilarity and 'complete' linkage clustering method via the 'hclust' function implemented in R package in the 'stats' package. Similarities were calculated within four Chembridge Screening Libraries, a 50K compound "diversity" library (DiverSET), and three "focused" libraries "GPCR", "Ioncore", and "NHRCore", in the same manner.

*Scaffold extraction.* To define chemical scaffolds, the same pairwise matrix was passed to Butina clustering, implemented using rdkit.ML.Cluster.Butina.ClusterData with cutoff = 0.7.

*Calculation of dye "hit rate".* Dye "hit rate" was calculated by dividing the number of proteins for which a dye was called a "hit" from the number of proteins against which it was tested. Pairwise dye hit rate similarity was calculated as the absolute value of the hit rate between two dyes, and a complete pairwise similarity matrix was calculated for the library. For example, if a dye was screened against 70 proteins, and was a "hit" for 45 of them, the hit rate for that dye would be 45/70, or 64.3 percent. Dye "sensitivity hit rate" was calculated in the same way, but using the number of proteins for which a dye was called "sensitive" instead of "hit."

Calculation of dye hit profile similarities

The pairwise similarity between paDSF dye screen results for different proteins was calculated using the Jaccard coefficient, J(A,B) =|A  B||A  B|, where A and B are each the set of outcomes (hit, sensitive, or none) for proteins for which a different dye was tested. Specifically, the union was defined as the number of dyes which received the same assignment between the two proteins (both hit, both sensitive, or both none). The intersection was defined as the number of dyes which were tested against both proteins, regardless of the assignment. Higher jaccard coefficients indicate more similar dye screen results: proteins which received different assignments for every dye would have J = 0, while proteins which received the same assignment for every dyes would have J = 1. Dyes which never received an assignment of "hit" or "sensitive" for any tested protein (103 dyes) were not were not included in this calculation. For each protein pair, the Jaccard coefficient was calculated using only the dyes which were tested against both proteins. The final dataset includes 2,211 unique comparisons made between the 67 screened proteins.

Calculation of correlation between protein sequence and dye fingerprint

Proteins which contained multiple distinct chains (eIF2B, nsp10/nsp16, nsp10/nsp14, proteasome, and nucleosome) were removed prior to calculation, as it was not possible to distinguish the contribution of each chain to the dye screen results. For the remaining set of all unique protein pairs (67 proteins, 2,211 unique pairs), the Spearman correlation between dye screen results (see above) and primary sequence similarity (see above) using the base::cor.test(method = "spearman") function in R.

**References.**

43. E. Thompson, thesis, University of California, San Francisco, Ann Arbor, United States (2020).

44. S. Militi, E. S. Maywood, C. R. Sandate, J. E. Chesham, A. R. Barnard, M. J. Parsons, J. L. Vibert, G. M. Joynson, C. L. Partch, M. H. Hastings, P. M. Nolan, Early doors (Edo) mutant mouse reveals the importance of period 2 (PER2) PAS domain structure for circadian

**Caption for Movie S1**

**Movie S1. Visual protocol for dye screen.** Annotated video of the primary liquid handling steps of a dye screen.

**Captions for Data S1 to S4**

**Data S1. Chemical structures and common names of Aurora dyes.** See Data S3 for more information, includes specific dye commercial sources, CAS numbers, and MSDS where available.

**Data S2. Full dye screen results for all tested proteins.** These results can also be explored interactively at https://padsfdyes.shinyapps.io/Exp1243_heatmap_cache/.

**Data S3. Primary sequences of protein panel constructs.** See Table S1 for more information on sources and biochemical properties of proteins.

**Data S4. Extended information on Aurora dyes.** A spreadsheet containing, for each dye in the library: Aurora-library name (e.g. L095), common name (where applicable), CAS, source, catalog number, MSDS, and SMILES. the 312 dyes in the Aurora library.

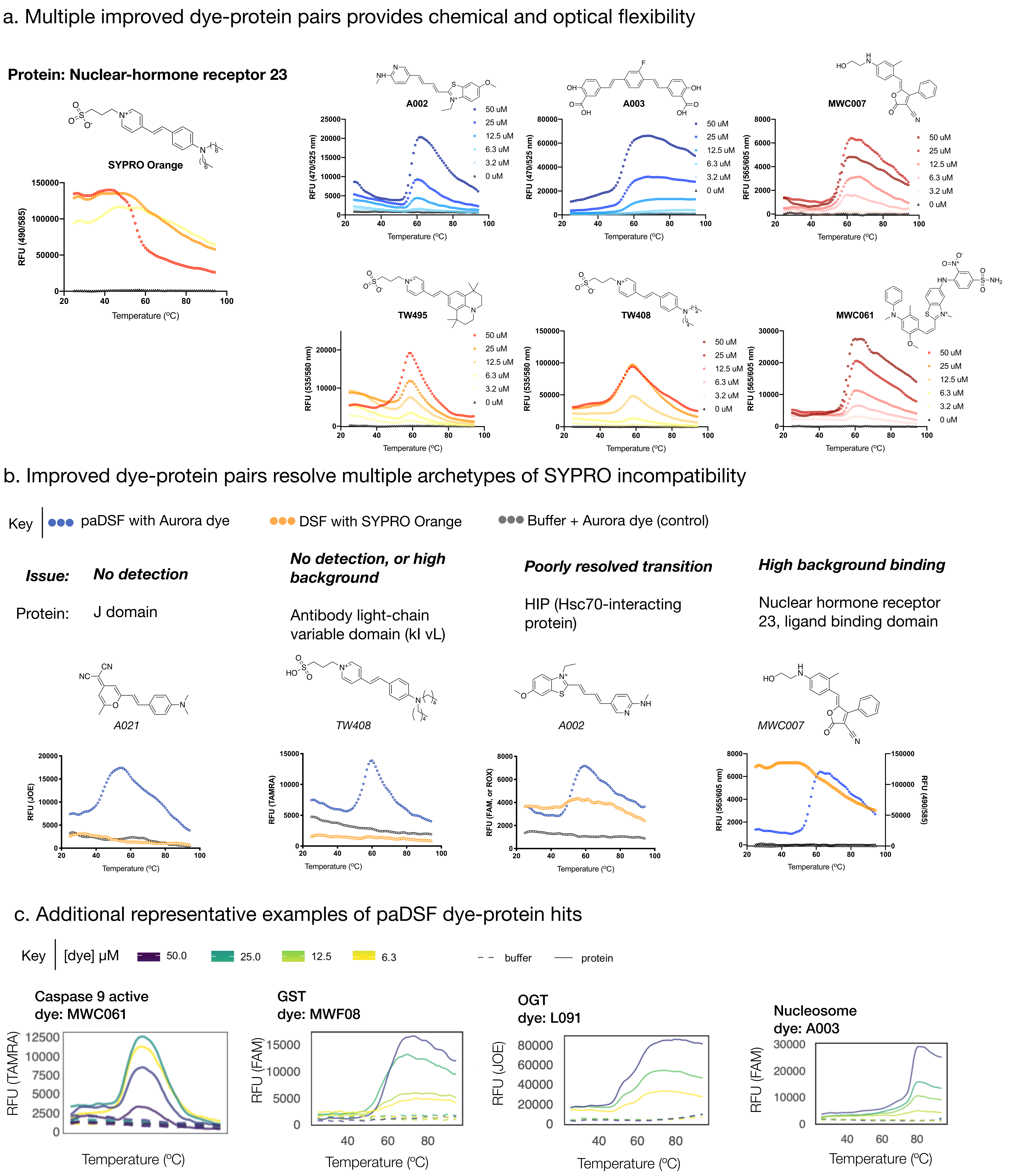


**Fig. S1. Representative examples of paDSF screening results.** For full screening results for all proteins, see Data S2. **(A)** Results of a dye screen against the SYPRO-incompatible protein nuclear-hormone receptor 23. Far left panel: representative unsuccessful DSF conducted with SYPRO Orange. Right panels: examples of successful paDSF data, collected with six different paDSF dyes. Plots show dyes at four different concentrations, demonstrating a dose-responsive relationship between RFU and dye concentration. **(B)** Representative examples of resolution of four archetypes of SYPRO Orange incompatibility by paDSF dyes. **(C)** Examples of successful paDSF data collected for four additional proteins. As in panel A, dyes are plots show dyes at four different concentrations, demonstrating a dose-responsive relationship between RFU and dye concentration.

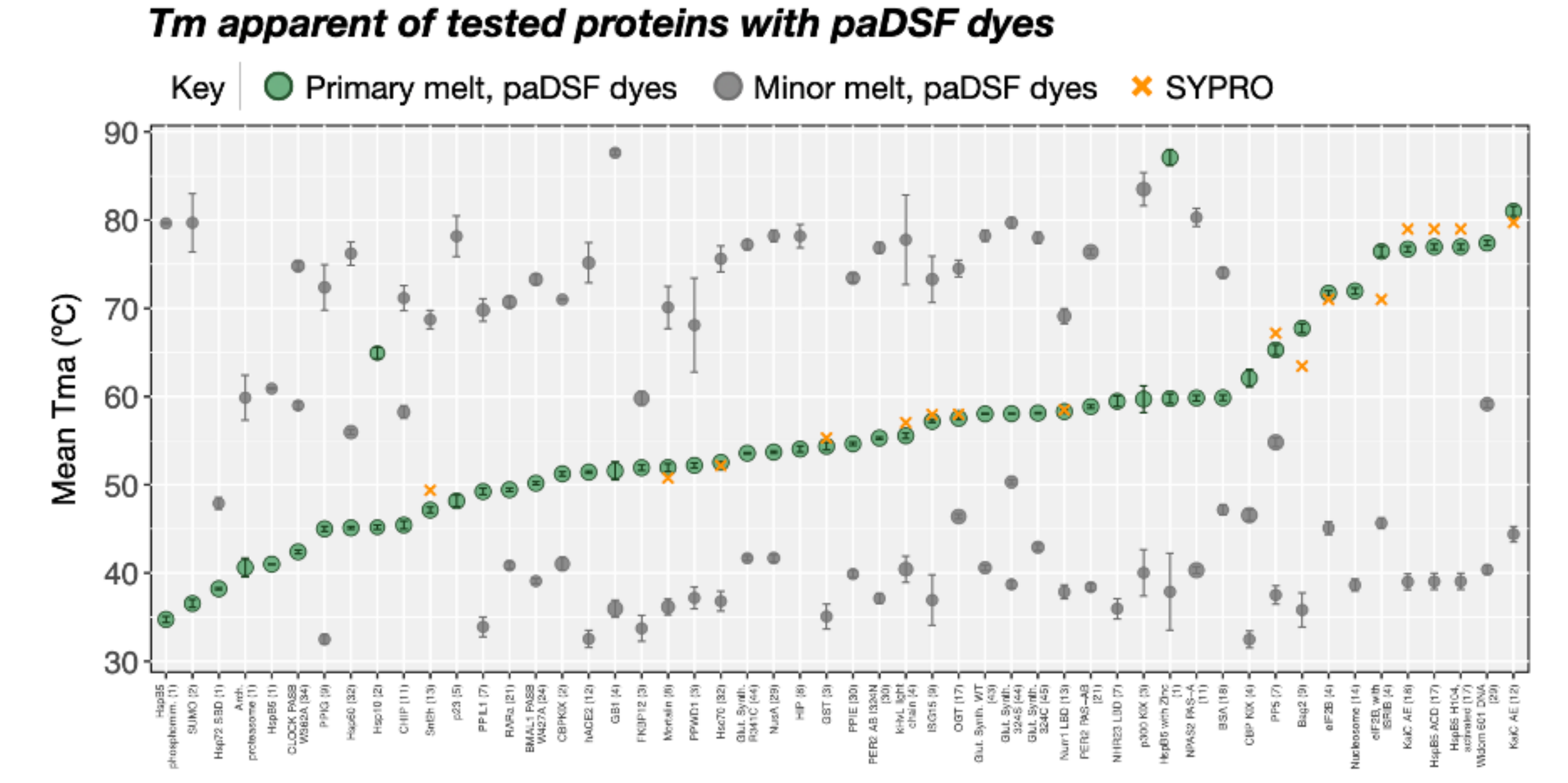


**Fig. S2. Apparent melting temperatures for paDSF dyes versus SYPRO Orange.** Tmas reported by compatible dyes for all tested proteins (single point: single compatible dye), compared to SYPRO Orange values. Error bars represent +/- standard error.

*See end of document for* **Table S1. Chemical structures and common names for all Aurora library dyes.** Chemical structures, Aurora library name, and common name (where applicable) for all 312 tested dyes in the Aurora library. Unless otherwise noted, dyes are available either commercially, or through the North Carolina State Max Weaver Dye Library. See Data S1 for extended information on dyes and sources.

*See end of document for* **Fig. S3. Apparent melting temperatures for paDSF dyes versus SYPRO Orange.** Tmas reported by compatible dyes for all tested proteins (single point: single compatible dye), compared to SYPRO Orange values. Error bars represent +/- standard error.

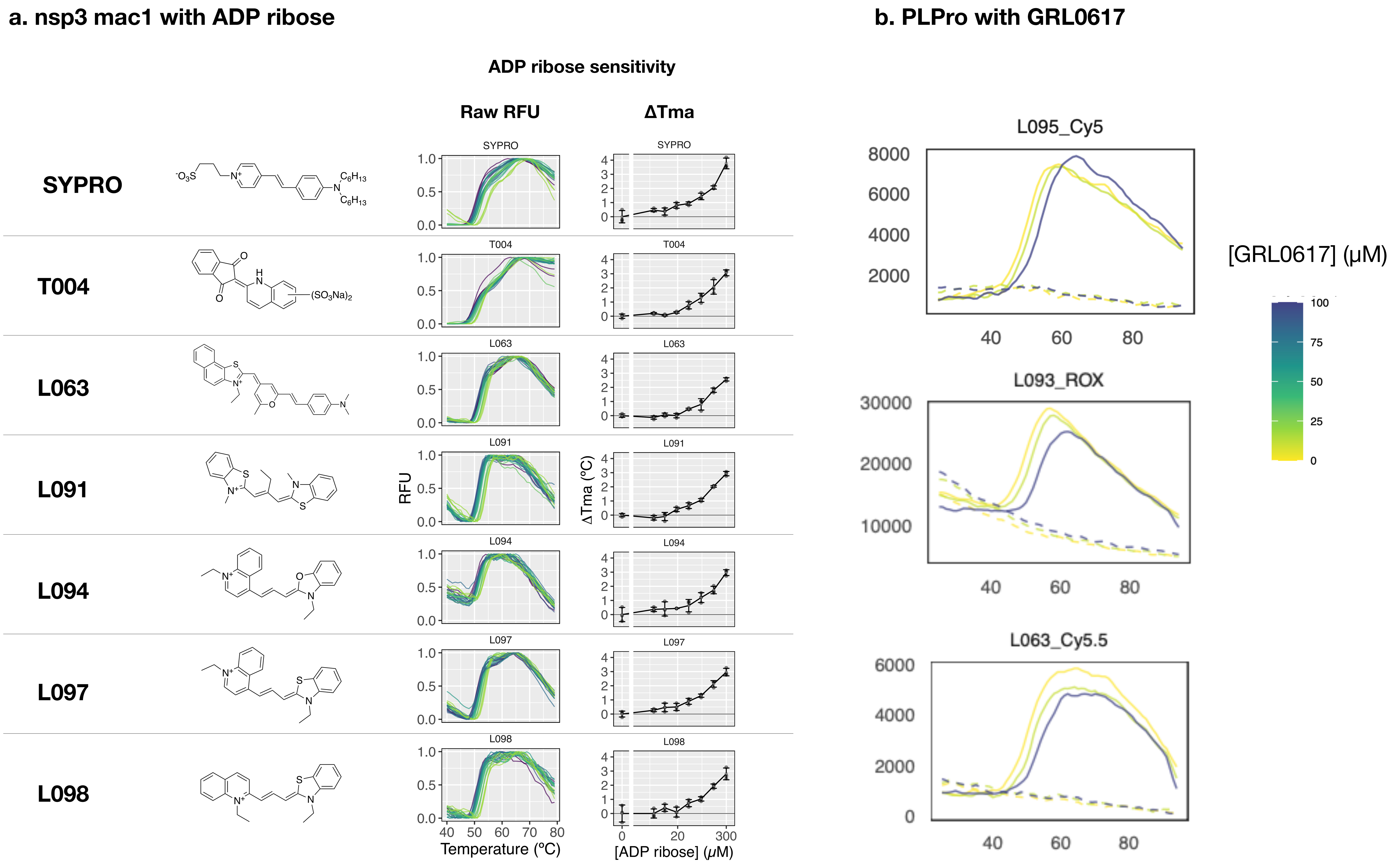


**Fig. S4. Apparent thermal shifts with paDSF dyes for SARS-CoV2 proteins.** (A) Thermal upshift in the presence of ADP ribose with nsp3 mac1, with 6 paDSF dyes and SYPRO orange. Error bars represent +/- standard deviation. (B) Thermal upshift in the presence of GRL0617 with PLPro, with 3 paDSF dyes.

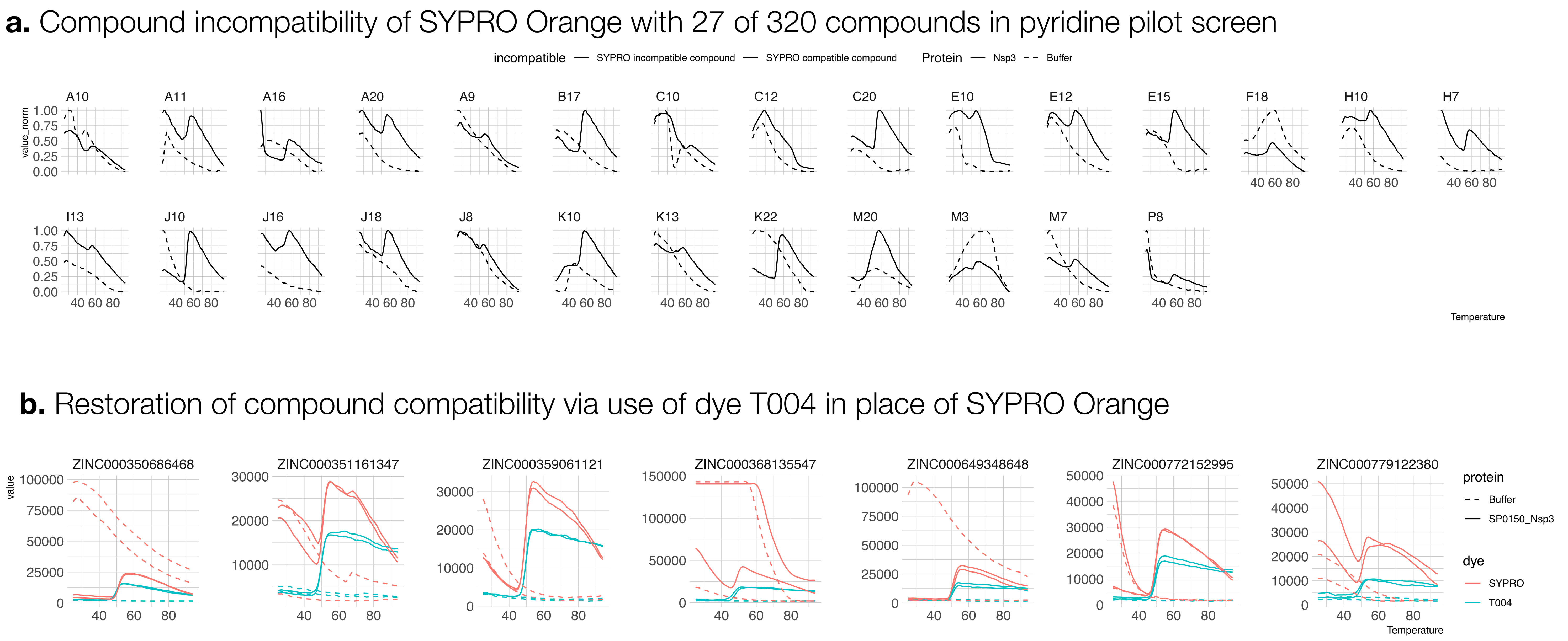


**Fig. S5. Reduction in artifactual dye activation by screening compounds for nsp3 mac1.** (**A**) Protein-independent activation of SYPRO Orange observed for compounds which produced uninterpretable raw DSF data from a 320-compound pyridine pilot screen using SYPRO Orange. (**B**) Comparison of DSF data collected using SYPRO Orange and dye T004, demonstrating improved compound compatibility of T004 over SYPRO Orange.

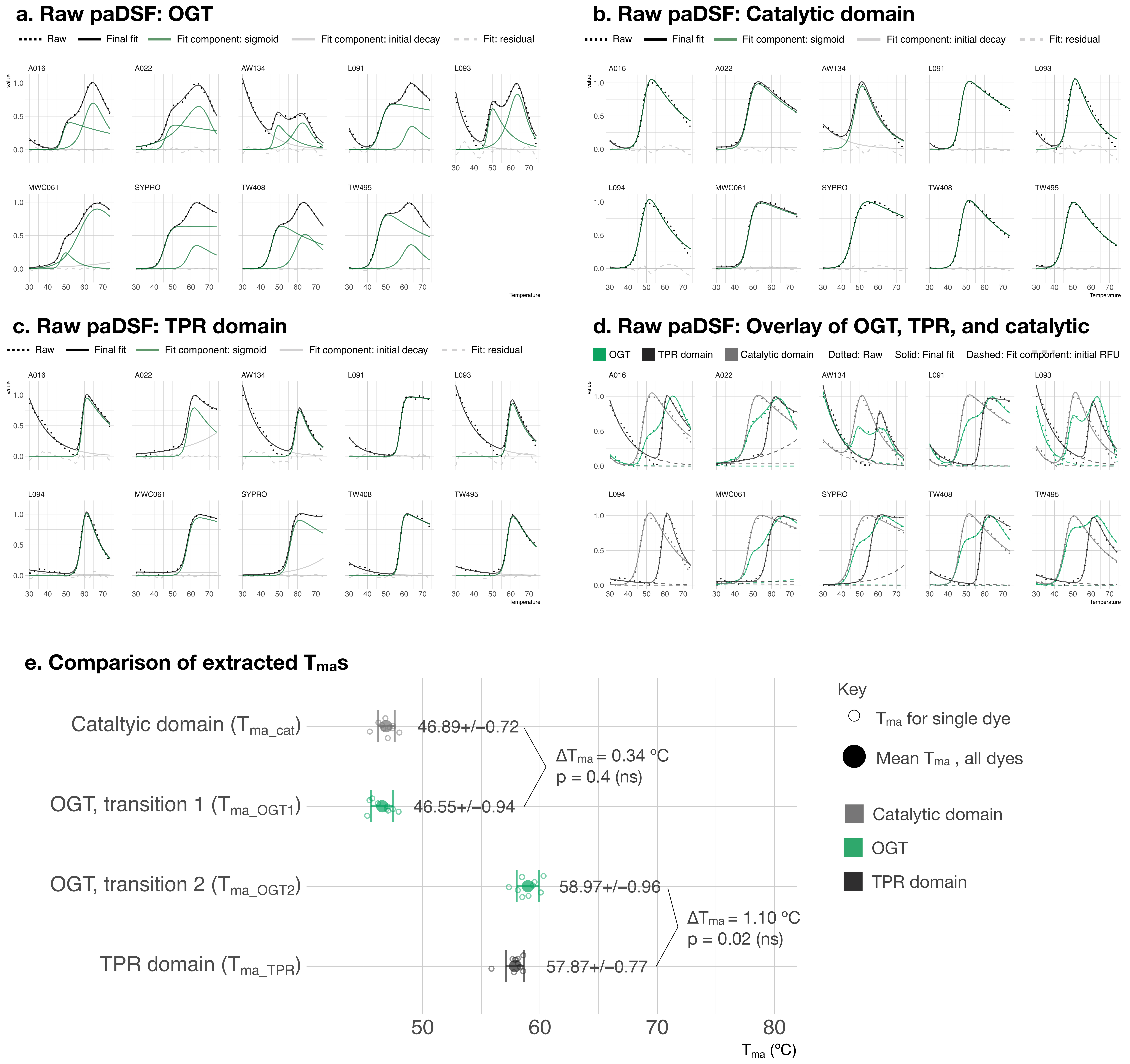


**Fig. S6. Raw and analyzed paDSF with OGT, catalytic domains and TPR domain.** paDSF results from the 10 hit dyes for OGT, with raw RFU data displayed alongside fitted models for (**A**) OGT alone, fitted to DSFworld Model 4 (two sigmoids, with initial RFU (**B**) catalytic domain alone, fitted to DSFworld Model 2 (one sigmoids, with initial RFU (**C**) TPR domain alone, fitted to DSFworld Model 2 (one sigmoids, with initial RFU (**D**) overlay of raw data and final fits for all three constructs. (**E**) $T_{ma}$s for the three constructs, extracted using the fits displayed in panels A-D. Data are presented as the mean of three technical replicates +/- standard deviation. No significant (ns) difference was found between $T_{ma\ OGT\text{-}1}$ and $T_{ma\ cat}$, nor $T_{ma\ OGT\text{-}2}$ and $T_{ma\ TPR}$. Significance determined using two-tailed t-test, and a significance threshold of $p < 0.01$.

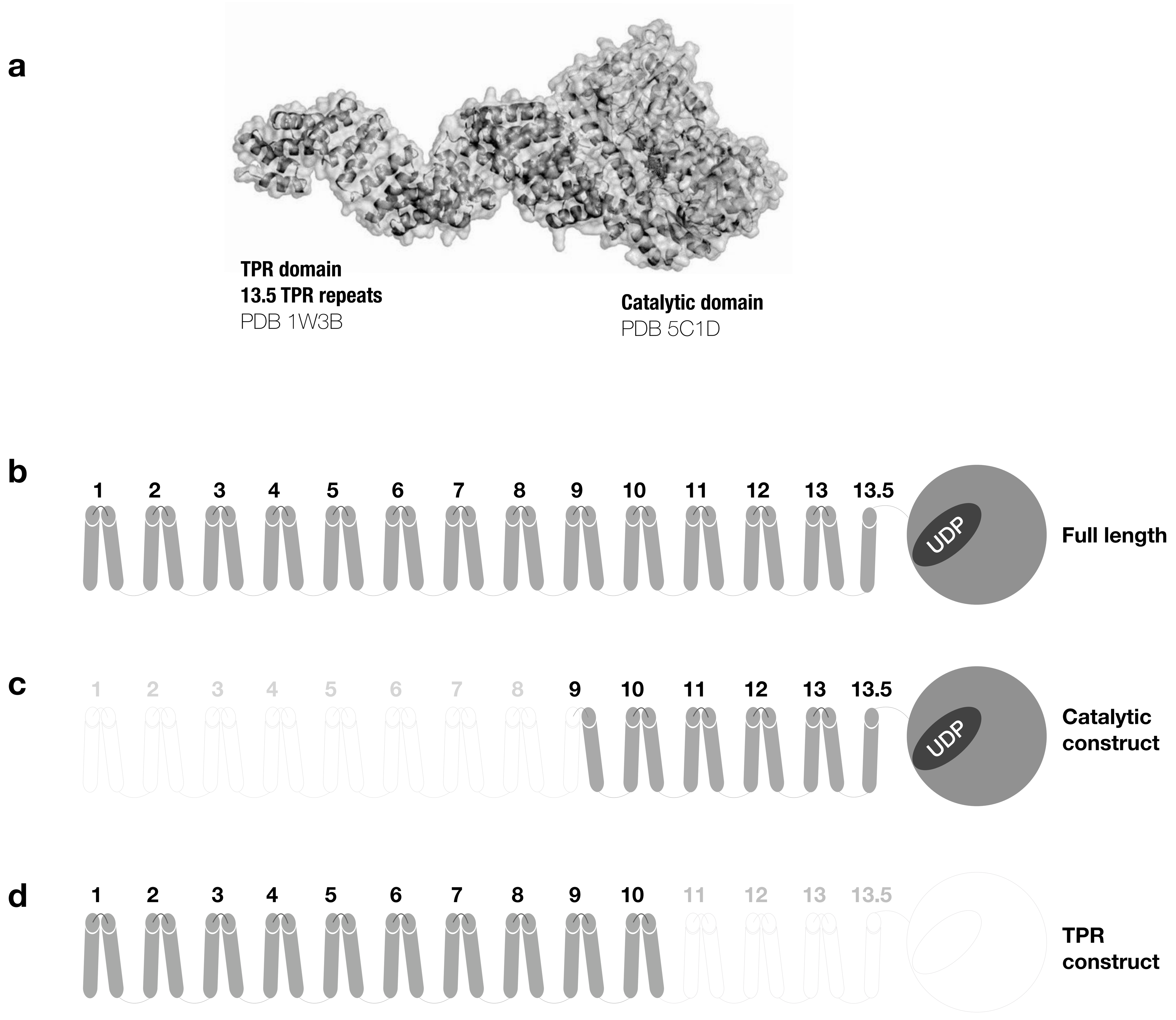


**Fig. S7. Diagram of OGT constructs used in this study.** (**A**) Structural model of full-length OGT, composed from crystal structures of TPR (PDB 1W3) and catalytic (PDB 5C1D) domains. Cartoons of the (**B**) full-length OGT construct, which includes the full TPR and catalytic domains, (**C**) catalytic domain construct, which includes the full catalytic domain and TPR repeats 9 - 13.5, and (**D**) TPR domain construct, which includes TPR repeats 1-10.

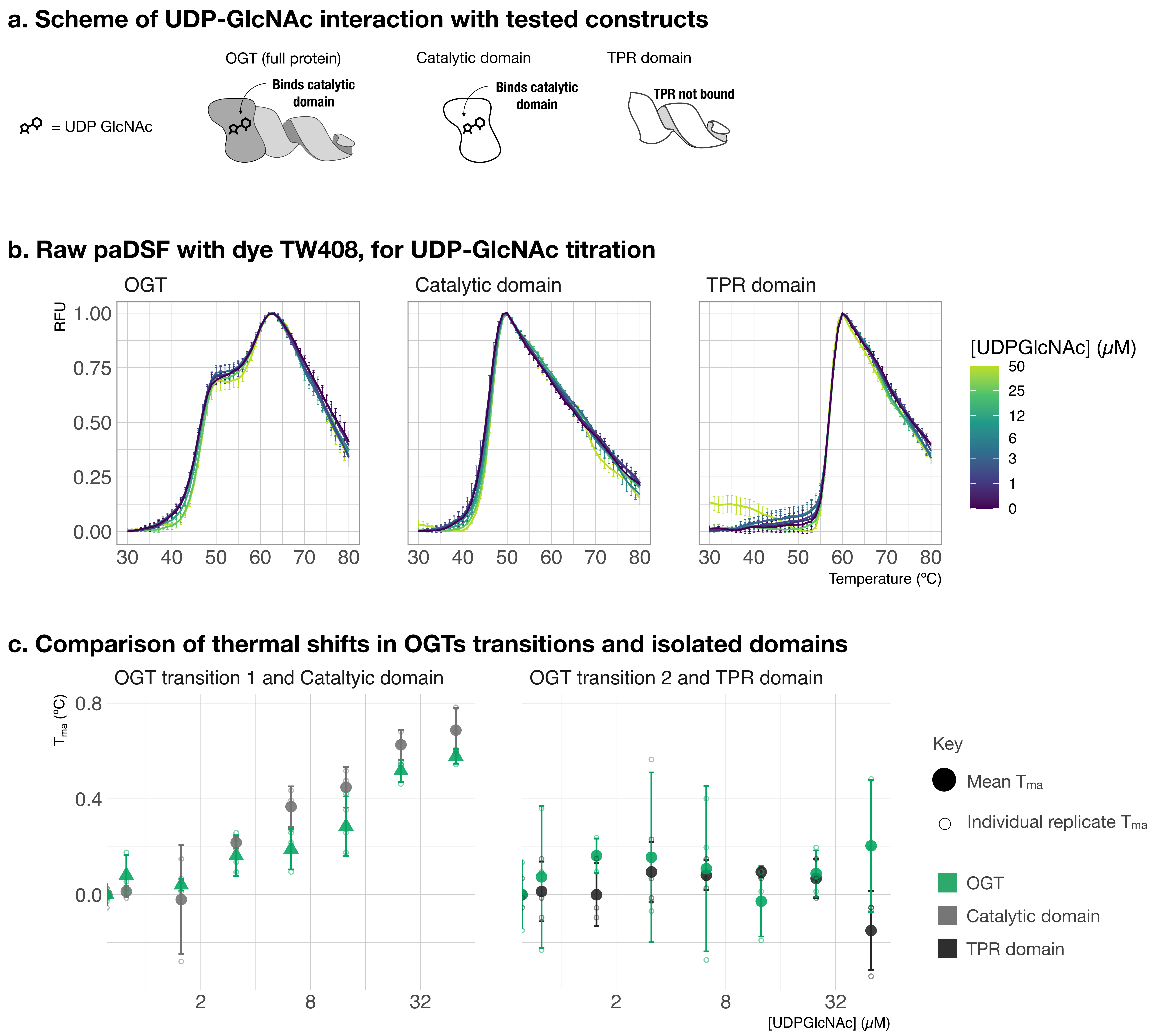


**Fig. S8. Thermal shifts with UDP-GlcNAc from TW408 paDSF.** paDSF results using dye TW408 to monitor titration of UDP-GlcNAc with OGT, catalytic domain, or TPR domain. (**A**) Schematic of interaction of UDP-GlcNAc with full length and isolated catalytic domain, but not TPR domain, of OGT. (**B**) Raw RFU for each construct in the presence of 0 - 50 µM UltraPure UDP-GlcNAc. (**C**) Comparison of resulting Tmas. Left panel: $T_{ma\ OGT1}$ and $T_{ma\ cat}$; right panel: $T_{ma\ OGT2}$ and $T_{ma\ TPR}$. Both $T_{ma\ OGT1}$ and $T_{ma\ cat}$ show significant thermal upshift with UDP-GlcNAc ($T_{ma\ OGT1}$ spearman estimate = 0.92, p = 1.13e-10; $T_{ma\ cat}$ spearman estimate = 0.94, p = 1.48e-12); while neither $T_{ma\ OGT2}$ and $T_{ma\ TPR}$ do ($T_{ma\ OGT2}$ spearman estimate = 0.095, p = 0.66; $T_{ma\ TPR}$ spearman estimate = 0.005, p = 0.98). Data are presented as the mean of three technical replicates +/- standard deviation.

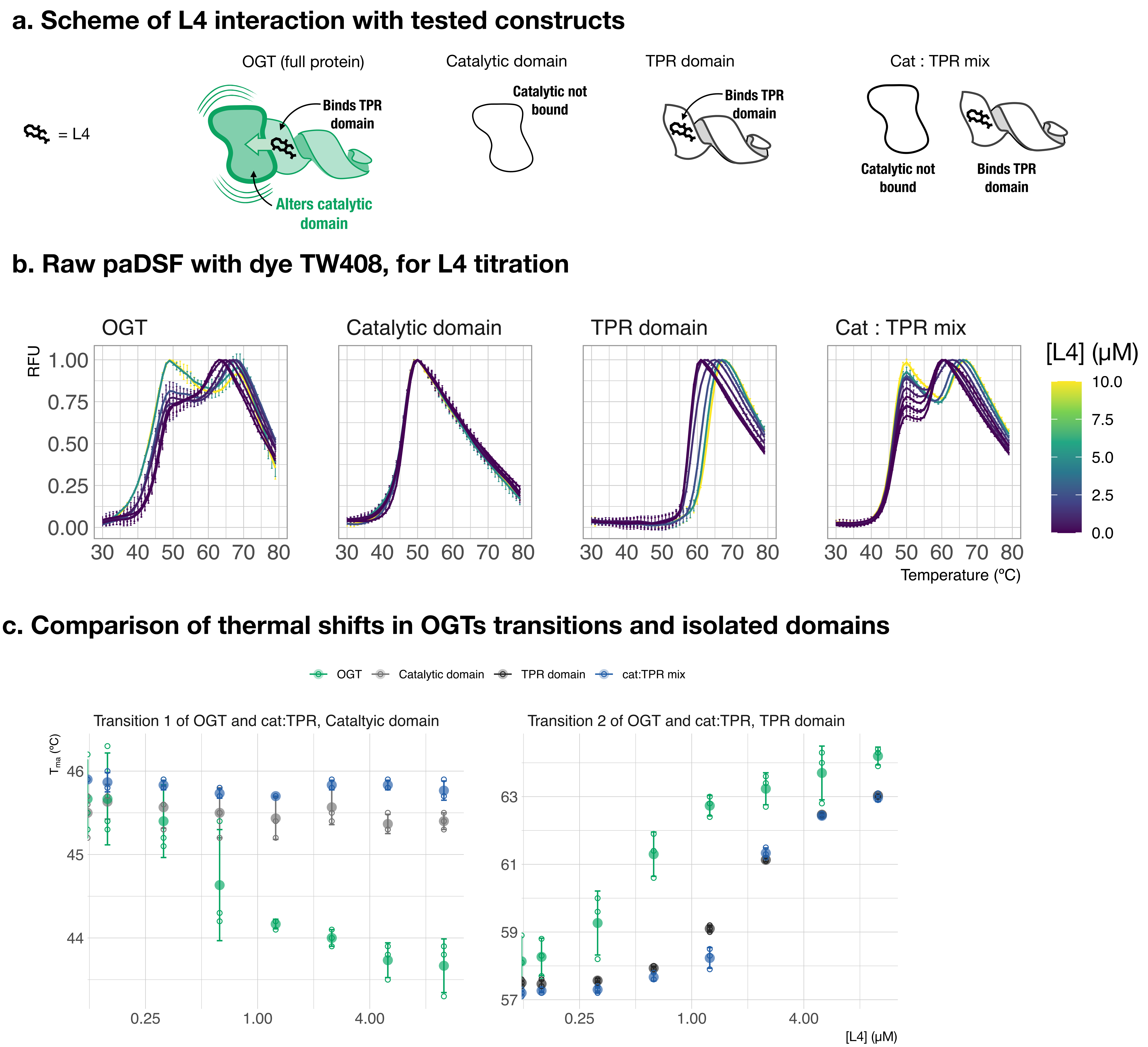


**Fig. S9. Thermal shifts with L4 from TW408 paDSF.** paDSF results using dye TW408 to monitor titration of L4 with OGT, catalytic domain, or TPR domain. (**A**) Schematic of L4 with the four tested conditions: full OGT, catalytic domain, TPR domain, and a 1:1 mix of catalytic and TPR domain. All constructs were tested at 1 μM final; 1:1 mix had 1μM catalytic domain and 1 mM TPR domain. (**B**) Raw RFU for each construct in the presence of 0 - 10 μM L4. (**C**) Comparison of resulting Tmas. Left panel: $T_{ma\ OGT1}$, $T_{ma\ cat:TPR\ 1}$, and $T_{ma\ cat}$; right panel: $T_{ma\ OGT2}$, $T_{ma\ cat:TPR\ 2}$, and $T_{ma\ TPR}$. Significant thermal upshifts with L4 treatment observed with $T_{ma\ OGT2}$ , $T_{ma\ cat:TPR\ 2}$, and $T_{ma\ TPR}$ ($T_{ma\ OGT2}$ spearman estimate = 0,95, p = 2.3e-12; $T_{ma\ cat:TPR\ 2}$ spearman estimate = 0.97, p = 4.7e-15; $T_{ma\ TPR}$ spearman estimate = 0.96, p = 3.6e-14). Significant thermal downshift with L4 observed in $T_{ma\ OGT1}$ (data in Fig. 3D, included here for reference, $T_{ma\ OGT1}$ spearman estimate = -0.92, p = 6.2e-11). No dose-responsive thermal shift with L4 observed in $T_{ma\ cat}$ or $T_{ma\ cat:TPR\ 1}$ ($T_{ma\ cat}$ spearman estimate = -0.27, p = 0.2; $T_{ma\ cat:TPR\ 1}$ spearman estimate = -0.34, p = 0.09). Data are presented as the mean of three technical replicates +/- standard deviation.

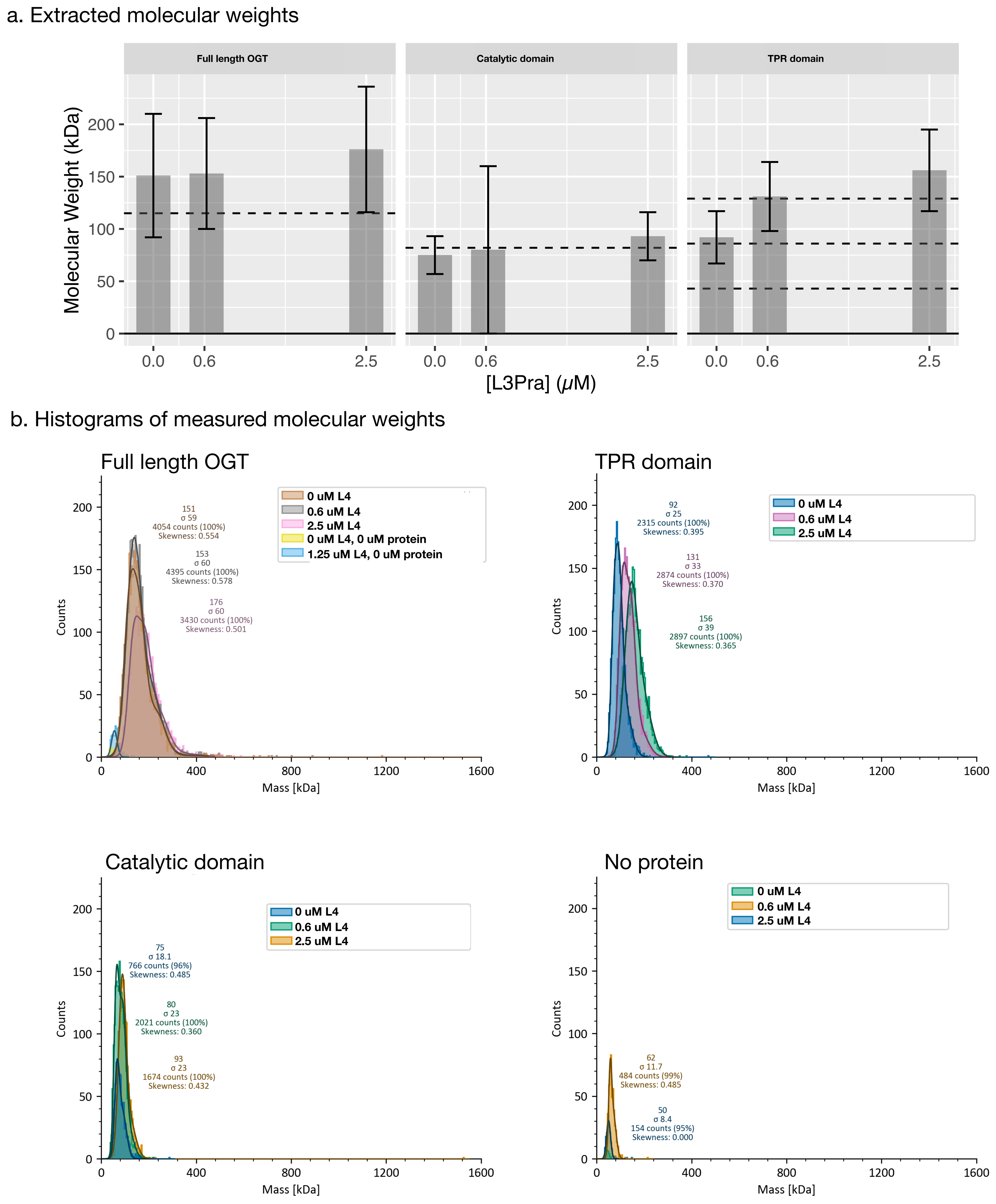


**Fig. S10. OGT oligomerization state and molecular mass not are affected by L4.** (**A**) Molecular weights of full length OGT, catalytic domain, and TPR domain, extracted from mass photometry data. Dashed lines: expected molecular weights for each species. OGT: 115 kDa, Catalytic domain; 82 kDa, TPR domain: 43, 86, 129 kDa (monomer, dimer, trimer). Error bars: +/- standard error in mass from mass photometry. (**B**) Histograms of counts of MW from mass photometry.

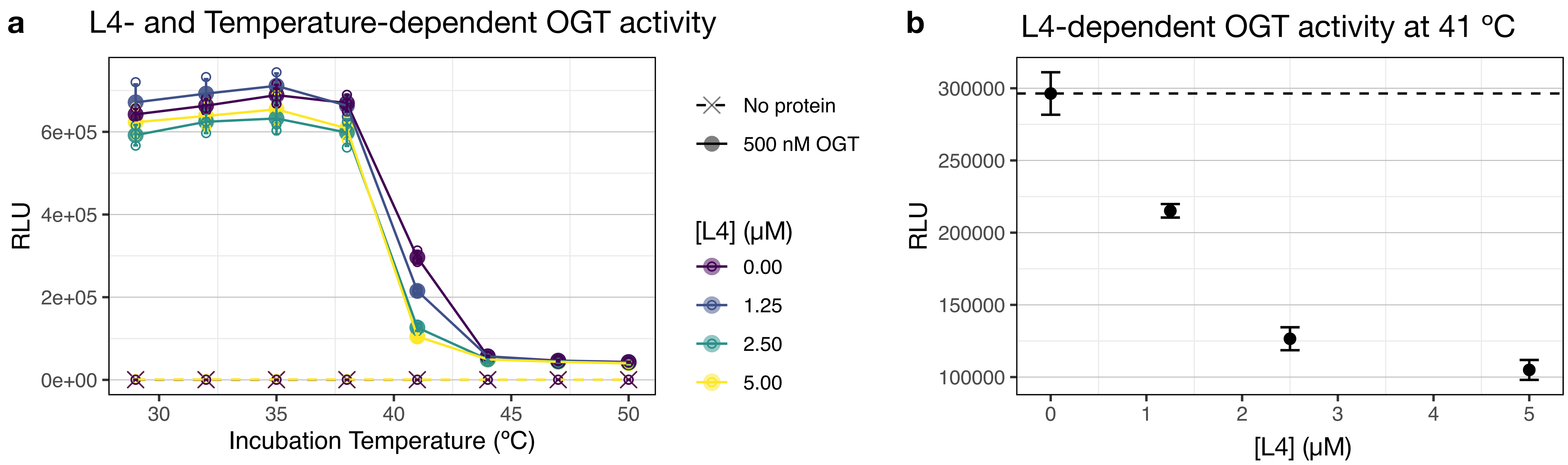


**Fig. S11. L4 treatment increases temperature sensitivity of OGT catalytic activity.** OGT catalytic activity with a model substrate peptide (KKKYPGGSTPVSSANMM), with enzymatic activity allowed to proceed at 29, 32, 35, 38, 41, 44, 47, or 50 ºC. Activity was measured as RLU using Promega UDP-Glo Assay, with higher RLU values corresponding to higher activity. (**A**) Comparison of temperature-dependent loss of enzymatic activity (RLU) in the presence of increasing concentrations of L4 Solid lines and dots: OGT-containing samples; Dashed lines and Xs: protein-free control. (**B**) L4-dependent increase in temperature sensitivity of catalytic activity at 41 ºC. In all panels, data are presented as the mean of three technical replicates +/- standard deviation.

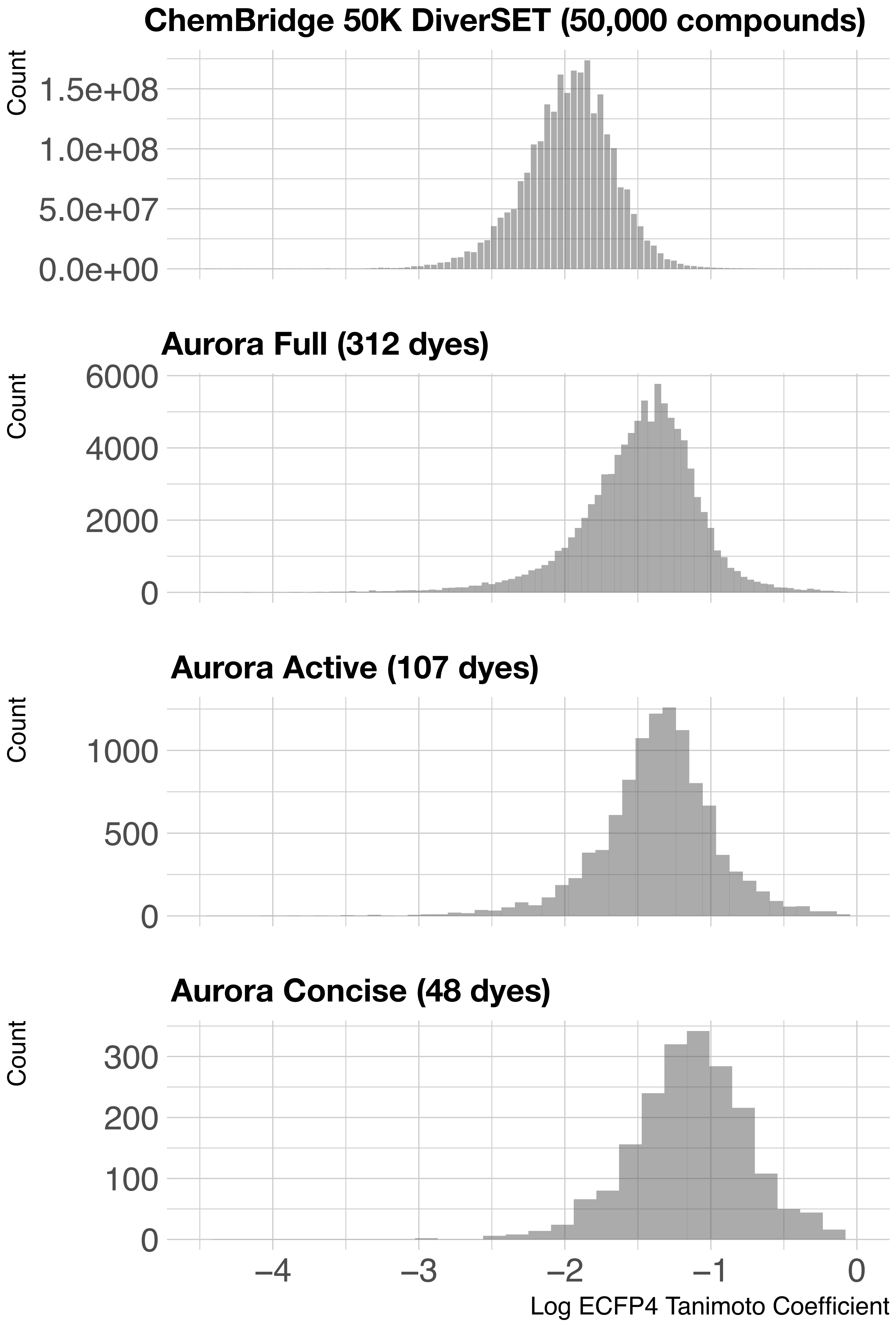


**Fig. S12. Chemical diversity of paDSF dyes versus ChemBridge DIVERSet library.**

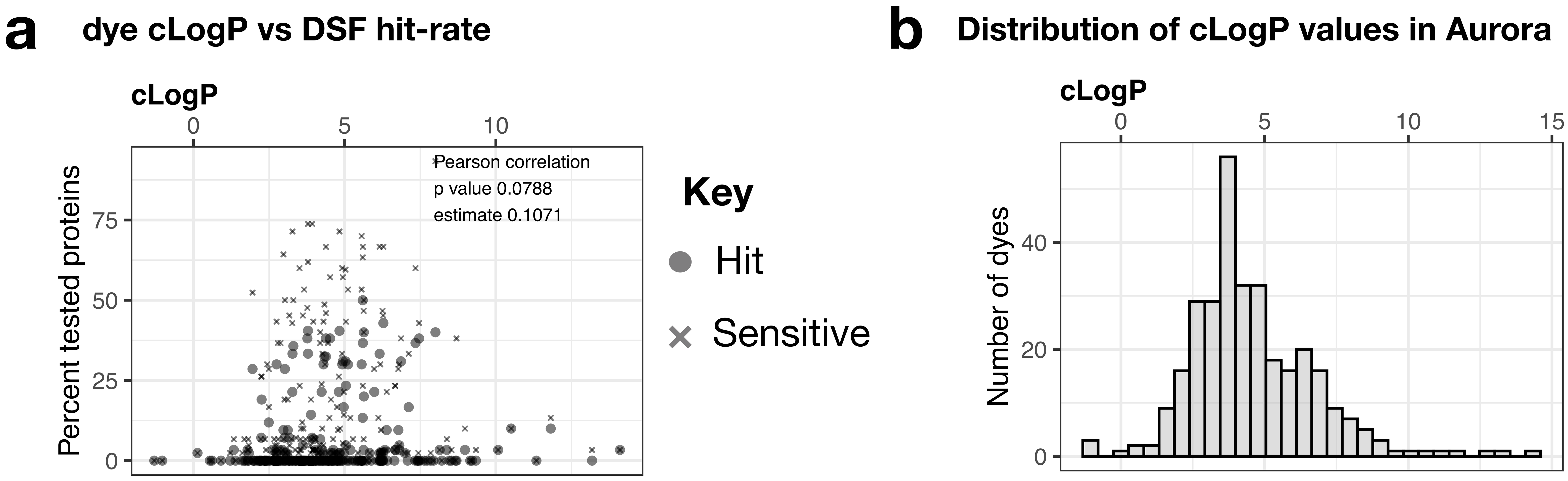


**Fig. S13. cLogP of dyes and relationship to paDSF performance.** (**A**) Comparison of cLogP, a calculated predictor of water solubility, to the paDSF performance of that dye. Filled circles: percent of tested proteins for which a dye was called a “hit”. Xs: percent of tested proteins for which a dye was called “sensitive”. No significant correlation was found between hit rate and cLogP (Pearson p value > 0.05, see panel for exact values) (**B**) Distribution of cLogP values for the full Aurora library (312 dyes), for reference.

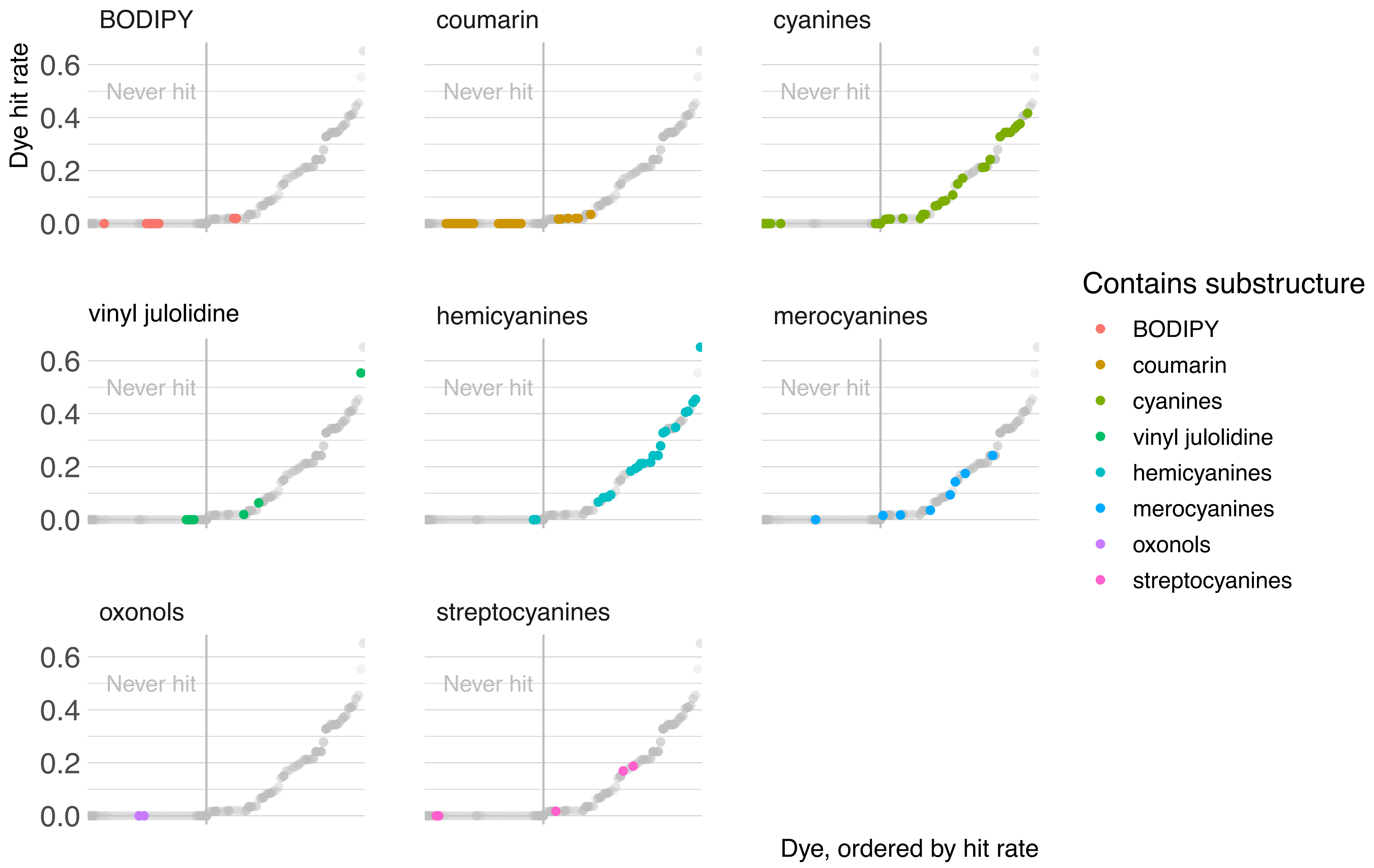


**Fig. S14. Presence and absence of known fluorogenic cores in paDSF dyes.** Individual points represent a single dye, and dyes are arranged along the axis in order of increasing hit rate. Data are shown for the full Aurora library (312 dyes). Colored points indicate dyes which contain a given substructure, grey points indicate dyes lacking that substructure. A singe substructure is presented in each panel, and panels are shown only for substructures present in at least one dye. All points left of the grey vertical line, in the area labeled "Never hit" were never a hit with any tested protein. Some dyes contain multiple substructures.

| Fluorogenic motif | | Dyes with motif | | | Assignments from dyes with this motif | | |
|---|---|---|---|---|---|---|---|
| **Name** | **Structure** | **Active** | **Inactive** | **Total** | **Hit** | **None** | **Percent of all hits** |
| | | *Summary* | | | | | |
| No queried motif | | 44 | 158 | 202 | 207 | 9,330 | 23 |
| Any queried motif | See below | 63 | 48 | 111 | 704 | 4,581 | 77 |
| | | *By substructure* | | | | | |
| BODIPY | 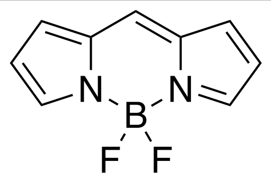 | 2 | 7 | 9 | 2 | 412 | 0.06 |
| 6DMN | 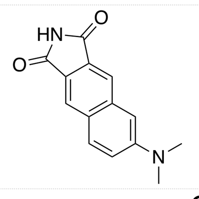 | 0 | 0 | 0 | 0 | | 0 |
| Prodan | 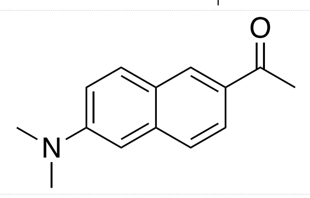 | 0 | 0 | 0 | 0 | | 0 |
| Anthradan | 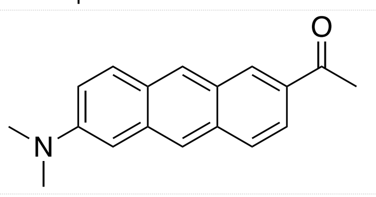 | 0 | 0 | 0 | 0 | | 0 |
| Dansyl | 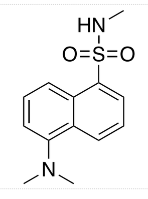 | 0 | 0 | 0 | 0 | | 0 |
| Coumarin | 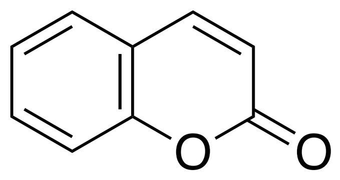 | 6 | 22 | 28 | 7 | 1,333 | 0.2 |
| Vinyl julolidine | 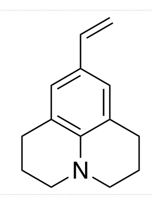 | 3 | 4 | 7 | 41 | 268 | 1.4 |
| Cyanine | 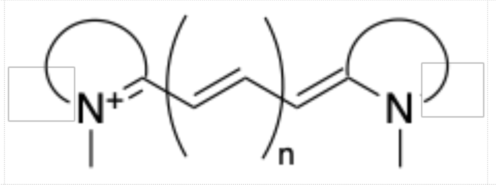 | 25 | 8 | 33 | 1,463 | 8,121 | 49 |
| Hemicyanine | 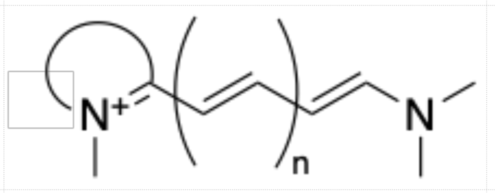 | 21 | 2 | 23 | 1,368 | 2,694 | 46 |
| Streptocyanine | 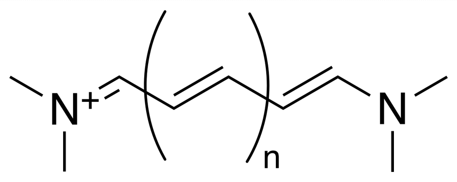 | 3 | 2 | 5 | 47 | 429 | 47 |
| Merocyanine | 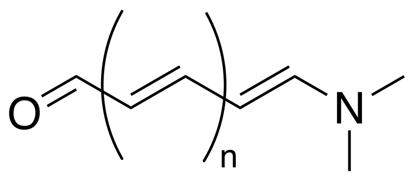 | 7 | 1 | 8 | 73 | 848 | 2.4 |
| Oxonol | 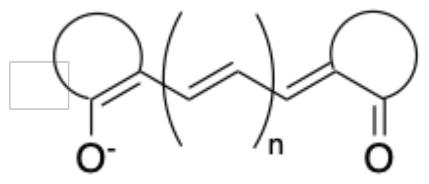 | 0 | 0 | 0 | 0 | | 0 |

**Table S3. Presence and activity of queried fluorogenic scaffolds in the full Aurora dye library.**

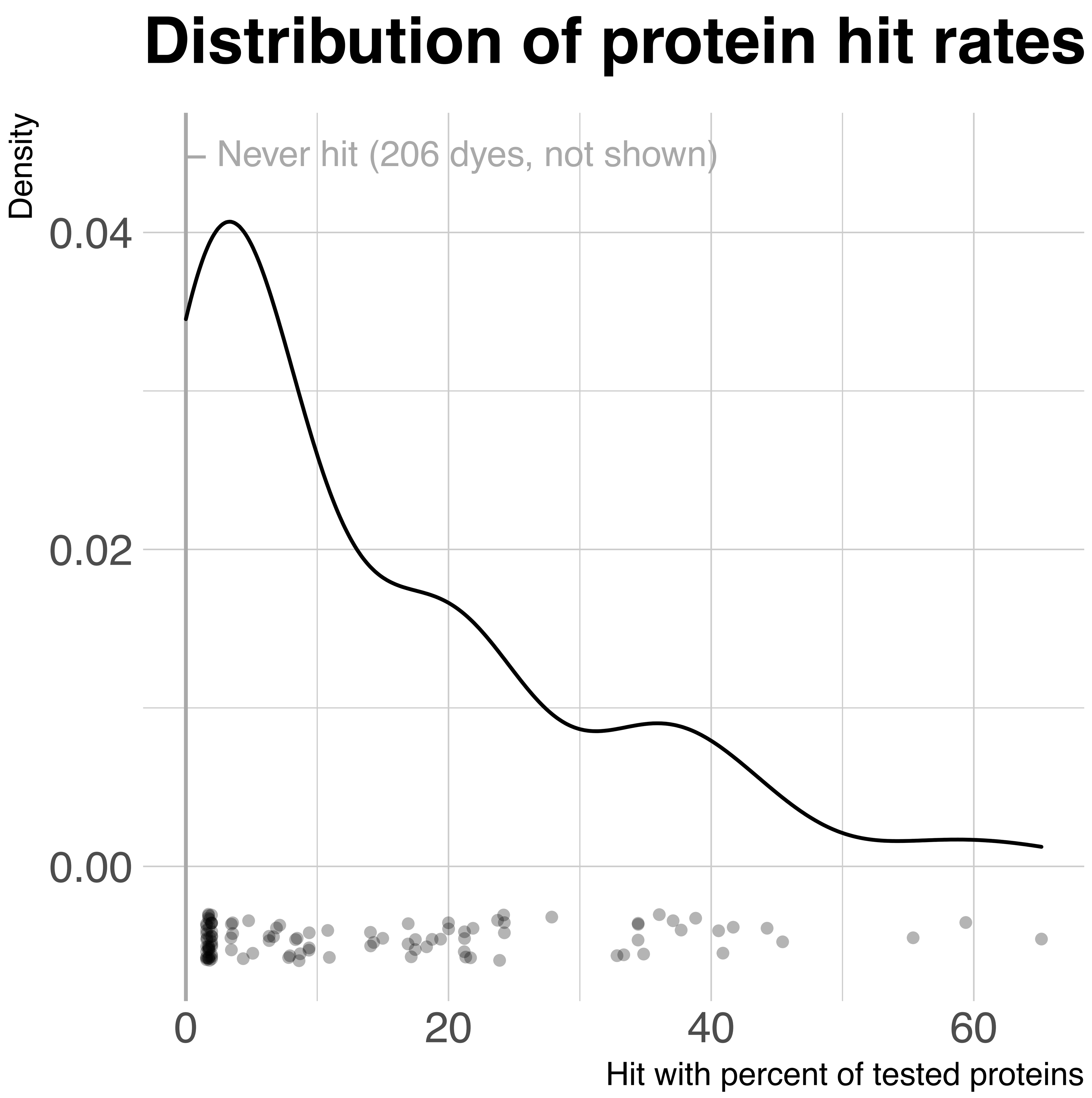


**Fig. S15. Distribution of protein hit rates of individual dyes.** Density plot: the distribution of hit rates across 107 dyes which were assigned “hit" with at least one tested protein (Aurora-active). Points: single points represent the protein hit rates for individual dyes. For dyes which were assigned “hit” with at least one tested protein, the protein selectivities ranged from detection of 1 protein out of 63 tested at the most selective, to three dyes which detected over 50 percent of tested proteins at the most promiscuous. There were 206 dyes which showed no activity with any tested protein (maximum of 64 different proteins). These 206 dyes were not included in the plot or density calculation.

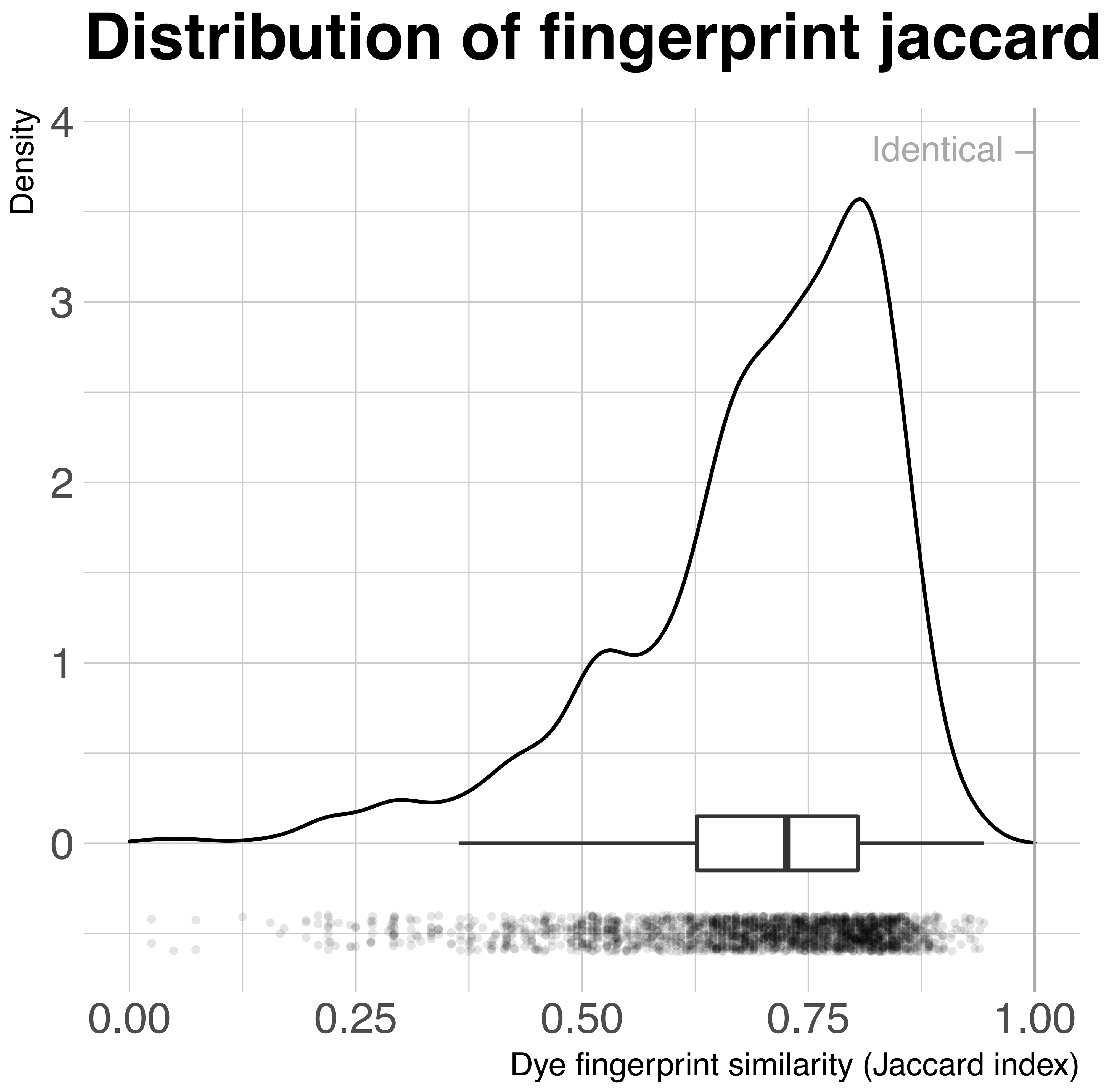


**Fig. S16. Dye fingerprints are unique to each protein.** Density plot of the pairwise Jaccard indices, which quantity the extent of overlaps in the assignments given to each dye across two different proteins. Identical dye fingerprints would have a jacquard value of 1. Box plot contains standard summary information (lower whisker = smallest observation greater than or equal to lower hinge - 1.5 * IQR, lower hinge = 25% quantile, median = 50% quantile, upper hinge = 75% quantile, upper whisker = largest observation less than or equal to upper hinge + 1.5 * IQR; outlier points not displayed). Jittered points: individual jaccard indices (2,211 points representing the 2,211 unique pairs between 67 proteins).

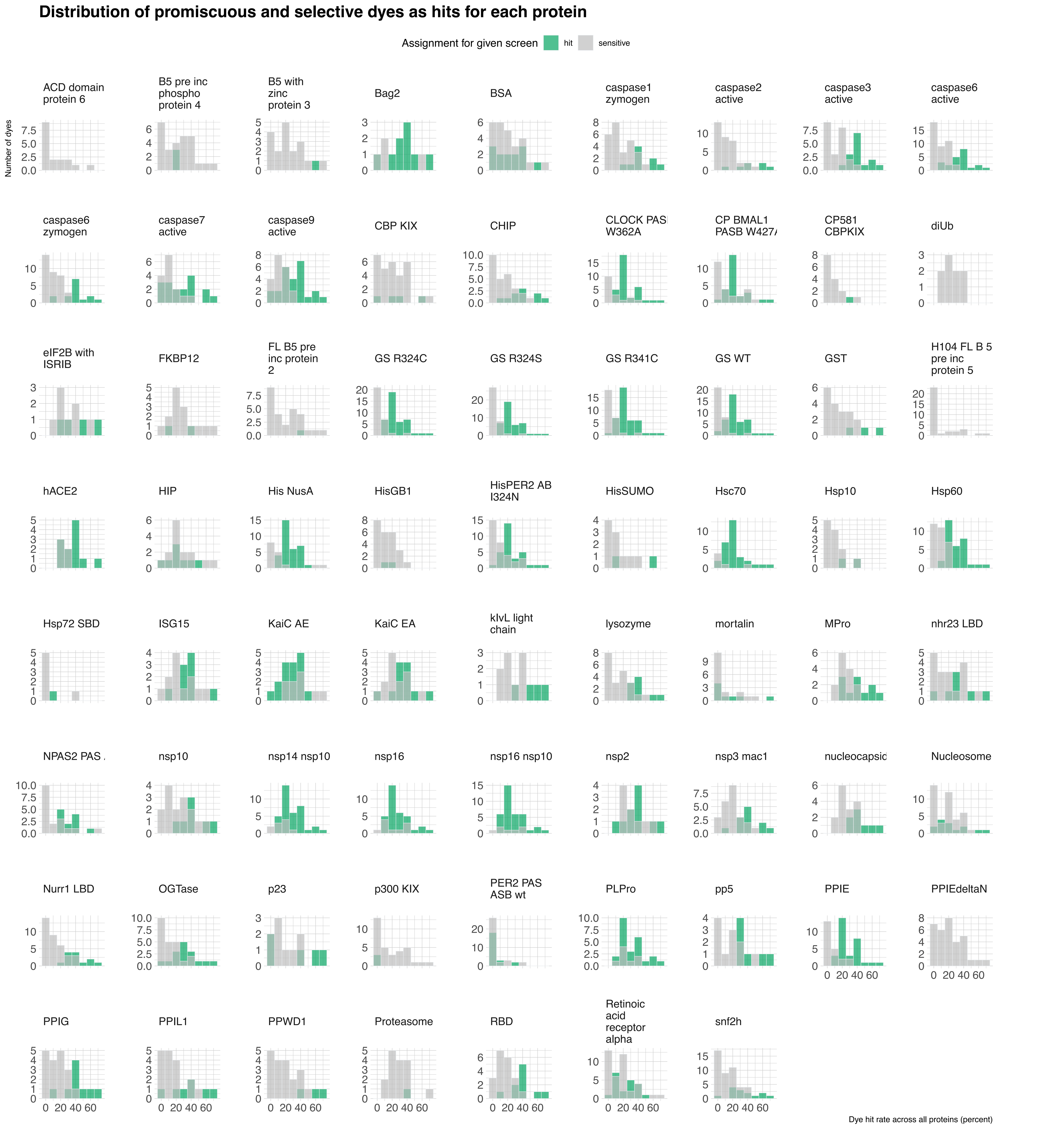


**Fig. S17. Distribution of promiscuous and selective dyes among the hit and sensitive dyes assigned for each protein.** Individual histograms displaying the number of hit dyes for a given protein, binned by the global hit rates of the specific hit dyes. Total hit rates are divided into ten evenly-spaced bins, containing hit rates of up to 10, 20, 30, 40, 50, 60, 70, 80, 90, or 100% of all tested proteins. Green bars: for "hit" assignment. Grey bars: for "sensitive" assignment. The presence of both selective and promiscuous dye hits in an individual dye screen appears as the presence of hits in both lower hit rate (e.g. < 20%) and higher hit rate (e.g. >40%) bins. Results for individual proteins are presented in individual panels, with the protein name indicated in the panel title.

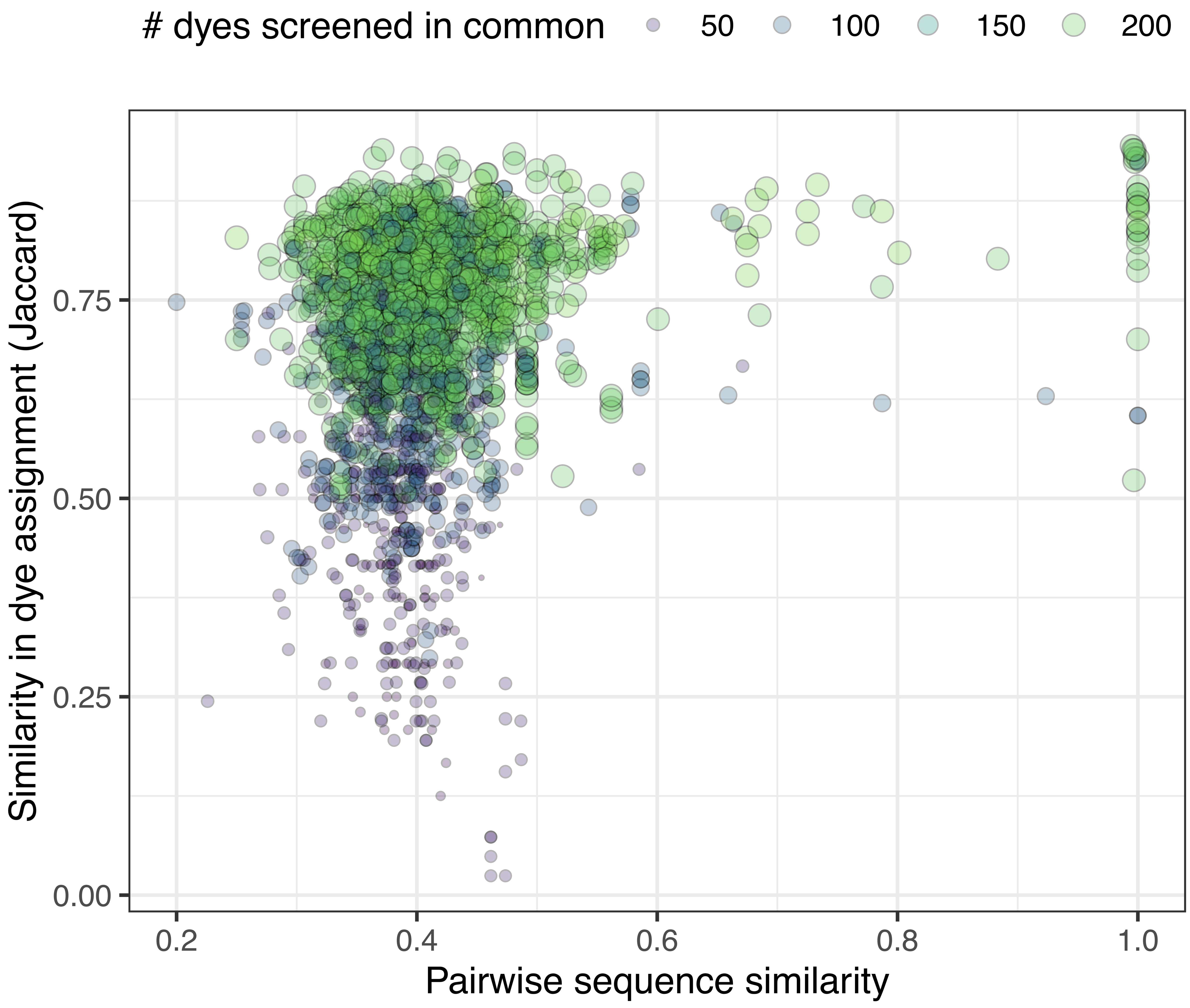


**Fig. S18. Weak positive correlation between protein sequence similarity and dye fingerprint.** Comparison of the relationship between primary sequence similarity and dye screen outcomes shows a weak positive correlation: 0.19, p-value 3.9e-20. Data shown includes 2,211 unique pairings between 67 proteins (72 proteins tested total - 5 proteins tested only as hetero-complexes; see Supplementary Methods). To provide context, the number of dyes screened in common between two proteins is indicated in point size and color. Smaller, darker points: fewer dyes screened in common. Larger, lighter points: more dyes screened in common. See figure key for exact mappings. Calculations of the Jaccard Index (y axis values) and pairwise sequence similarity (x axis values) are described in Supplementary Methods.

*See caption for complete list of dyes in Aurora concise*

Class | **Amyloid**
Dye | **A001**
Common name | **BF-188**

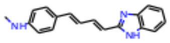

*In Aurora concise*

Class | **Amyloid**
Dye | **A002**
Common name | **PBB3**

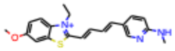

*In Aurora concise*

Class | **Amyloid**
Dye | **A003**
Common name | **FSB**

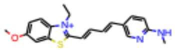

*In Aurora concise*

Class | **Amyloid**
Dye | **A004**
Common name | **BSB**

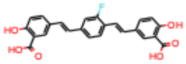

Class | **Amyloid**
Dye | **A005**
Common name | **K114**

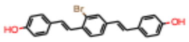

Class | **Amyloid**
Dye | **A006**
Common name | **Resorufin B.E.**

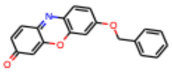

Class | **Amyloid**
Dye | **A007**
Common name | **Resorufin E.E.**

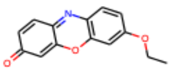

Class | **Amyloid**
Dye | **A008**
Common name | **Resorufin**

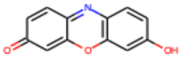

Class | **Amyloid**
Dye | **A009**
Common name | **Congo Red**

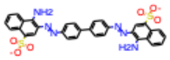

*In Aurora concise*

Class | **Amyloid**
Dye | **A010**
Common name | **NIAD4**

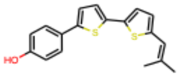

*In Aurora concise*

Class | **Amyloid**
Dye | **A011**
Common name | **Thiazine Red**

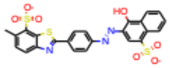

Class | **Amyloid**
Dye | **A012**
Common name | **Nile Red**

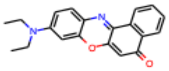

Class | **Amyloid**
Dye | **A013**
Common name | **LDS 698**

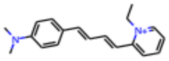

*In Aurora concise*

Class | **Amyloid**
Dye | **A014**
Common name | **LDS 722**

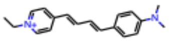

*In Aurora concise*

Class | **Amyloid**
Dye | **A015**
Common name | **LDS 730**

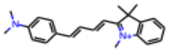

*In Aurora concise*

Class | **Amyloid**
Dye | **A016**
Common name | **PBB5 (aka Amb615160)**

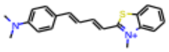

*In Aurora concise*

Class | **Amyloid**
Dye | **A017**
Common name | **CRANAD-2**

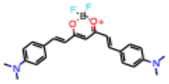

Class | **Amyloid**
Dye | **A018**
Common name | **THK 265**

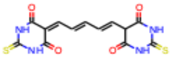

Class | **Amyloid**
Dye | **A019**
Common name | **BAP1**

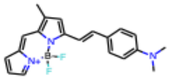

Class | **Amyloid**
Dye | **A020**
Common name | **ThT**

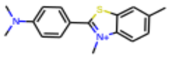

*In Aurora concise*

Class | **Amyloid**
Dye | **A021**
Common name | **DCM 6490**

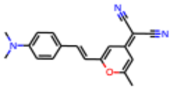

*In Aurora concise*

Class | **Amyloid**
Dye | **A022**
Common name | **LDS 798**

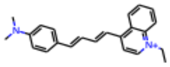

*In Aurora concise*

Class | **Amyloid**
Dye | **A023**
Common name | **Amb 1861079**

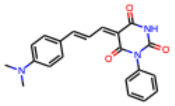

*In Aurora concise*

Class | **Amyloid**
Dye | **A024**
Common name | **Amb 2442356**

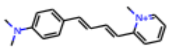

*In Aurora concise*

Class | **Amyloid**
Dye | **A025**
Common name | **Amb 1484600**

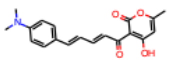

Class | **Common organic**
Dye | **C001**
Common name | **Meldola's Blue**

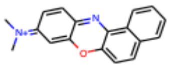

*In Aurora concise*

Class | **Common organic**
Dye | **C002**
Common name | **Xylene Cyanol FF**

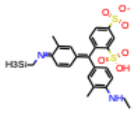

Class | **Common organic**
Dye | **C003**
Common name | **Victoria Blue R**

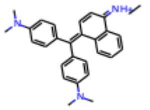

*In Aurora concise*

Class | **Common organic**
Dye | **C004**
Common name | **Acid Yellow 17**

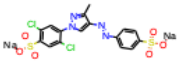

Class | **Common organic**
Dye | **C005**
Common name | **Sudan II**

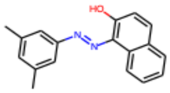

Class | **Common organic**
Dye | **C006**
Common name | **Naphthol Yellow S**

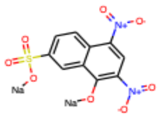

Class | **Common organic**
Dye | **C007**
Common name | **Naphthol Green B**

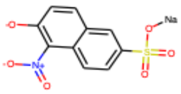

Class | **Common organic**
Dye | **C008**
Common name | **Fuchsin Acid**

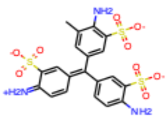

Class | **Common organic**
Dye | **C009**
Common name | **Rose Bengal**

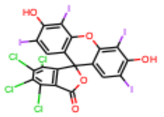

Class | **Common organic**
Dye | **C010**
Common name | **Victoria Blue B**

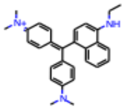

*In Aurora concise*

Class | **Common organic**
Dye | **C011**
Common name | **Azophloxine**

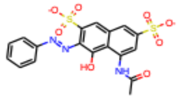

Class | **Common organic**
Dye | **C012**
Common name | **Orange G**

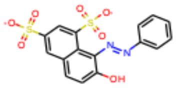

Class | **Common organic**
Dye | **C013**
Common name | **Chrysoidine G**

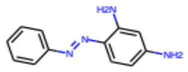

Class | **Common organic**
Dye | **C014**
Common name | **Bismark Brown Y**

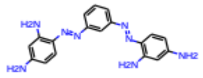

Class | **Common organic**
Dye | **C015**
Common name | **Janus Green B**

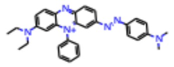

Class | **Common organic**
Dye | **C016**
Common name | **Martius Yellow**

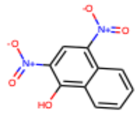

Class | **Common organic**
Dye | **C017**
Common name | **Tropaeolin O**

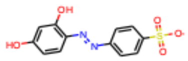

Class | **Common organic**
Dye | **C018**
Common name | **DCVJ**

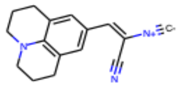

Class | **Common organic**
Dye | **C019**
Common name | **Titan Yellow**

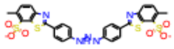

Class | **Common organic**
Dye | **C020**
Common name | **1,8,-ANS**

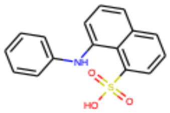

Class | **Common organic**
Dye | **C021**
Common name | **Amaranth Red**

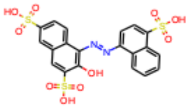

Class | **Common organic**
Dye | **C022**
Common name | **CPM**

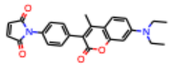

Class | **Laser**
Dye | **L001**
Common name | **LD390**

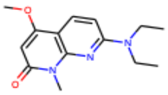

Class | **Laser**
Dye | **L002**
Common name | **POPOP**

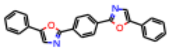

Class |
Dye | **LaserL003Bis-MSB**
Common name |

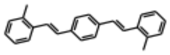

Class | **Laser**
Dye | **L004**
Common name | **LD423**

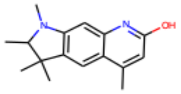

Class | **Laser**
Dye | **L005**
Common name | **LD425**

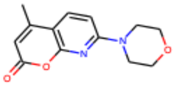

Class | **Laser**
Dye | **L006**
Common name | **Coumarin440**

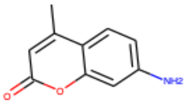

Class | **Laser**
Dye | **L007**
Common name | **Coumarin445**

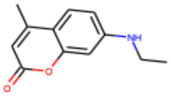

Class | **Laser**
Dye | **L008**
Common name | **Coumarin450**

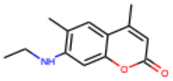

Class | **Laser**
Dye | **L009**
Common name | **Coumarin456**

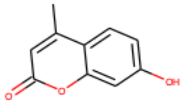

Class | **Laser**
Dye | **L010**
Common name | **Coumarin460**

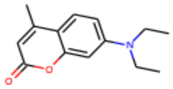

Class | **Laser**
Dye | **L011**
Common name | **Coumarin461**

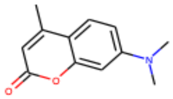

Class | **Laser**
Dye | **L012**
Common name | **LD466**

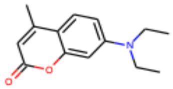

Class | **Laser**
Dye | **L013**
Common name | **LD473**

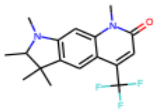

Class | **Laser**
Dye | **L014**
Common name | **Coumarin478**

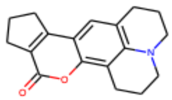

Class | **Laser**
Dye | **L015**
Common name | **Coumarin480**

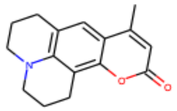

Class | **Laser**
Dye | **L016**
Common name | **Coumarin481**

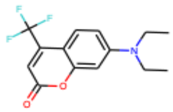

Class | **Laser**
Dye | **L017**
Common name | **Coumarin485**

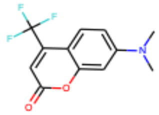

Class | **Laser**
Dye | **L018**
Common name | **LD489**

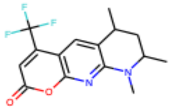

Class | **Laser**
Dye | **L019**
Common name | **Coumarin490**

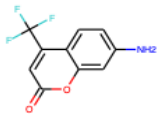

Class | **Laser**
Dye | **L020**
Common name | **LD490**

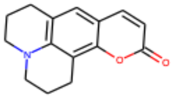

Class | **Laser**
Dye | **L021**
Common name | **Coumarin498**

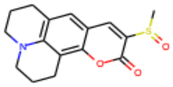

Class | **Laser**
Dye | **L022**
Common name | **Coumarin500**

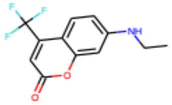

Class | **Laser**
Dye | **L023**
Common name | **Coumarin503**

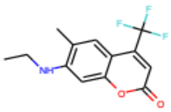

Class | **Laser**
Dye | **L024**
Common name | **Coumarin504**

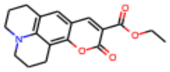

Class | **Laser**
Dye | **L025**
Common name | **Coumarin510**

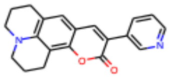

Class | **Laser**
Dye | **L026**
Common name | **Coumarin515**

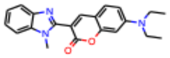

Class | **Laser**
Dye | **L027**
Common name | **Coumarin519**

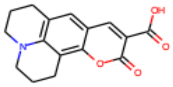

Class | **Laser**
Dye | **L028**
Common name | **Coumarin521**

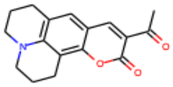

Class | **Laser**
Dye | **L029**
Common name | **Coumarin523**

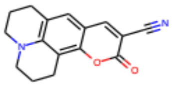

Class | **Laser**
Dye | **L030**
Common name | **Coumarin525**

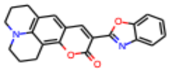

Class | **Laser**
Dye | **L031**
Common name | **Coumarin535**

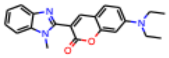

Class | **Laser**
Dye | **L032**
Common name | **Coumarin540**

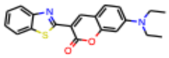

Class | **Laser**
Dye | **L033**
Common name | **Coumarin540A**

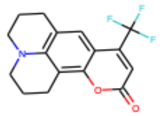

Class | **Laser**
Dye | **L034**
Common name | **Coumarin545**

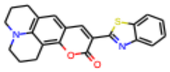

Class | **Laser**
Dye | **L035**
Common name | **Pyrromethene546**

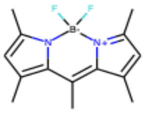

Class | **Laser**
Dye | **L036**
Common name | **Pyrromethene556**

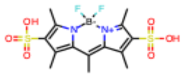

Class | **Laser**
Dye | **L037**
Common name | **Pyrromethene567**

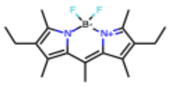

Class | **Laser**
Dye | **L038**
Common name | **Pyrromethene580**

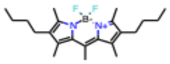

Class | **Laser**
Dye | **L039**
Common name | **Pyrromethene597**

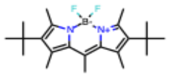

Class | **Laser**
Dye | **L040**
Common name | **Pyrromethene597-8C9**

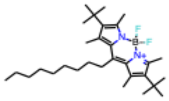

Class | **Laser**
Dye | **L041**
Common name | **Pyrromethene605**

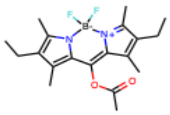

Class | **Laser**
Dye | **L042**
Common name | **Pyrromethene650**

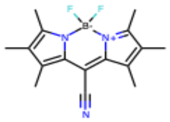

Class | **Laser**
Dye | **L043**
Common name | **Fluorescein548**

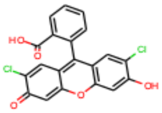

Class | **Laser**
Dye | **L044**
Common name | **DisodiumFluorescein**

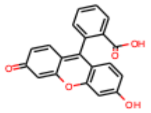

Class | **Laser**
Dye | **L045**
Common name | **Fluorol555**

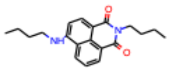

Class | **Laser**
Dye | **L046**
Common name | **Rhodamine3BPerchlorate**

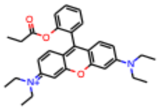

Class | **Laser**
Dye | **L047**
Common name | **Rhodamine560Chloride**

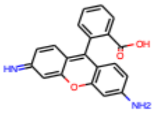

Class | **Laser**
Dye | **L048**
Common name | **Rhodamine560Perchlorate**

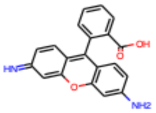

Class | **Laser**
Dye | **L049**
Common name | **Rhodamine575**

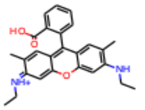

Class | **Laser**
Dye | **L050**
Common name | **Rhodamine19Perchlorate**

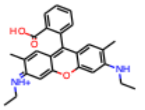

Class | **Laser**
Dye | **L051**
Common name | **Rhodamine590Chloride**

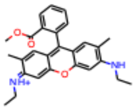

Class | **Laser**
Dye | **L052**
Common name | **Rhodamine590Tetrafluoro borate**

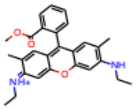

Class | **Laser**
Dye | **L053**
Common name | **Rhodamine590Perchlorate**

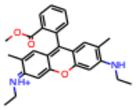

Class | **Laser**
Dye | **L054**
Common name | **Rhodamine610Chloride**

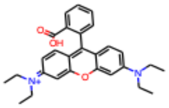

Class | **Laser**
Dye | **L055**
Common name | **Rhodamine610Perchlorate**

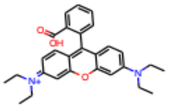

Class | **Laser**
Dye | **L056**
Common name | **Rhodamine610Tetrafluoro borate**

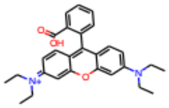

Class | **Laser**
Dye | **L057**
Common name | **KitonRed620**

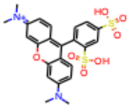

Class | **Laser**
Dye | **L058**
Common name | **Rhodamine640Perchlorate**

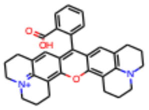

Class | **Laser**
Dye | **L059**
Common name | **Sulforhodamine640**

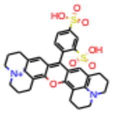

Class | **Laser**
Dye | **L060**
Common name | **DODClodide**

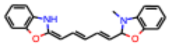

*In Aurora concise*

Class | **Laser**
Dye | **L061**
Common name | **LD688**

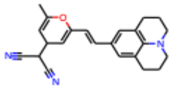

Class | **Laser**
Dye | **L062**
Common name | **LDS750**

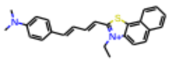

*In Aurora concise*

Class | **Laser**
Dye | **L063**
Common name | **LDS759**

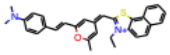

*In Aurora concise*

Dyes in Aurora concise:,,,

L060 , L062, L063, L064 ,L089, L090 , L091,L093,L094, L095 ,L096,  L097, , L098, ,  L099

Class | **Laser**
Dye | **L064**
Common name | **LDS821**

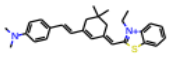

*In Aurora concise*

Class | **Laser**
Dye | **L065**
Common name | **LDS925**

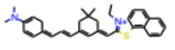

Class | **Laser**
Dye | **L066**
Common name | **LDS950**

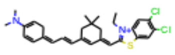

Class | **Laser**
Dye | **L067**
Common name | **LD800**

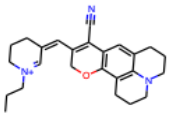

Class | **Laser**
Dye | **L068**
Common name | **Phenoxazone660**

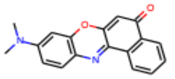

Class | **Laser**
Dye | **L069**
Common name | **CresylViolet670Perchlorate**

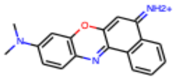

Class | **Laser**
Dye | **L070**
Common name | **NileBlue690Perchlorate**

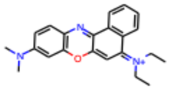

Class | **Laser**
Dye | **L071**
Common name | **LD690Perchlorate**

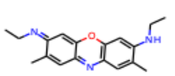

Class | **Laser**
Dye | **L072**
Common name | **LD700Perchlorate**

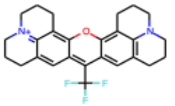

Class | **Laser**
Dye | **L073**
Common name | **Oxazine720Perchlorate**

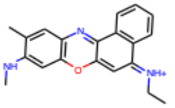

Class | **Laser**
Dye | **L074**
Common name | **Oxazine725Perchlorate**

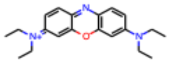

Class | **Laser**
Dye | **L075**
Common name | **Oxazine725Chloride**

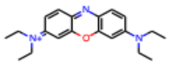

Class | **Laser**
Dye | **L076**
Common name | **Oxazine750Perchlorate**

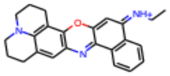

Class | **Laser**
Dye | **L077**
Common name | **HIDClodide**

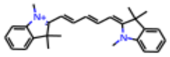

Class | **Laser**
Dye | **L078**
Common name | **DOTClodide**

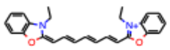

Class | **Laser**
Dye | **L079**
Common name | **DOTCPerchlorate**

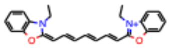

Class | **Laser**
Dye | **L080**
Common name | **HITCPerchlorate**

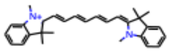

Class | **Laser**
Dye | **L081**
Common name | **HITClodide**

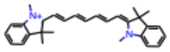

Class | **Laser**
Dye | **L082**
Common name | **HITCSpecial**

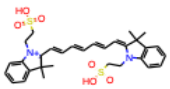

Class | **Laser**
Dye | **L083**
Common name | **DTTClodide**

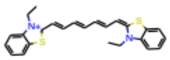

Class | **Laser**
Dye | **L084**
Common name | **IR-26**

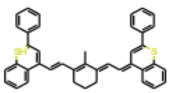

Class | **Laser**
Dye | **L085**
Common name | **IR-125**

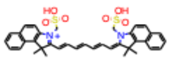

Class | **Laser**
Dye | **L097**
Common name | **DQTCI**

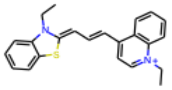

*In Aurora concise*

Class | **Laser**
Dye | **L098**
Common name | **DQTCI(')**

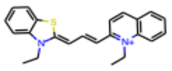

*In Aurora concise*

Class | **Laser**
Dye | **L099**
Common name | **DTCI**

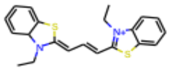

*In Aurora concise*

Class | **Laser**
Dye | **L100**
Common name | **Cryptocyanine**

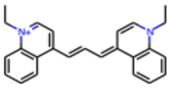

Class | **Laser**
Dye | **L101**
Common name | **DTDCI**

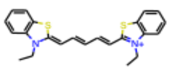

Class | **Laser**
Dye | **L102**
Common name | **NCI**

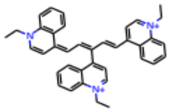

Class | **Laser**
Dye | **L103**
Common name | **HDITCP**

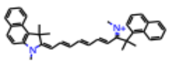

Class | **Laser**
Dye | **L104**
Common name | **DQTrCI**

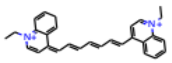

Class | **Laser**
Dye | **L105**
Common name | **Q-SwitchI(Kodak9740)**

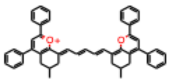

Class | **Laser**
Dye | **L106**
Common name | **Q-Switch5**

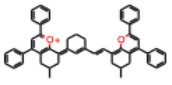

Class | **Laser**
Dye | **L107**
Common name | **MalachiteGreen**

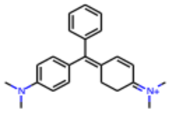

Class | **Laser**
Dye | **L086**
Common name | **IR-140**

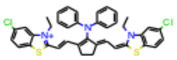

Class | **Laser**
Dye | **L087**
Common name | **IR-143**

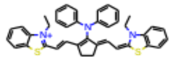

Class | **Laser**
Dye | **L088**
Common name | **IR-144**

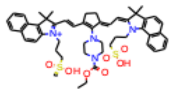

Class | **Laser**
Dye | **L089**
Common name | **DASPI**

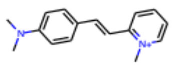

*In Aurora concise*

Class | **Laser**
Dye | **L090**
Common name | **PICI**

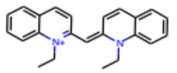

*In Aurora concise*

Class | **Laser**
Dye | **L091**
Common name | **DMETCI**

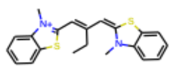

*In Aurora concise*

Class | **Laser**
Dye | **L092**
Common name | **DOCI**

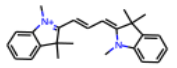

Class | **Laser**
Dye | **L093**
Common name | **HICI**

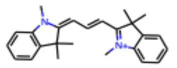

*In Aurora concise*

Class | **Laser**
Dye | **L094**
Common name | **DQOCI**

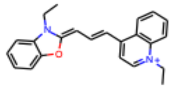

*In Aurora concise*

Class | **Laser**
Dye | **L095**
Common name | **Pinacyanol**

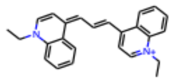

*In Aurora concise*

Class | **Laser**
Dye | **L096**
Common name | **DDBCI**

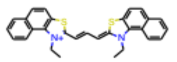

*In Aurora concise*

Class | **Laser**
Dye | **L108**
Common name | **BDN**

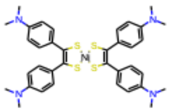

Class | **Max Weaver Set 1**
Dye | **MWA001**
Common name | **MWA001**

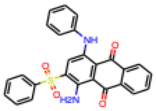

Class | **Max Weaver Set 1**
Dye | **MWA002**
Common name | **MWA002**

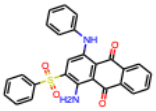

Class | **Max Weaver Set 1**
Dye | **MWA003**
Common name | **MWA003**

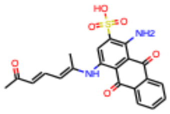

Class | **Max Weaver Set 1**
Dye | **MWA004**
Common name | **MWA004**

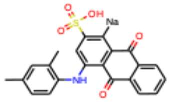

Class | **Max Weaver Set 1**
Dye | **MWA005**
Common name | **MWA005**

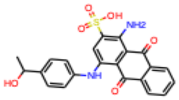

Class | **Max Weaver Set 1**
Dye | **MWA006**
Common name | **MWA006**

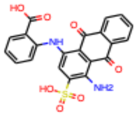

Class | **Max Weaver Set 1**
Dye | **MWA007**
Common name | **MWA007**

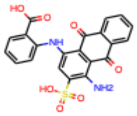

Class | **Max Weaver Set 1**
Dye | **MWA008**
Common name | **MWA008**

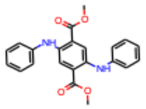

Class | **Max Weaver Set 1**
Dye | **MWA010**
Common name | **MWA010**

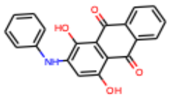

Class | **Max Weaver Set 1**
Dye | **MWA011**
Common name | **MWA011**

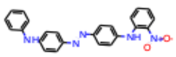

Class | **Max Weaver Set 1**
Dye | **MWA013**
Common name | **MWA013**

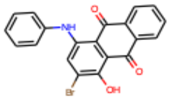

Class | **Max Weaver Set 1**
Dye | **MWA014**
Common name | **MWA014**

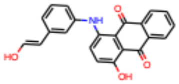

Class | **Max Weaver Set 1**
Dye | **MWA015**
Common name | **MWA015**

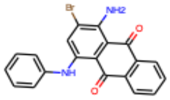

Class | **Max Weaver Set 1**
Dye | **MWA016**
Common name | **MWA016**

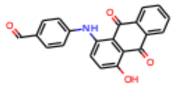

Class | **Max Weaver Set 1**
Dye | **MWA017**
Common name | **MWA017**

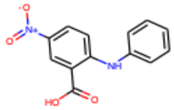

Class | **Max Weaver Set 1**
Dye | **MWA018**
Common name | **MWA018**

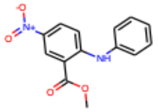

Class | **Max Weaver Set 1**
Dye | **MWA020**
Common name | **MWA020**

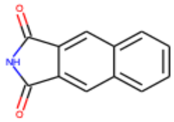

Class | **Max Weaver Set 1**
Dye | **MWA021**
Common name | **MWA021**

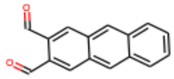

Class | **Max Weaver Set 2**
Dye | **MWB001**
Common name | **MWB001**

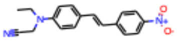

Class | **Max Weaver Set 2**
Dye | **MWB002**
Common name | **MWB002**

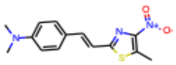

Class | **Max Weaver Set 2**
Dye | **MWB003**
Common name | **MWB003**

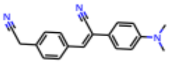

Class | **Max Weaver Set 2**
Dye | **MWB004**
Common name | **MWB004**

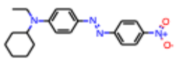

Class | **Max Weaver Set 2**
Dye | **MWB005**
Common name | **MWB005**

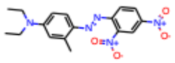

Class | **Max Weaver Set 2**
Dye | **MWB006**
Common name | **MWB006**

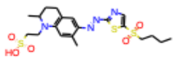

Class | **Max Weaver Set 2**
Dye | **MWB007**
Common name | **MWB007**

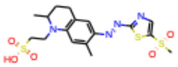

*Dye MWB008 not received/tested*

Class | **Max Weaver Set 2**
Dye | **MWB009**
Common name | **MWB009**

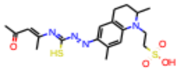

*In Aurora concise*

*Dye MWB011 not received/tested*

Class | **Max Weaver Set 2**
Dye | **MWB011**
Common name | **MWB011**

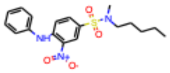

Class | **Max Weaver Set 2**
Dye | **MWB012**
Common name | **MWB012**

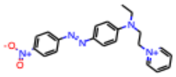

Class | **Max Weaver Set 2**
Dye | **MWB013**
Common name | **MWB013**

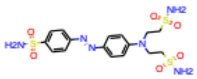

Class | **Max Weaver Set 2**
Dye | **MWB014**
Common name | **MWB014**

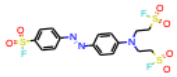

Class | **Max Weaver Set 2**
Dye | **MWB015**
Common name | **MWB015**

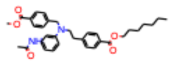

Class | **Max Weaver Set 2**
Dye | **MWB016**
Common name | **MWB016**

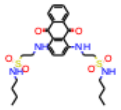

Class | **Max Weaver Set 2**
Dye | **MWB017**
Common name | **MWB017**

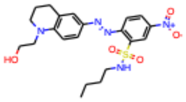

Class | **Max Weaver Set 2**
Dye | **MWB019**
Common name | **MWB019**

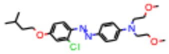

Class | **Max Weaver Set 3**
Dye | **MWC001**
Common name | **MWC001**

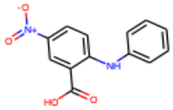

Class | **Max Weaver Set 3**
Dye | **MWC002**
Common name | **MWC002**

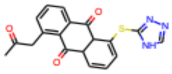

Class | **Max Weaver Set 3**
Dye | **MWC003**
Common name | **MWC003**

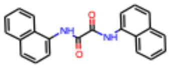

Class | **Max Weaver Set 3**
Dye | **MWC004**
Common name | **MWC004**

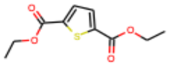

Class | **Max Weaver Set 3**
Dye | **MWC005**
Common name | **MWC005**

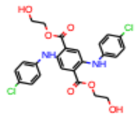

Class | **Max Weaver Set 3**
Dye | **MWC006**
Common name | **MWC006**

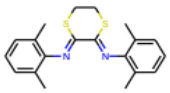

Class | **Max Weaver Set 3**
Dye | **MWC007**
Common name | **MWC007**

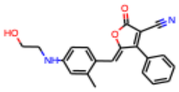

*In Aurora concise*

Class | **Max Weaver Set 3**
Dye | **MWC008**
Common name | **MWC008**

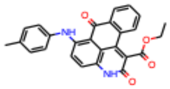

*In Aurora concise*

Class | **Max Weaver Set 3**
Dye | **MWC009**
Common name | **MWC009**

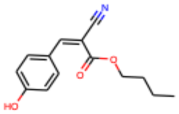

Class | **Max Weaver Set 3**
Dye | **MWC010**
Common name | **MWC010**

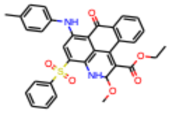

Class | **Max Weaver Set 3**
Dye | **MWC011**
Common name | **MWC011**

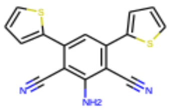

Class | **Max Weaver Set 3**
Dye | **MWC012**
Common name | **MWC012**

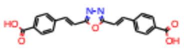

Class | **Max Weaver Set 3**
Dye | **MWC013**
Common name | **MWC013**

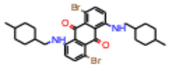

Class | **Max Weaver Set 3**
Dye | **MWC014**
Common name | **MWC014**

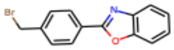

Class | **Max Weaver Set 3**
Dye | **MWC015**
Common name | **MWC015**

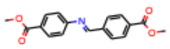

Class | **Max Weaver Set 3**
Dye | **MWC016**
Common name | **MWC016**

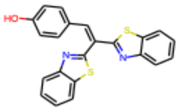

Class | **Max Weaver Set 3**
Dye | **MWC017**
Common name | **MWC017**

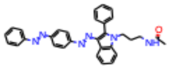

Class | **Max Weaver Set 3**
Dye | **MWC018**
Common name | **MWC018**

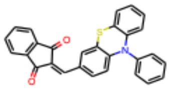

Class | **Max Weaver Set 3**
Dye | **MWC019**
Common name | **MWC019**

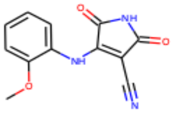

Class | **Max Weaver Set 3**
Dye | **MWC020**
Common name | **MWC020**

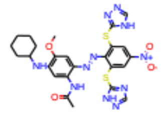

Class | **Max Weaver Set 3**
Dye | **MWC021**
Common name | **MWC021**

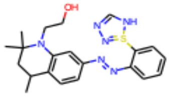

*In Aurora concise*

Class | **Max Weaver Set 3**
Dye | **MWC022**
Common name | **MWC022**

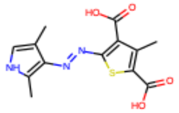

Class | **Max Weaver Set 3**
Dye | **MWC023**
Common name | **MWC023**

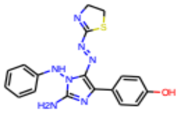

Class | **Max Weaver Set 3**
Dye | **MWC024**
Common name | **MWC024**

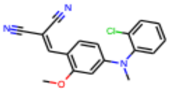

Class | **Max Weaver Set 3**
Dye | **MWC025**
Common name | **MWC025**

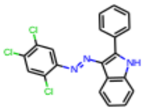

Class | **Max Weaver Set 3**
Dye | **MWC026**
Common name | **MWC026**

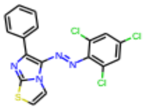

Class | **Max Weaver Set 3**
Dye | **MWC027**
Common name | **MWC027**

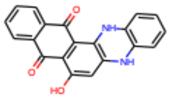

Class | **Max Weaver Set 3**
Dye | **MWC028**
Common name | **MWC028**

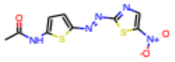

Class | **Max Weaver Set 3**
Dye | **MWC029**
Common name | **MWC029**

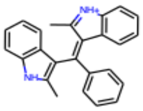

Class | **Max Weaver Set 3**
Dye | **MWC030**
Common name | **MWC030**

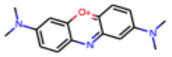

Class | **Max Weaver Set 3**
Dye | **MWC031**
Common name | **MWC031**

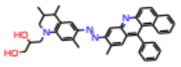

Class | **Max Weaver Set 3**
Dye | **MWC032**
Common name | **MWC032**

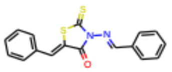

Class | **Max Weaver Set 3**
Dye | **MWC033**
Common name | **MWC033**

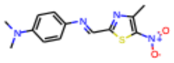

Class | **Max Weaver Set 3**
Dye | **MWC034**
Common name | **MWC034**

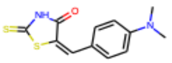

*In Aurora concise*

Class | **Max Weaver Set 3**
Dye | **MWC035**
Common name | **MWC035**

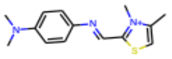

Class | **Max Weaver Set 3**
Dye | **MWC036**
Common name | **MWC036**

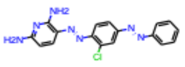

Class | **Max Weaver Set 3**
Dye | **MWC037**
Common name | **MWC037**

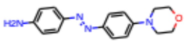

Class | **Max Weaver Set 3**
Dye | **MWC038**
Common name | **MWC038**

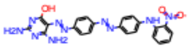

Class | **Max Weaver Set 3**
Dye | **MWC039**
Common name | **MWC039**

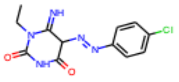

Class | **Max Weaver Set 3**
Dye | **MWC040**
Common name | **MWC040**

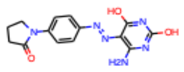

Class | **Max Weaver Set 3**
Dye | **MWC041**
Common name | **MWC041**

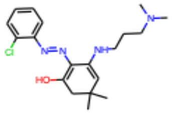

Class | **Max Weaver Set 3**
Dye | **MWC042**
Common name | **MWC042**

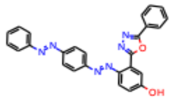

Class | **Max Weaver Set 3**
Dye | **MWC043**
Common name | **MWC043**

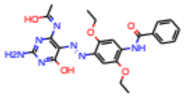

Class | **Max Weaver Set 3**
Dye | **MWC044**
Common name | **MWC044**

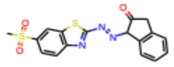

Class | **Max Weaver Set 3**
Dye | **MWC045**
Common name | **MWC045**

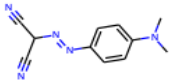

*In Aurora concise*

Class | **Max Weaver Set 3**
Dye | **MWC046**
Common name | **MWC046**

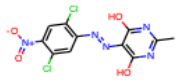

Class | **Max Weaver Set 3**
Dye | **MWC047**
Common name | **MWC047**

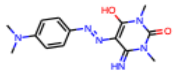

Class | **Max Weaver Set 3**
Dye | **MWC048**
Common name | **MWC048**

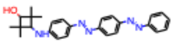

Class | **Max Weaver Set 3**
Dye | **MWC049**
Common name | **MWC049**

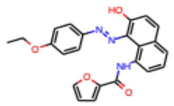

Class | **Max Weaver Set 3**
Dye | **MWC050**
Common name | **MWC050**

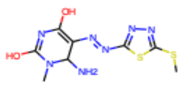

Class | **Max Weaver Set 3**
Dye | **MWC051**
Common name | **MWC051**

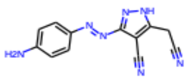

Class | **Max Weaver Set 3**
Dye | **MWC052**
Common name | **MWC052**

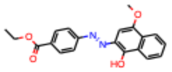

Class | **Max Weaver Set 3**
Dye | **MWC053**
Common name | **MWC053**

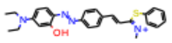

*In Aurora concise*

Class | **Max Weaver Set 3**
Dye | **MWC054**
Common name | **MWC054**

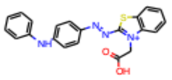

Class | **Max Weaver Set 3**
Dye | **MWC055**
Common name | **MWC055**

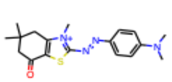

Class | **Max Weaver Set 3**
Dye | **MWC056**
Common name | **MWC056**

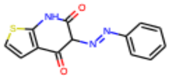

Class | **Max Weaver Set 3**
Dye | **MWC057**
Common name | **MWC057**

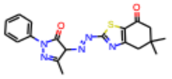

Class | **Max Weaver Set 3**
Dye | **MWC058**
Common name | **MWC058**

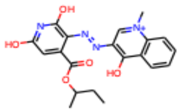

Class | **Max Weaver Set 3**
Dye | **MWC059**
Common name | **MWC059**

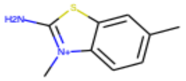

Class | **Max Weaver Set 3**
Dye | **MWC060**
Common name | **MWC060**

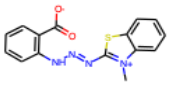

Class | **Max Weaver Set 3**
Dye | **MWC061**
Common name | **MWC061**

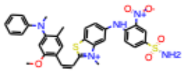

*In Aurora concise*

Class | **Max Weaver Set 3**
Dye | **MWC062**
Common name | **MWC062**

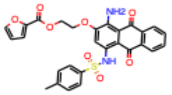

Class | **Max Weaver Set 4**
Dye | **MWD01**
Common name | **MWD01**

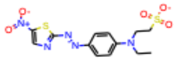

Class | **Max Weaver Set 4**
Dye | **MWD02**
Common name | **MWD02**

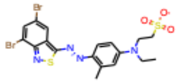

Class | **Max Weaver Set 4**
Dye | **MWD03**
Common name | **MWD03**

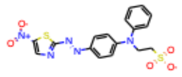

Class | **Max Weaver Set 4**
Dye | **MWD04**
Common name | **MWD04**

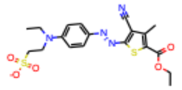

Class | **Max Weaver Set 4**
Dye | **MWD05**
Common name | **MWD05**

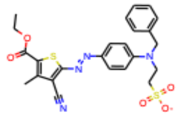

Class | **Max Weaver Set 4**
Dye | **MWD06**
Common name | **MWD06**

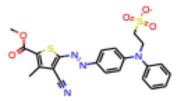

Class | **Max Weaver Set 4**
Dye | **MWD07**
Common name | **MWD07**

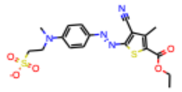

Class | **Max Weaver Set 4**
Dye | **MWD08**
Common name | **MWD08**

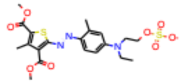

Class | **Max Weaver Set 4**
Dye | **MWD09**
Common name | **MWD09**

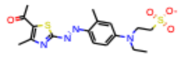

Class | **Max Weaver Set 4**
Dye | **MWD10**
Common name | **MWD10**

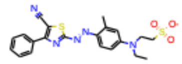

Class | **Max Weaver Set 4**
Dye | **MWD11**
Common name | **MWD11**

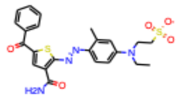

Class | **Max Weaver Set 4**
Dye | **MWD12**
Common name | **MWD12**

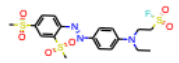

Class | **Max Weaver Set 4**
Dye | **MWD13**
Common name | **MWD13**

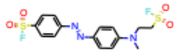

Class | **Max Weaver Set 4**
Dye | **MWD14**
Common name | **MWD14**

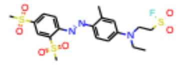

Class | **Max Weaver Set 4**
Dye | **MWD15**
Common name | **MWD15**

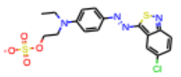

Class | **Max Weaver Set 4**
Dye | **MWD16**
Common name | **MWD16**

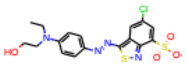

Class | **Max Weaver Set 4**
Dye | **MWD17**
Common name | **MWD17**

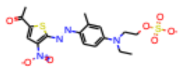

Class | **Max Weaver Set 5**
Dye | **MWE01**
Common name | **MWE01**

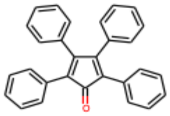

Class | **Max Weaver Set 5**
Dye | **MWE02**
Common name | **MWE02**

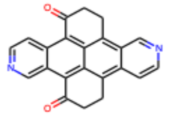

Class | **Max Weaver Set 5**
Dye | **MWE03**
Common name | **MWE03**

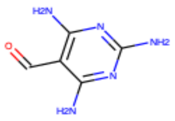

Class | **Max Weaver Set 5**
Dye | **MWE04**
Common name | **MWE04**

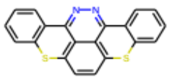

Class | **Max Weaver Set 5**
Dye | **MWE05**
Common name | **MWE05**

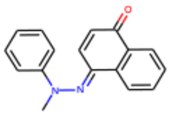

Class | **Max Weaver Set 5**
Dye | **MWE06**
Common name | **MWE06**

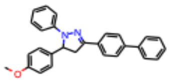

Class | **Max Weaver Set 5**
Dye | **MWE07**
Common name | **MWE07**

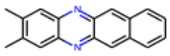

Class | **Max Weaver Set 5**
Dye | **MWE08**
Common name | **MWE08**

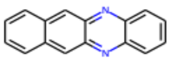

Class | **Max Weaver Set 5**
Dye | **MWE09**
Common name | **MWE09**

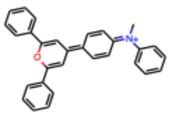

Class | **Max Weaver Set 5**
Dye | **MWE10**
Common name | **MWE10**

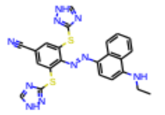

Class | **Max Weaver Set 5**
Dye | **MWE11**
Common name | **MWE11**

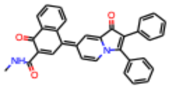

Class | **Max Weaver Set 5**
Dye | **MWE12**
Common name | **MWE12**

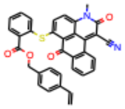

Class | **Max Weaver Set 5**
Dye | **MWE13**
Common name | **MWE13**

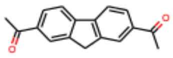

Class | **Max Weaver Set 5**
Dye | **MWE14**
Common name | **MWE14**

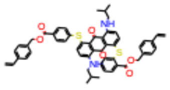

Class | **Max Weaver Set 6**
Dye | **MWF01**
Common name | **MWF01**

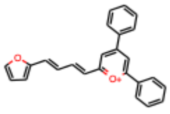

Class | **Max Weaver Set 6**
Dye | **MWF02**
Common name | **MWF02**

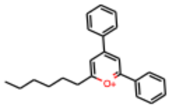

Class | **Max Weaver Set 6**
Dye | **MWF03**
Common name | **MWF03**

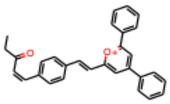

Class | **Max Weaver Set 6**
Dye | **MWF04**
Common name | **MWF04**

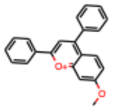

Class | **Max Weaver Set 6**
Dye | **MWF05**
Common name | **MWF05**

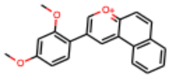

Class | **Max Weaver Set 6**
Dye | **MWF06**
Common name | **MWF06**

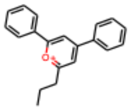

Class | **Max Weaver Set 6**
Dye | **MWF07**
Common name | **MWF07**

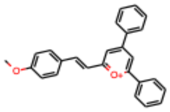

Class | **Max Weaver Set 6**
Dye | **MWF08**
Common name | **MWF08**

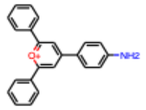

*In Aurora concise*

Class | **SYPRO**
Dye | **SYPRO**
Common name | **SYPRO Orange**

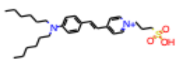

*In Aurora concise*

Class | **Commerial textile, order 1**
Dye | **T001**
Common name | **Direct Brown 210**

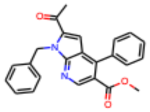

Class | **Commerial textile, order 1**
Dye | **T002**
Common name | **Direct Yellow 96**

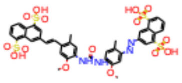

Class | **Commerial textile, order 1**
Dye | **T004**
Common name | **Quinoline Yellow WS**

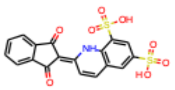

*In Aurora concise*

Class | **Commerial textile, order 1**
Dye | **T008**
Common name | **Direct Red 227**

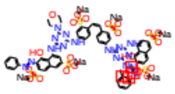

Class | **Commerial textile, order 1**
Dye | **T009**
Common name | **Direct Red 224**

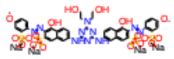

Class | **Commerial textile, order 1**
Dye | **T010**
Common name | **Direct Red 23**

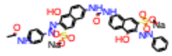

Class | **Commerial textile, order 1**
Dye | **T011**
Common name | **Direct Red 81**

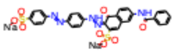

Class | **Commerial textile, order 1**
Dye | **T014**
Common name | **Direct Violet 51**

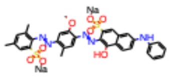

Class | **Commerial textile, order 1**
Dye | **T016**
Common name | **Direct Blue 199**

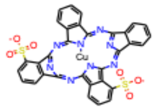

Class | **Commerial textile, order 1**
Dye | **T027**
Common name | **Direct Black 80**

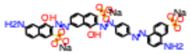

Class | **SYPROlogs**
Dye | **TW383**
Common name | **SYPROlog1**

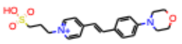

Class | **SYPROlogs**
Dye | **TW384***
Common name | **SYPROlog2**

*synthesized in-house (not commercially available)*

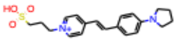

*In Aurora concise*

Class | **SYPROlogs**
Dye | **TW385***
Common name | **SYPROlog3**

*synthesized in-house (not commercially available)*

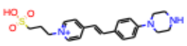

Class | **SYPROlogs**
Dye | **TW408***
Common name | **SYPROlog4**

*synthesized in-house (not commercially available)*

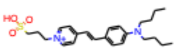

*In Aurora concise*

Class | **SYPROlogs**
Dye | **TW418***
Common name | **SYPROlog5**

*synthesized in-house (not commercially available)*

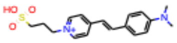

*In Aurora concise*

Class | **SYPROlogs**
Dye | **TW419***
Common name | **SYPROlog6**
**synthesized in-house (not commercially available)*

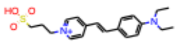

*In Aurora concise*

Class | **SYPROlogs**
Dye | **TW495***
Common name | **SYPROlog7**
**synthesized in-house (not commercially available)*

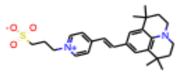

*In Aurora concise*

Class | **SYPROlogs**
Dye | **AW131***
Common name | **SYPROlog8**
**synthesized in-house (not commercially available)*

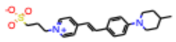

*In Aurora concise*

Class | **SYPROlogs**
Dye | **AW134***
Common name | **SYPROlog9**
**synthesized in-house (not commercially available)*

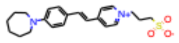

*In Aurora concise*

Class | **SYPROlogs**
Dye | **AW132***
Common name | **SYPROlog10**
**synthesized in-house (not commercially available)*

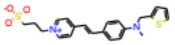

**Table S1. Chemical structures and common names for all Aurora library dyes.** Chemical structures, Aurora library name, and common name (where applicable) for all 312 tested dyes in the Aurora library. Unless otherwise noted, dyes are available either commercially, or through the North Carolina State Max Weaver Dye Library. Detailed information on dye sources is available in Data S1.

Complete list of dyes in Aurora concise: TW408, MWF08, TW495, MWC061, L063, L096, A022, A016, T004, L091, L097, L093, A014, L095, L098, L094, SYPRO Orange, A003, L062, A024, MWC007, A002, A020, L060, A001 , L089, L099, TW384, A013, MWC053, TW419, A009, A015, C003, L064, A021, MWC034, AW134, MWC045, C010, L090, A023, MWC021, MWC008, A010, AW131, MWB009, C001, TW418

| Protein | | | | Dye screen conditions | | Biochemical / structural properties | | | | | | | Source / references | | | |
|---|---|---|---|---|---|---|---|---|---|---|---|---|---|---|---|---|
| Abbreviated protein name | Full protein name | Construct details | Annotation | Screening conc. (µM) | Screening buffer | Sequence Length | Molecular Weight | GRAVY hydrophobicity | Percent disordered | Fraction Helix | Fraction Turn | Fraction Sheet | Protein source | Uniprot ID | PDB Code | Ref. |
| ACD_domain_protein_6 | Alpha-crystallin B chain | Alpha crystallin domain of HSPB5 | Eye lens refractivity, chaperone-like properties | 10 | 25 mM MOPS, 150 mM NaCl, pH 7.50 | 89 | 10180.3614 | -0.652809 | 10 | 0.30337079 | 0.2247191 | 0.21348315 | Maria Janowska, Klevit Lab, UW | P02511 | | Rajagopal, Ponni, et al. 2015 |
| B5_pre_inc_phospho_protein_4 | Alpha-crystallin B chain | phosphomimetic construct, incubated at 37C to activate | Oligomeric/conformational modulation of HSPB5 | 10 | 25 mM MOPS, 150 mM NaCl, pH 7.50 | 175 | 20242.6725 | -0.5777143 | 21 | 0.30285714 | 0.23428571 | 0.21142857 | Maria Janowska, Klevit Lab, UW | P02511 | | Rajagopal, Ponni, et al. 2015 |
| B5_with_zinc_protein_3 | Alpha-crystallin B chain | HSPB5 in the presence of Zinc | Oligomeric/conformational modulation of HSPB5 | 10 | 25 mM MOPS, 150 mM NaCl, pH 7.50 | 175 | 20158.6422 | -0.5314286 | 21 | 0.30285714 | 0.25142857 | 0.21142857 | Maria Janowska, Klevit Lab, UW | P02511 | | Rajagopal, Ponni, et al. 2015 |
| Bag2 | BAG family molecular chaperone regulator 2 | | Co-chaperone, nucletide-exchange factor | 5 | 10 mM HEPES, pH 7.20, 200 mM NaCl | 211 | 23771.6663 | -0.5919431 | 22 | 0.2464455 | 0.20853081 | 0.32701422 | Rebecca Freilich, Gestwicki Lab, UCSF | O95816 | | |
| BSA | Bovine Serum Albumin | | Blood osmotic pressure regulation; ion and small molecule binding | 10 | 20 mM HEPES, pH 7.5, 150 mM KCl, 20 mM NaCl, 5 mM MgCl2 | 607 | 69292.6835 | -0.4289951 | 0 | 0.28336079 | 0.14991763 | 0.29159802 | Commercial - Sigma Aldrich Cat A2153-100G >96% | P02769 | 3V03 | |
| caspase1_zymogen | Caspase-1 | Zymogen form | Protease, inflammatory processes | 1 | 50 mM HEPES, pH 7.40, 250 mM NaCl | 404 | 45158.155 | -0.3334158 | 11 | 0.26980198 | 0.22524752 | 0.25247525 | Andy Ambrose, Arkin Lab, UCSF | P29466 | 1BMQ | |
| caspase2_active | Caspase-2 | Active form | Protease, apoptotic cascade | 1 | 50 mM HEPES, pH 7.40, 250 mM NaCl | 452 | 50684.7558 | -0.3011062 | 6 | 0.28318584 | 0.20132743 | 0.29867257 | Andy Ambrose, Arkin Lab, UCSF | P42575 | 1PYO | |
| caspase3_active | Caspase-3 | Active form | Protease, apoptotic cascade | 1 | 50 mM HEPES, pH 7.40, 250 mM NaCl | 277 | 31607.5495 | -0.4570397 | 12 | 0.28519856 | 0.23104693 | 0.22382671 | Andy Ambrose, Arkin Lab, UCSF | P42574 | 1CP3 | |
| caspase6_active | Caspase-6 | Active form | Protease, programmed cell death, development, innate immunity, inflammation | 1 | 50 mM HEPES, pH 7.40, 250 mM NaCl | 293 | 33309.597 | -0.3552901 | 6 | 0.29692833 | 0.19795222 | 0.24573379 | Andy Ambrose, Arkin Lab, UCSF | P55212 | 2WDP | |
| caspase6_zymogen | Caspase-6 | Zymogen form | Protease, programmed cell death, development, innate immunity, inflammation | 1 | 50 mM HEPES, pH 7.40, 250 mM NaCl | 293 | 33309.597 | -0.3552901 | 6 | 0.29692833 | 0.19795222 | 0.24573379 | Andy Ambrose, Arkin Lab, UCSF | P55212 | 2WDP | |
| caspase7_active | Caspase-7 | Active form | Protease, apoptotic cascade | 1 | 50 mM HEPES, pH 7.40, 250 mM NaCl | 303 | 34276.4229 | -0.5089109 | 9 | 0.27392739 | 0.21452145 | 0.21122112 | Andy Ambrose, Arkin Lab, UCSF | P55210 | 1F1J | |
| caspase9_active | Caspase-9 | Active form | Protease, apoptotic cascade | 1 | 50 mM HEPES, pH 7.40, 250 mM NaCl | 416 | 46280.3047 | -0.2370192 | 10 | 0.29807692 | 0.24519231 | 0.25721154 | Andy Ambrose, Arkin Lab, UCSF | P55212 | 1JXQ | |
| CBP_KIX | Histone lysine acetyltransferase CREBBP | | Histone acetylation, transcriptional activation | 10 | 10 mM HEPES, pH 7.20, 200 mM NaCl | 81 | 9534.7203 | -0.8037037 | 0 | 0.2962963 | 0.13580247 | 0.32098765 | Partch Lab, UCSC | P45481 | 1KDX_1 | Gustafson, Chelsea L., et al. 2017 |
| CHIP | E3 ubiquitin-protein ligase CHIP | | E3 ubiquitin ligase | 1 | 10 mM HEPES, pH 7.20, 200 mM NaCl, 1 mM TCEP | 303 | 34855.8215 | -0.8108911 | 20 | 0.24422442 | 0.18151815 | 0.31353135 | Matt Ravalin, Gestwicki Lab, UCSF | Q9UNE7 | 4KBQ | Ravalin M, et al. 2019 |
| CLOCK_PASB_W362A | Circadian locomoter output cycles protein kaput | point mutant reduces formation of soluble aggregates at room temp | Transcriptional activator, circadiam rhythm | 10 | 10 mM HEPES, pH 7.20, 200 mM NaCl | 361 | 42025.9864 | -0.5260388 | 8 | 0.31301939 | 0.19944598 | 0.23822715 | Partch Lab, UCSC | O08785 | | Michael, Alicia K., et al. 2017 |
| CoV2Orf9bMark2 | SARS-CoV2 complex Orf9b-Mark2 | screened as pre-assembled complex | SARS-CoV2 host-virus protein complex | 10 | 50 mM Tris HCL, pH 7.5, 150 mM NaCl, 0.1 mM TCEP | | | | | | | | Amy Diallo, Verba Lab, UCSF | P0DTD1 | | |
| CP_BMAL1_PASB_W427A | Aryl hydrocarbon receptor nuclear translocator-like protein 1 | point mutant reduces formation of soluble aggregates at room temp | Transcriptional activator, circadiam rhythm | 10 | 10 mM HEPES, pH 7.20, 200 mM NaCl | 387 | 44123.1476 | -0.4434109 | 7 | 0.28423773 | 0.20671835 | 0.23514212 | Partch Lab, UCSC | O00327 | 4F3I | Michael, Alicia K., et al. 2017 |
| CP581_CBPKIX | CREB-binding protein | | Histone acetylation, transcriptional activation | 10 | 10 mM HEPES, pH 7.20, 200 mM NaCl | 80 | 9405.6063 | -0.77 | 0 | 0.3 | 0.1375 | 0.3125 | Partch Lab, UCSC | Q92793 | 1SB0 | Gustafson, Chelsea L., et al. 2017 |
| diUb | K48 linked di-Ubiquitin | | Post-translational protein tag, protein regulation and degradation | 5 | 50 mM Tris HCl, 150 mM NaCl, 1% DMSO, 0.01% Triton X-100, 1 mM TCEP | 76 | 8548.6933 | -0.5605263 | 26 | 0.28947368 | 0.19736842 | 0.22368421 | Commercial - R&D Systems UCN-200 | P0CG48 | 1UBQ | |
| eIF2B_with_ISRIB_all | Eukaryotic translation initiation factor 2 subunit 2 | Screened as pre-assembed complex | Initition factor in eukaryotic protein synthesis | 0.25 | 50 mM TRIS, pH 7.4, 150 mM NaCl, 2 mM MgCl2, 1 mM DTT | 2369 | 262753.142 | -0.2064584 | | 0.30434783 | 0.22119038 | 0.28366399 | Michael Schoof, Walter Lab, UCSF | | 6CAJ | Tsai JC, et al. 2018 |
| eIF2B_with_ISRIB_alpha | Eukaryotic translation initiation factor 2 subunit 2, alpha subunit | Screened as pre-assembed complex | Initition factor in eukaryotic protein synthesis | 0.25 | 50 mM TRIS, pH 7.4, 150 mM NaCl, 2 mM MgCl2, 1 mM DTT | 305 | 33711.8019 | 0.07409836 | 0 | 0.3442623 | 0.17704918 | 0.29508197 | Michael Schoof, Walter Lab, UCSF | | 6CAJ_3 | Tsai JC, et al. 2018 |
| eIF2B_with_ISRIB_beta | Eukaryotic translation initiation factor 2 subunit 2, beta subunit | Screened as pre-assembed complex | Initition factor in eukaryotic protein synthesis | 0.25 | 50 mM TRIS, pH 7.4, 150 mM NaCl, 2 mM MgCl2, 1 mM DTT | 368 | 40952.1788 | -0.2054348 | 22 | 0.28804348 | 0.22282609 | 0.30706522 | Michael Schoof, Walter Lab, UCSF | | 6CAJ_4 | Tsai JC, et al. 2018 |
| eIF2B_with_ISRIB_delta | Eukaryotic translation initiation factor 2 subunit 2, deta subunit | Screened as pre-assembed complex | Initition factor in eukaryotic protein synthesis | 0.25 | 50 mM TRIS, pH 7.4, 150 mM NaCl, 2 mM MgCl2, 1 mM DTT | 523 | 57556.561 | -0.3544933 | 32 | 0.27724665 | 0.23709369 | 0.29063098 | Michael Schoof, Walter Lab, UCSF | | 6CAJ_5 | Tsai JC, et al. 2018 |
| eIF2B_with_ISRIB_epsilon | Eukaryotic translation initiation factor 2 subunit 2, epsilon subunit | Screened as pre-assembed complex | Initition factor in eukaryotic protein synthesis | 0.25 | 50 mM TRIS, pH 7.4, 150 mM NaCl, 2 mM MgCl2, 1 mM DTT | 721 | 80364.9077 | -0.2768377 | 17 | 0.30651872 | 0.23162275 | 0.27461859 | Michael Schoof, Walter Lab, UCSF | | 6CAJ_1 | Tsai JC, et al. 2018 |
| eIF2B_with_ISRIB_gamma | Eukaryotic translation initiation factor 2 subunit 2, gamma subunit | Screened as pre-assembed complex | Initition factor in eukaryotic protein synthesis | 0.25 | 50 mM TRIS, pH 7.4, 150 mM NaCl, 2 mM MgCl2, 1 mM DTT | 452 | 50239.7541 | -0.1130531 | 4 | 0.31858407 | 0.21460177 | 0.26327434 | Michael Schoof, Walter Lab, UCSF | | 6CAJ_2 | Tsai JC, et al. 2018 |
| FKBP12 | Peptidyl-prolyl cis-trans isomerase FKBP1A | | Kinase receptor regulation | 10 | 20 mM Tris, pH 8.0 | 108 | 11950.569 | -0.4111111 | 15 | 0.27777778 | 0.24074074 | 0.21296296 | Douglas Wassarman, Shokat Lab, UCSF | P62942 | | Wassarman, Douglas R., et al. 2022 |
| FL_B5_no_inc_protein_1 | Alpha-crystallin B chain | Phosphomimetic mutant, no pre-incubation | Oligomeric/conformational modulation of HSPB5 | | | 175 | 20158.6422 | -0.5314286 | 21 | 0.30285714 | 0.25142857 | 0.21142857 | Maria Janowska, Klevit Lab, UW | P02511 | | Rajagopal, Ponni, et al. 2015 |
| FL_B5_pre_inc_protein_2 | Alpha-crystallin B chain | pre-incubated at 37C to activate | Oligomeric/conformational modulation of HSPB5 | 10 | 25 mM MOPS, 150 mM NaCl, pH 7.50 | 175 | 20158.6422 | -0.5314286 | 21 | 0.30285714 | 0.25142857 | 0.21142857 | Maria Janowska, Klevit Lab, UW | P02511 | | Rajagopal, Ponni, et al. 2015 |
| GB1 | Immunoglobulin G-binding protein G | | Binds to immunoglobulin Fc region | 10 | 10 mM HEPES, pH 7.20, 200 mM NaCl | 56 | 6151.7266 | -0.5392857 | 4 | 0.25 | 0.125 | 0.28571429 | Partch Lab, UCSC | P06654 | 2J52 | |
| GS_R324C | Glutamine synthetase | Pathogenic mutant of glutamine synthetase | Implicated in Glutamine Deficiency, a rare genetic disease | 10 | 20 mM HEPES, pH 7.5, 150 mM KCl, 20 mM NaCl, 5 mM MgCl2 | 384 | 43252.2819 | -0.5648437 | 5 | 0.25520833 | 0.26302083 | 0.22916667 | Erin Thompson, Fraser Lab, UCSF | P15104 | | Thompson, Erin et al. 2020 |
| GS_R324S | Glutamine synthetase | Pathogenic mutant of glutamine synthetase | Implicated in Glutamine Deficiency, a rare genetic disease | 10 | 20 mM HEPES, pH 7.5, 150 mM KCl, 20 mM NaCl, 5 mM MgCl2 | 384 | 43236.2163 | -0.5734375 | 7 | 0.25520833 | 0.265625 | 0.22916667 | Erin Thompson, Fraser Lab, UCSF | P15104 | | Thompson, Erin et al. 2020 |
| GS_R341C | Glutamine synthetase | Pathogenic mutant of glutamine synthetase | Implicated in Glutamine Deficiency, a rare genetic disease | 10 | 20 mM HEPES, pH 7.5, 150 mM KCl, 20 mM NaCl, 5 mM MgCl2 | 384 | 43252.2819 | -0.5648437 | 4 | 0.25520833 | 0.26302083 | 0.22916667 | Erin Thompson, Fraser Lab, UCSF | P15104 | | Thompson, Erin et al. 2020 |
| GS_WT | Glutamine synthetase | Glutamine synthetase | Metabolic enzyme, ATP-dependent conversion of glutamate to glutamine | 10 | 20 mM HEPES, pH 7.5, 150 mM KCl, 20 mM NaCl, 5 mM MgCl2 | 384 | 43305.3247 | -0.5830729 | 7 | 0.25520833 | 0.26302083 | 0.22916667 | Erin Thompson, Fraser Lab, UCSF | P15104 | 2OJW | Thompson, Erin et al. 2020 |
| GST | Glutathione S-transferase A3 | | Enzymatic conjugation of reduced glutatione to various electrophiles | 1 | 10 mM HEPES, pH 7.20, 200 mM NaCl | 222 | 25740.7871 | -0.3189189 | 3 | 0.32882883 | 0.16216216 | 0.32882883 | Partch Lab, UCSC | Q16772 | 5LCZ | |
| H104_FL_B5_pre_inc_protein_5 | Alpha-crystallin B chain | Phosphomimetic mutant of HSPB5, incubated at 37C to activate | Oligomeric/conformational modulation of HSPB5 | 10 | 25 mM MOPS, 150 mM NaCl, pH 7.50 | 175 | 20149.6752 | -0.5354286 | 20 | 0.30285714 | 0.25142857 | 0.21142857 | Maria Janowska, Klevit Lab, UW | P02511 | | Rajagopal, Ponni, et al. 2020 |

| | | | | | | | | | | | | | | | | |
|---|---|---|---|---|---|---|---|---|---|---|---|---|---|---|---|---|
| hACE2 | Angiotensin-converting enzyme 2 | Hormone regulation in cardiovascular homeostasis | Carboxypeptidase essential for cardiovasular homeostasis | 0.5 | 50 mM Tris HCl, 150 mM NaCl, 1% DMSO, 0.01% Triton X-100, 1 mM TCEP | 723 | 83594.7348 | -0.4614108 | 2 | 0.31673582 | 0.22821577 | 0.26970954 | Commercial - R&D Systems R&D 933-ZN-101 | Q9BYF1 | 1R42 | |
| HIP | Hsc70-interacting protein | | State-dependent protein-interacting partner for nucleotide domain of Hsp70 | 1 | 10 mM HEPES, pH 7.20, 200 mM NaCl | 369 | 41331.2623 | -0.8574526 | 69 | 0.19783198 | 0.21680217 | 0.34146341 | Rebecca Freilich, Gestwicki Lab, UCSF | P50502 | | |
| His_NusA | Transcription termination/antitermination protein NusA | | rRNA transciption and and antitermination complex component | 10 | 10 mM HEPES, pH 7.20, 200 mM NaCl | 329 | 36252.7006 | -0.2179331 | 2 | 0.2887538 | 0.16717325 | 0.32826748 | Partch Lab, UCSC | P0AFF6 | 5LM9 | |
| HisPER2_AB_I324N | Period circadian protein homolog 2 | 324N "Edo" mutant in interdomain linker | Transcriptional repressor complex component in circadium rhythm | 10 | 10 mM HEPES, pH 7.20, 200 mM NaCl | 304 | 34638.3344 | -0.2555921 | 9 | 0.31907895 | 0.23355263 | 0.22697368 | Partch Lab, UCSC | O54943 | 4F3L | Militi, Stefania, et al. 2016 |
| HisSUMO | Small ubiquitin-related modifier 4 | | Post-translational protein tag | 1 | 10 mM HEPES, pH 7.20, 200 mM NaCl | 80 | 9237.4484 | -0.945 | 30 | 0.2375 | 0.2 | 0.25 | Partch Lab, UCSC | Q6EEV6 | 2ASQ | |
| Hsc70 | Heat shock cognate 71 kDa protein | | Molecular chaperone | 10 | 10 mM HEPES, pH 7.20, 200 mM NaCl | 646 | 70897.2343 | -0.4560372 | 20 | 0.27089783 | 0.22755418 | 0.24148607 | Rebecca Freilich, Gestwicki Lab, UCSF | P11142 | 3AGY | |
| Hsp10 | 10 kDa heat shock protein, mitochondrial | | Mitochondrial molecular chaperonin | 10 | 10 mM HEPES, pH 7.20, 200 mM NaCl | 80 | 8343.6766 | -0.02 | 4 | 0.3 | 0.2625 | 0.25 | Dr. Hao Shao, Gestwicki Lab, UCSF | P61604 | 4JP1 | |
| Hsp60 | 60 kDa heat shock protein, mitochondrial | | Mitochondrial molecular chaperonin | 10 | DPBS | 558 | 59777.8702 | -0.168638 | 10 | 0.28853047 | 0.20430108 | 0.27598566 | Dr. Hao Shao, Gestwicki Lab, UCSF | P10809 | 4JP1 | |
| Hsp72_SBD | 72kDa Heat shock protein, substrate binding domain | Substrate binding domain only | Molecular chaperone | 10 | 25 mM Tween, 100 mM NaCl, 1mM TCEP, 1 mM EDTA | 235 | 26241.3454 | -0.5902128 | 23 | 0.26382979 | 0.20425532 | 0.28510638 | Rebecca Freilich, Gestwicki Lab, UCSF | P54652 | | |
| ISG15 | Ubiquitin-like protein ISGylation 15 | | Innate immune signalling | 5 | 50 mM Tris HCl, 150 mM NaCl, 1% DMSO, 0.01% Triton X-100, 1 mM TCEP | 165 | 17887.294 | -0.1387879 | 5 | 0.31515152 | 0.27272727 | 0.26666667 | Commercial - R&D Systems UL-601 | P05161 | 1Z2M | |
| KaiC_AE | Circadian clock protein kinase KaiC | Phosphomimetic of the S431/pT432 day-time state of KaiC | Cyanobacterial circadium rhythm oscillator, protein-protein interaction hub | 10 | 25 mM Tris, pH 7.40, 130 mM NaCl, 1 mM MgCl2, 1 mM TCEP, 1 mM ATP | 519 | 58014.2318 | -0.2231214 | 6 | 0.30828516 | 0.22928709 | 0.24662813 | Jeffery Swan, Partch Lab, UCSC | Q79V60 | 2GBL | |
| KaiC_EA | Circadian clock protein kinase KaiC | Phosphomimetic of the pS431/T432 night-time state of KaiC | Cyanobacterial circadium rhythm oscillator, protein-protein interaction hub | 10 | 25 mM Tris, pH 7.40, 130 mM NaCl, 1 mM MgCl2, 1 mM TCEP, 1 mM ATP | 519 | 58014.2318 | -0.2231214 | 6 | 0.30828516 | 0.22928709 | 0.24662813 | Jeffery Swan, Partch Lab, UCSC | Q79V60 | 2GBL | Tseng, Roger, et al. 2017 |
| kIvL_light_chain | Kappa-I variable domain of immunoglobulin light chain | | Aggregates in genetic disease Light Chain Amyloidosis | 17.5 | 50 mM TRIS, pH 7.4, 150 mM NaCl, 2 mM MgCl2 | 108 | 11932.0587 | -0.5222222 | 5 | 0.27777778 | 0.2962963 | 0.15740741 | Marina Ramirz-Alvarado | | | |
| lysozyme | Hen egg white lysozyme | | Egg white innate immunity , model folding protein | 5 | 10 mM HEPES, pH 7.20, 200 mM NaCl | 129 | 14313.004 | -0.472093 | 0 | 0.24806202 | 0.29457364 | 0.18604651 | Commercial - Sigma Adrich 10837059001 | P00698 | 1DPX | |
| mortalin | Stress-70 protein, mitochondrial "Mortalin" | | Mitochondrial chaperone | 1 | 20 mM HEPES ,pH 7.50, 100 mM NaCl | 679 | 73679.6436 | -0.4002946 | 24 | 0.25184094 | 0.21502209 | 0.27982327 | Dr. Hao Shao, Gestwicki Lab, UCSF | P38646 | 3N8E | |
| MPro | SARS-CoV2 3C-like proteinase "Main protease" | | Viral protease | 1 | 50 mM Tris HCl, 150 mM NaCl, 1% DMSO, 0.01% Triton X-100, 1 mM TCEP | 306 | 33796.295 | -0.0189542 | 0 | 0.32026144 | 0.24836601 | 0.2124183 | Commercial - R&D Systems E-720-050 | P0DTD1 | 6Z2E | |
| nhr23_LBD | Nuclear hormonr receptor 23 | ligand biding domain alone | Hormone receptor involved in skin regeneration and molting of C. elegans | 1 | 10 mM HEPES, pH 7.20, 200 mM NaCl, 1 mM TCEP | 242 | 26998.3636 | -0.0681818 | 2 | 0.30991736 | 0.18181818 | 0.31404959 | Partch Lab, UCSC | P41828 | | |
| NPAS2_PAS_A | Neuronal PAS domain-containing protein 2 | Pas A domain | | 10 | 10 mM HEPES, pH 7.20, 200 mM NaCl | 163 | 18383.6534 | -0.0650307 | 0 | 0.35582822 | 0.27607362 | 0.2392638 | Partch Lab, UCSC | P97460 | | |
| nsp10 | SARS-CoV2 nonstructural protein 10 | screened in isolation (not in presence of nsp16 or 14) | Viral immune evasion complex component | 1 | 50 mM HEPES, pH 8, 500 mM NaCl, 2 mM MgCl2, 0.5 mM TCEP | 142 | 15062.0745 | -0.0838028 | 4 | 0.22535211 | 0.25352113 | 0.21126761 | Victor Long, Gross Lab, UCSF | P0DTD1 | 6W4H | |
| nsp14 | SARS-CoV2 nonstructural protein 14 | screened in isolation (not in presence of nsp10) | Viral methyltransferase, immune evasion | 1 | 50 mM HEPES, pH 8, 500 mM NaCl, 2 mM MgCl2, 0.5 mM TCEP | 527 | 59815.0269 | -0.1335863 | 1 | 0.33017078 | 0.21252372 | 0.19734345 | Victor Long, Gross Lab, UCSF | P0DTD1 | 7N0B_2 | |
| nsp14_nsp10 | Heterodimeric complex of SARS-CoV2 nonstructural protein 14 and 10 | screened as a pre-assembled complex | Viral methyltransferase, immune evasion | 1 | 50 mM HEPES, pH 8, 500 mM NaCl, 2 mM MgCl2, 0.5 mM TCEP | 669 | 74859.0861 | -0.1230194 | | 0.30792227 | 0.22122571 | 0.20029895 | Victor Long, Gross Lab, UCSF | P0DTD1 | 7N0B | |
| nsp16 | SARS-CoV2 nonstructural protein 16 | screened in isolation (not in presence of nsp10) | Viral RNA capping, immune evasion | 1 | 50 mM HEPES, pH 8, 500 mM NaCl, 2 mM MgCl2, 0.5 mM TCEP | 301 | 33595.2116 | -0.0933555 | 1 | 0.32890365 | 0.25249169 | 0.24252492 | Victor Long, Gross Lab, UCSF | P0DTD1 | 6W4H | |
| nsp16_nsp10 | Heterodimeric complex of SARS-CoV2 nonstructural protein 16 and 10 | screend as a pre-assembled complex | Viral RNA capping, immune evasion | 1 | 50 mM HEPES, pH 8, 500 mM NaCl, 2 mM MgCl2, 0.5 mM TCEP | 443 | 48639.2708 | -0.0902935 | | 0.29571106 | 0.25282167 | 0.23250564 | Victor Long, Gross Lab, UCSF | P0DTD1 | 6W4H | |
| nsp2 | SARS-CoV2 nonstructural protein 2 | | Viral protein with putative zinc-finger | 1 | 30 mM HEPES, pH 7.50, 60 mM NaCl, 1 mM TCEP, 10 uM ZnCl2 | 638 | 70510.6111 | -0.062069 | 0 | 0.31661442 | 0.21630094 | 0.26332288 | Meghna Gupta and Klim Verba, Stroud and Verba Lab, UCSF | P0DTD1 | 7MSX_1 | |
| nsp3_mac1 | SARS-CoV2 nonstructural protein 3, macrodomain 1 | macrodomain 1 of SARS-CoV2 non-structural protein 3 | Viral modulation/evasion of host innate immunity | 2 | 50 mM Tris HCl, 150 mM NaCl, 1% DMSO, 0.01% Triton X-100, 1 mM TCEP | 176 | 18971.4669 | -0.0198864 | 0 | 0.34090909 | 0.26136364 | 0.25568182 | Galen Correy, Fraser Lab, UCSF | P0DTD1 | 6WOJ | |
| nucleocapsid | SARS-CoV2 Nucleocapsid protein | | Viral genome packing | 1 | 50 mM Tris HCl, 150 mM NaCl, 1% DMSO, 0.01% Triton X-100, 1 mM TCEP | 419 | 45625.1383 | -0.9713604 | 68 | 0.18615752 | 0.31026253 | 0.19809069 | Commercial - R&D Systems 10474-CV-050 | P0DTD1 | | |
| Nucleosome_all | Nucleosome | Screened as a fully assembled nucleosome with Widom 601 DNA | Key component of chromatin organization and regulation | 1 | TCS | 495 | 54742.3457 | -0.5757576 | | 0.24848485 | 0.2020202 | 0.26464646 | Hayden Saunders, Narlikar Lab, UCSF | | 2CV5_2 | Gamarra, Nathan et al. 2021 |
| Nucleosome_H2A | Histone protein 2A | Screen as a pre-assembled histone mixture | Key component of chromatin organization and regulation | 1 | TCS | 130 | 14135.3121 | -0.4776923 | 28 | 0.26153846 | 0.23076923 | 0.30769231 | Hayden Saunders, Narlikar Lab, UCSF | Q6FI13 | 2CV5_4 | Gamarra, Nathan et al. 2021 |
| Nucleosome_H2B | Histone protein 2B | Screen as a pre-assembled histone mixture | Key component of chromatin organization and regulation | 1 | TCS | 126 | 13889.9692 | -0.7174603 | 33 | 0.22222222 | 0.23015873 | 0.23809524 | Hayden Saunders, Narlikar Lab, UCSF | Q93079 | 2CV5_5 | Gamarra, Nathan et al. 2021 |
| Nucleosome_H3 | Histone protein 3 | Screen as a pre-assembled histone mixture | Key component of chromatin organization and regulation | 1 | TCS | 136 | 15403.9072 | -0.5794118 | 34 | 0.23529412 | 0.13970588 | 0.29411765 | Hayden Saunders, Narlikar Lab, UCSF | P68431 | 2CV5_2 | Gamarra, Nathan et al. 2021 |
| Nucleosome_H4 | Histone protein 4 | Screen as a pre-assembled histone mixture | Key component of chromatin organization and regulation | 1 | TCS | 103 | 11367.2031 | -0.5213592 | 20 | 0.2815534 | 0.21359223 | 0.2038835 | Hayden Saunders, Narlikar Lab, UCSF | P62805 | 2CV5_3 | Gamarra, Nathan et al. 2021 |
| Nurr1_LBD | Nuclear receptor subfamily 4 group A member 2 | Ligand binding domain only | Transcriptional regulator, dopaminergic neuron development | 1 | | 272 | 30627.0299 | -0.1808824 | 12 | 0.33088235 | 0.25367647 | 0.26102941 | Ward and Partch Lab, UCSC | P43354 | | |
| OGTase | UDP-N-acetylglucosamine-peptide N-acetylglucosaminyltransferase | | O-linked GlcNAcylation enzyme, cellular process regulation | 1 | 50 mM HEPES, 300 mM NaCl, pH 7.40, 1 mM DTT | 1046 | 116923.285 | -0.2424474 | 2 | 0.29923518 | 0.21797323 | 0.29445507 | Matt Alteen, Volcadlo Lab, U Alberta | O15294 | | Alteen, Matthew G., et al. 2020 |

| | | | | | | | | | | | | | | | | |
|---|---|---|---|---|---|---|---|---|---|---|---|---|---|---|---|---|
| p23 | Prostaglandin E synthase 3 | | Hormone-dependent transcriptional activation | 5 | 50 mM Tris, pH 9.0, 150 mM NaCl | 160 | 18697.1939 | -1.049375 | 26 | 0.2375 | 0.2375 | 0.2125 | Andrew Ambrose, Arkin Lab, UCSF | Q15185 | 1EJF | |
| p300_KIX | Histone acetyltransferase p300 | KIX domain | Histone acetylation, chromatin remodeling | 1 | 10 mM HEPES, pH 7.20, 200 mM NaCl | 80 | 9405.6063 | -0.77 | 0 | 0.3 | 0.1375 | 0.3125 | Partch Lab, UCSC | Q09472 | 2KWF | |
| PER2_PAS_ASB_wt | Period circadian protein homolog 2 | wild-type tandem PAS A and B domains | Transcriptional repressor complex component in circadium rhythm | 10 | 10 mM HEPES, pH 7.20, 200 mM NaCl | 304 | 34637.3894 | -0.2292763 | 7 | 0.32236842 | 0.23026316 | 0.22697368 | Partch Lab, UCSC | O54943 | | Militi, Stefania, et al. 2016 |
| PLPro | SARS-CoV2 Papain-like protease | | Viral polyprotein cleavage, host immune evagion | 1 | 50 mM Tris HCl, 150 mM NaCl, 1% DMSO, 0.01% Triton X-100, 1 mM TCEP | 315 | 35631.1902 | -0.3749206 | 0 | 0.2984127 | 0.19365079 | 0.22539683 | Commercial - R&D Systems E-611 | P0DTD1 | 7CMD | |
| pp5 | Serine/threonine-protein phosphatase 5 | | Modulation of diverse signalling pathways via dephosphorylation | 1 | 20 mM HEPES, pH 7.50, 50 mM KCl, 1 mM TCEP | 556 | 61859.1062 | -0.1893885 | 5 | 0.30935252 | 0.22302158 | 0.28057554 | Aye Thwin, Gestwicki and Southworth Lab, UCSF | P53041 | 1WAO | |
| PPIE | Peptidyl-prolyl cis-trans isomerase E | | pre-mRNA splicing, spliceosome component | 10 | 10 mM HEPES, pH 7.20, 200 mM NaCl | 301 | 33430.4114 | -0.5212625 | 14 | 0.26245847 | 0.23920266 | 0.23920266 | Partch Lab, UCSC | Q9UNP9 | 3UCH | |
| PPIEdeltaN | Peptidyl-prolyl cis-trans isomerase E | Isomerase domain only | pre-mRNA splicing, spliceosome component | 10 | 10 mM HEPES, pH 7.20, 200 mM NaCl | 170 | 18717.1834 | -0.4417647 | 5 | 0.26470588 | 0.27058824 | 0.17058824 | Partch Lab, UCSC | Q9UNP9 | 2R99 | |
| PPIG | Peptidyl-prolyl cis-trans isomerase G | | Proline isomerization, protein folding | 10 | 10 mM HEPES, pH 7.20, 200 mM NaCl | 754 | 88616.1421 | -1.8831565 | 81 | 0.12732095 | 0.28647215 | 0.16180371 | Partch Lab, UCSC | Q13427 | 2GW2 | |
| PPIL1 | Peptidyl-prolyl cis-trans isomerase-like 1 | | pre-mRNA splicing, spliceosome component | 10 | 10 mM HEPES, pH 7.20, 200 mM NaCl | 166 | 18236.6204 | -0.3042169 | 5 | 0.28313253 | 0.26506024 | 0.19879518 | Partch Lab, UCSC | Q9Y3C6 | 2X7K | |
| PPWD1 | Peptidylprolyl isomerase domain and WD repeat-containing protein 1 | | Proline isomerization, protein folding | 10 | 10 mM HEPES, pH 7.20, 200 mM NaCl | 646 | 73574.0335 | -0.4479876 | 16 | 0.28947368 | 0.21362229 | 0.21671827 | Partch Lab, UCSC | Q96BP3 | 2A2N | |
| proteasome_alpha | Archael proteasome | screened as a single, pre-assembled complex with beta subunit | Non-specific protein degradation | 1 | 50 mM Tris pH 7.5,10 mM MgCl2, 1 mM DTT | 233 | 25799.1671 | -0.211588 | 0 | 0.32618026 | 0.19313305 | 0.27467811 | Sarah Williams, Gestwicki Lab, UCSF | P25156 | 3IPM_1 | Opoku-Nsiah, Kwadwo A., et al. 2021 |
| proteasome_beta | Archael proteasome | screened as a single, pre-assembled complex with alpha subunit | Non-specific protein degradation | 1 | 50 mM Tris pH 7.5,10 mM MgCl2, 1 mM DTT | 217 | 23969.4185 | -0.0175115 | 1 | 0.31797235 | 0.18894009 | 0.26728111 | Sarah Williams, Gestwicki Lab, UCSF | P28061 | 3IPM_2 | Opoku-Nsiah, Kwadwo A., et al. 2021 |
| RBD | SARS-CoV2 spike protein receptor-binidng somain | | Facilitates viral entry into host cells | 1 | 50 mM Tris HCl, 150 mM NaCl, 1% DMSO, 0.01% Triton X-100, 1 mM TCEP | 223 | 25098.1047 | -0.2587444 | 0 | 0.34080717 | 0.29596413 | 0.14798206 | Commercial - R&D Systems R&D 10500-CV-100 | P0DTD1 | 7CH5 | |
| Retinoic_acid_receptor_alpha | Retinoic acid receptor RXR-alpha | | Transcription factor | 10 | | 210 | 23633.3325 | 0.06238095 | 0 | 0.34285714 | 0.17619048 | 0.35238095 | Mericka McCabe, Cuervo Lab, Albert Einstein College of Medicine | P19793 | 5EC9 | Anguiano, Jaime, et al. 2013 |
| snf2h | SWI/SNF-related matrix-associated actin-dependent regulator of chromatin subfamily A member 5 | | Helicase, chromatin remodeling | 1 | 15 mM HEPES, pH 7.5, 140 mM KCl, 1 mM Mg-ADP-BeFx | 1052 | 121903.937 | -0.7191065 | 25 | 0.2756654 | 0.20627376 | 0.27756654 | Hayden Saunders, Narlikar Lab, UCSF | O60264 | 6NE3 | |
| Widom_601_DNA | Widom 601 sequence, nucleosomal DNA | Screened as DNA alone | Classical DNA sequence for in vitro nucleosome assembly | 1 | TAE | | | | | | | | Hayden Saunders, Narlikar Lab, UCSF | | | Gamarra, Nathan et al. 2021 |


Alteen, Matthew G., et al. "A direct fluorescent activity assay for glycosyltransferases enables convenient high-throughput screening: Application to O-GlcNAc Transferase." Angewandte Chemie International Edition 59.24 (2020): 9601-9609.

Anguiano, Jaime, et al. "Chemical modulation of chaperone-mediated autophagy by retinoic acid derivatives." Nature chemical biology 9.6 (2013): 374-382.

Gamarra, Nathan, and Geeta J. Narlikar. "Histone dynamics play a critical role in SNF2h-mediated nucleosome sliding." Nature structural & molecular biology 28.7 (2021): 548-551.

Gustafson, Chelsea L., et al. "A slow conformational switch in the BMAL1 transactivation domain modulates circadian rhythms." Molecular cell 66.4 (2017): 447-457.

Michael, Alicia K., et al. "Formation of a repressive complex in the mammalian circadian clock is mediated by the secondary pocket of CRY1." Proceedings of the National Academy of Sciences 114.7 (2017): 1560-1565.

Militi, Stefania, et al. "Early doors (Edo) mutant mouse reveals the importance of period 2 (PER2) PAS domain structure for circadian pacemaking." Proceedings of the National Academy of Sciences 113.10 (2016): 2756-2761.

Opoku-Nsiah, Kwadwo A., et al. "The YΦ motif defines the structure-activity relationships of human 20S proteasome activators." Nature communications 13.1 (2022): 1-12.

Rajagopal, Ponni, et al. "A conserved histidine modulates HSPB5 structure to trigger chaperone activity in response to stress-related acidosis." Elife 4 (2015): e07304.

Ravalin M, Theofilas P, Basu K, Opoku-Nsiah KA, Assimon VA, Medina-Cleghorn D, Chen YF, Bohn MF, Arkin M, Grinberg LT, Craik CS, Gestwicki JE. Specificity for latent C termini links the E3 ubiquitin ligase CHIP to caspases. Nat Chem Biol. 2019 Aug;15(8):786-794. doi: 10.1038/s41589-019-0322-6. Epub 2019 Jul 18. PMID: 31320752; PMCID: PMC6693490.

Thompson, Erin. Mechanism of compensation for disease variants in human glutamine synthetase. University of California, San Francisco, 2020.

Tsai JC, Miller-Vedam LE, Anand AA, Jaishankar P, Nguyen HC, Renslo AR, Frost A, Walter P. Structure of the nucleotide exchange factor eIF2B reveals mechanism of memory-enhancing molecule. Science. 2018 Mar 30;359(6383):eaaq0939. doi: 10.1126

Tseng, Roger, et al. "Structural basis of the day-night transition in a bacterial circadian clock." Science 355.6330 (2017): 1174-1180.

Wassarman, Douglas R., et al. "Tissue-Restricted Inhibition of mTOR Using Chemical Genetics." bioRxiv (2022).


**Table S2. Protein panel and biochemical properties.** A complete list of all proteins which underwent dye screening for this manuscript. Dye screening conditions and calculated biochemical parameters are included, as well as construct specifications, PDB codes, protein donors and construct-specific citations where applicable. Further information, including primary sequence and amino acid composition, are available in Data S3.

### paDSF dye screening procedure

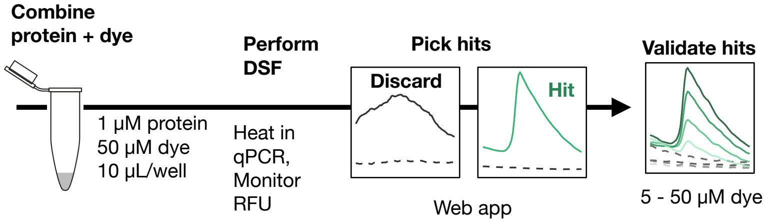


**Total protein:**
~100 μL 5 μM
(2 μL/dye)

**Total time:**
~2 hrs active
(6 hrs total)

Conducting a paDSF dye screen is nearly identical to a typical DSF experiment, with two modifications: (1) a paDSF dye is used in place of the commercial dye (e.g. SYPRO Orange), and (2) during heating, fluorescence readings are taken in multiple wavelengths. Analyzing data from a paDSF dye screen is different from analyzing a typical DSF experiment, because the goal is to identify which dyes produce a clear, reliable melting curve for the protein.

**A general protocol for a paDSF screen testing 96 dyes against a single protein is provided below**. Specific guidance for adapting this protocol, such as altering the protein concentration and number of dyes tested, is provided below.

Video protocols of experimental set up are provided in Movie S1.

**Starting materials** (prepared before dye screen):
- 50 μM purified protein stock
- 50 mL buffer (e.g. 10 mM HEPES pH 7.20, 200 mM NaCl, 1 mM TCEP)
- A 384-well plate, containing 1 μL of 5 mM DMSO stock of a different dye individual wells
- A white, low-volume 384-well qPCR plate
- An optically-clear plate seal

**Experimental set-up**

1. Dilute protein stock to 5X final screening concentration by adding 21 μL of 50 μM protein stock to 189 μL buffer and mixing thoroughly, yielding 210 μL of 5 μM protein***.
   *Calculation: 2 μL of 5X protein is required for each dye tested. 2 μL per dye * 96 dyes = 192 μL protein solution needed total. Make 210 μL of protein solution, to accommodate for sample loss during pipetting. Standard final protein concentration in paDSF dye screens is 1 μM, so make 210 μL of 5 μM protein stock.*

2. Dilute dye stocks to 1.25X final screening concentration by adding 79 μL of buffer to each well of the pre-prepared dye-containing 384-well plate. Note: only 8μL of 1.25X dye solution is required, so if nanoliter-volume liquid handling is possible, this volume can be reduced, e.g., by adding 20 μL of buffer to 250 nL of 5 mM dye.
   *Calculation: 8 μL of 1.25X dye is required for each protein tested. 1.25X * 50 μM final concentration = 62.5 μM working dye concentration. 1 μL is the smallest volume that can be accurately pipetted, so 1 μL starting volume * (5 mM stock concentration / 62.5 μM desired concentration) = 80 μL resuspended volume per dye.*

3. Add 2 μL of 5X protein to 96 wells of the qPCR plate***, For the no-protein control, add 2 μL of buffer to the following 96 wells. Spin the plate down to settle droplets to the bottom of the well.

4. Transfer 8 μL of each 1.25X dye solution from the resuspended dye plate to the protein- and buffer-containing qPCR plate. Final well volumes are 10 μL, and each dye is present in two wells of the plate: one well containing 50 μM dye and 1 μM protein, and the other well containing 50 μM dye and buffer. Do not skip the no-protein control; it is crucial for accurate hit calling.

5. Seal the qPCR plate*** in optically-clear sealing film***, centrifuge briefly (e.g. 30 seconds at 1000 rcf) to remove bubbles.

6. Place the sealed qPCR plate in the qPCR instrument ***.

7. In the qPCR instrument, heat the plate from 25 ºC to 95 ºC at a rate of 1 ºC per minute***. At every degree, measure the fluorescence in FAM, JOE, TAMRA, ROX, Cy5, and Cy5.5 channels***.

*** See the following page for specific recommendations on adapting and modifying these conditions.

### paDSF dye screening procedure continued: hit validation

**A general protocol for the validation of 10 paDSF hits against a single protein is provided below**.
**Starting materials** (prepared before dye screen):

- 50 µM purified protein stock
- 50 mL buffer (e.g. 10 mM HEPES pH 7.20, 200 mM NaCl, 1 mM TCEP)
- A 384-well plate, containing 1 µL of 5, 2.5, 1.25, and 0.63 mM DMSO stock of each hit dye in individual wells
- A white, low-volume 384-well qPCR plate
- An optically-clear plate seal

**Experimental set-up**

1. Dilute protein stock to 5X final screening concentration by adding 10 µL of 50 µM protein stock to 90 µL buffer and mixing thoroughly, yielding 100 µL of 5 µM protein***.
   *Calculation: 2 µL of 5X protein is required for each concentration of each hit dye tested. 2 µL per dye * 10 dyes * 4 concentrations per dye = 80 µL protein solution needed total. Make 100 µL of protein solution, to accommodate for sample loss during pipetting. The same protein concentrations and volumes should be used for screening and validation.*

2. Dilute dye stocks to 1.25X final screening concentration by adding 79 µL of buffer to each well of the pre-prepared dye-containing 384-well plate. For validation, this plate should contain four concentrations of each hit dye, at decreasing two-fold serial dilutions in DMSO.
   *Calculation: if a dye was screened at 50 µM, validation should test 50, 25, 12.5, and 6.3 µM dye. As before, 8 µL of 1.25X dye is required for each protein tested.*

3. Proceed with steps 3 - 7 of the dye screening protocol, unmodified.

*** See the following page for specific recommendations on adapting and modifying these conditions.

### paDSF dye screening procedure continued: adapting the general procedure

#### *Adapting protein, buffer, and dyes*

1 of 3 | **Protein** Purpose: combines with paDSF dyes to find matches

**Protein requirements for paDSF dye screens**
In the standard procedure, each dye tested requires 10 uL of 1 uM protein (prepared first as a 5 μM 5X solution). In general, however much protein you would like to use to create a paDSF assay, we recommend using about half of that for the screen. The other half of the protein sample can then be used for hit validation and replicates of the final chosen conditions.

Recommended protein concentration
- **Standard: 1 μM**
- Lowest: 250 nM
- Highest: 20 μM

*Comment: like in standard DSF, protein concentrations in paDSF are flexible. Protein concentrations below 1 μM can be used for large proteins or complexes (>80 kDa or so), or to conserve precious samples. Protein concentrations above 10 μM may increase low hit rates for small proteins (<12 kDa or so), or produce marginally smoother data if protein is abundant. If the protein concentration is below 2 μM and screening data have only noisy or low-signal hits, increasing protein concentration is often the easiest fix. Increasing well volumes is less effective in our hands, and screening dyes at > 50 μM may introduce artifacts.*

Recommended well volume
- **Standard: 10 μL** in low-volume 384-well plates
- Lowest: plate manufacturers minimum well volume
- Highest: plate manufacturers maximum well volume

*Comment: like in standard DSF, well volumes in paDSF are flexible. Well volumes have rarely, if ever, majorly influenced results in our hands.*

Recommended screen size
- **Standard: 96 dyes (1000 μL of 1 μM protein)**
- Lowest: ~10 dyes
- Full library: 3 mL of 1 μM protein

*Comment: If protein is precious, fewer dyes can be tested. Most proteins get at least a couple of matches in the highest-matching 10 - 25 dyes. The 96 dyes tested in the concise sub-library have the highest protein match-rates, and most proteins get multiple matches with this library. Testing the full library (~300 dyes) will maximize the number of matches. If you want counter-selectivity against another protein (e.g. detection of one protein but not another), screening the full library can help, because the more protein-selective dyes are not included in the standard, highest-matching (and therefore, most protein-general) 96 dyes*.

2 of 3 | **50 mL buffer** Purpose: resuspends protein and dyes to appropriate concentrations. Also used in place of protein for protein-free negative controls.

Dye screens are broadly buffer compatible. We recommend screening in whatever buffer you like to use for your system. Standard additives such as DMSO, Triton, EDTA, and reducing agents are fine. High concentrations of glycerol (>5%) are tolerated, but not preferred, as viscous solutions can increase background fluorescence of many dyes.

3 of 3 | **Dyes** Purpose: combines with protein to find paDSF matches

To ease assembly, the dye library was designed to be largely commercially available. See Supporting Data Table 1 for detailed information on dye sources, how to assemble the dye library, and recommended subsets of the library to assemble. See Supporting Methods 1 for detailed information on dye handling and storage.
- **Standard: 1 μL of 5 mM dye in DMSO**, pre-dispensed into a well of a 384-well plate.
- If nL liquid handling possible: 250 nL of 5 mM dye in DMSO, pre-dispensed into a well of a 384-well plate.

### paDSF dye screening procedure continued: adapting the general procedure

#### *Adapting consumables, instruments, measurements*

| **White 384-well qPCR plate** | Contains the final DSF experiment, run in the qTower |
|---|---|

DSF can be plate-sensitive, so substitutions for qPCR plates are discouraged. We use Axygen PCR-384-LC480-W-NF, and have very few compatibility issues with this product code.

| **Optically-clear plate seals** | Seals the qPCR plate during the run |
|---|---|

Any kind of optically-clear sealing film should work, so substitutions for sealing film are fine. We use and like Applied Biosystems MicroAmp Optical Adhesive Film ref 4311971.

| **qPCR instrument** | Used to run the paDSF experiment |
|---|---|

Any qPCR instrument capable of running DSF can be used to run a paDSF dye screen. If possible, use an instrument capable of monitoring in more than one set of fluorescent wavelengths at a time. We use the Analytik Jena qTower 384G, and monitor in FAM, JOE, TAMRA, ROX, Cy5, and Cy5.5, but use of this instrument and precise channels is not necessary. Note that measuring in more wavelengths can, on some qPCR instruments, change increase the run-time, so testing the heating and measuring protocol prior to the experiment, and keeping the measured channels the same across experiments, can reduce unexpected changes to heating rate.

Like typical DSF, the heating procedure can be modified for the chosen protein. paDSF dyes do not have specific requirements for heating and measurement.

| **Optional: web-connected computer** | Used to analyze paDSF dye screen data |
|---|---|

paDSF dye screens can produce a large amount of data. You can use the provided web app to assist data analysis. Detailed information on hit calling is described in Supplementary Note 1.
